# Transfer and Capture of Envelope Protein Receptor-Binding Domains in the Retroviruses

**DOI:** 10.1101/2024.03.17.585361

**Authors:** Isidro Hötzel

## Abstract

The surface subunit (SU) of the envelope protein (Env) or retroviruses is highly variable due to adaptation to different hosts during their long evolutionary history. Several exogenous and endogenous retroviral gamma-like Env have a receptor-binding domain (RBD) in the amino-terminal region of SU that folds independently from the carboxyterminal SU C-domain. Two structurally distinct RBD classes have been described, one that adopts a modified immunoglobulin (Ig)-like domain fold and a second, in the Env of RDR interference group retroviruses, with a distinct β-sheet fold. Here, the distribution of different RBD classes among exogenous and endogenous gamma-type Env was determined by phylogenetic analyses of Env and structural modeling of retroviral SU with AlphaFold2. The patterns of RBD distribution indicate multiple RBD transfer events in the retroviruses. In addition, SU structural modeling identified an endogenous alpharetroviral-like Env in mammalian species with an amino-terminal RBD. This RDB has a typical IgV domain fold closely related both structurally and in sequence to the mammalian signal regulatory proteins (SIRP) α, β and γ, indicating, for the first time, acquisition of an RBD from a mammalian gene by the retroviruses. The results described herein indicate intragenic *env* recombination between retroviruses and between retroviruses and their hosts as a major factor in the evolution of retroviral Env.

## INTRODUCTION

The study of the evolution of retroviruses over geological timescales is enabled by their ability to infect and integrate into the germline of vertebrate species (1). Of the three canonical retroviral genes, the *pol* gene encoding the retroviral reverse-transcriptase that defines retroviruses is the most conserved and useful to understand long-range evolutionary relationships among retroviruses. The *env* gene encoding the envelope protein (Env) mediating cell infection is the most variable, related to the ability of retroviruses to adapt to a wide range of vertebrate species (2, 3). Despite its variation, Env still retains sufficient information about the evolution of retroviruses. The transmembrane subunit (TM) mediating membrane fusion during infection is the more conserved region of Env, whereas the surface subunit (SU) recognizing specific receptors on cells evolves more rapidly, especially in the receptor binding regions, with consequent loss of sequence similarity between more distantly related retroviruses (3). Endogenous *env* genes from germline-integrated retroviral proviruses encoding full-length or nearly full-length Env proteins are widely spread in vertebrate species. Some of these endogenous Env products, among these the syncytins, have been coopted by their hosts for key physiological mechanisms in reproduction and maintained in a functional state by purifying selection (3, 4).

Recombination has been a factor in both short and longterm evolutionary processes in the retroviruses (5). Recombination in the retroviruses can involve both lateral gene transfers among different retroviruses or between retroviruses and their hosts. In short or relatively short timescales, recombination of closely related retroviruses has played a key role in increasing viral genetic diversity and in the generation of pathogenic variants. The latter includes recombination events between exogenous retroviruses and closely related endogenous counterparts that lead to the generation of pathogenic variants with recombinant *env* genes (6–8). In longer time-scales, recombination between widely distinct retroviruses has led to the formation of new viral lineages harboring the *gag* and *pol* structural genes from one retroviral lineage and the *env* gene from another (9, 10). Finally, recombination has enabled the capture of cellular proto-oncogenes by the retroviruses, leading to retroviruses with oncogenes closely related to cellular genes in vertebrate species (11, 12). However, although general recombination and transfer of entire *env* genes between distinct retroviral lineages and intragene *env* recombination between closely related retroviruses and strains has been described, the possible role of intragene recombination in the evolution of retroviral Env over long time scales has not been documented.

The Env of retroviruses is structurally diverse. Most of this diversity is in SU, which may differ in length and retain little to no sequence similarity among diverse Env types. A major class of retroviral Env is the gamma-type Env, characterized by a CX_n_CC motif in TM (13). There are two main classes of gamma-type Env, the first one being the gamma-type proper, characterized by a CXXC motif in SU that forms an intersubunit disulfide with the TM CX_n_CC motif. This gamma-type Env is found mostly in the gammaretroviruses but it has also been transferred to other retroviral genera by recombination. The second class is the gamma-type Env of avian alpharetroviruses, hence named the avian gamma-type Env, which lacks a clear CXXC motif in SU and has instead a lone amino-terminal Cys residue forming the intersubunit disulfide bond. There are two main subclasses of gamma-type Env within extant retroviruses, the Type-C and the RDR-interference group (3, 10). Both have an SU segregated into two major domains, an amino-terminal receptor-binding domain (RBD) and a carboxy-terminal C-domain, with the boundary between both domains defined approximately by the CXXC motif in the linear sequence. In the Type-C gamma-type Env, the RBD is connected to the CXXC motif and C-domain by a proline-rich region (PRR). Although some similarity can be detected in the C-domain sequences of more distantly related retroviruses, the RBD regions are highly variable.

The only orthoretroviral SU regions that have been structurally characterized in detail are the RBD of Type-C retroviruses, the SU of the gamma-type syncytin-2 (Syn-2), encoded by the human endogenous retrovirus FRD (HERV-FRD) and the RBD of the endogenous HERV-Pb(1), or Env-Pb(1) (14–17). The RBD-C of murine (MLV) and feline (FeLV) leukemia viruses have very similar structures consisting of an L-shaped modified IgV set immunoglobulin (Ig)-like domain with three expanded loops, two of which, variable regions A and B (VRA and VRB), form the receptor-interacting regions. A similar structure is observed for the RBD of the endogenous Env-Pb(1), which has shorter VRA and VRB loops and a more canonical IgV-like domain. In contrast, Syn-2, which lacks an amino-terminal RBD, interacts with its receptor through SU loops located C-terminally from the CXXC motif (17).

The recent advent of structural modeling tools based on deep-learning artificial intelligence methods such as AlphaFold2 (AF2) has enabled characterization of Env proteins from a diverse set of retroviruses (18–23). An unexpected finding was that the SU of orthoretroviruses share a conserved and structurally unique β-sheet domain that interacts with TM, the proximal domain (PD), named after the corresponding domain of the SU of primate lentiviruses (21, 23). In the gamma-type Env of extant mammalian retroviruses, the PD corresponds to the C-domain, with the CXXC motif included in the first β-strand of the PD (23). The PD is not only found in the SU of gamma-type Env and lentiviruses but also in the SU-equivalent GP1 of the filoviral spike protein (23). The structural modeling results leading to the identification of the conserved PD structure in retroviral SU were confirmed with the subsequent description of the crystal structure of Syn-2 SU (17). Structural modeling of SU revealed that several gamma-type Env do not have an RBD and have instead an SU that is composed of a simple C-domain with an amino-terminal CXXC motif (23). The RBD of Type-C retroviruses is simply an amino-terminal extension of the PD that is not universally observed in gamma-type Env. Several gamma-type Env variants without and amino-terminal RBD have topologically distinct internal expansions of loops within the PD β -sheet that extend distally, including Syn-2 and the SU of avian leukemia virus (ALV). The PD of gamma-type Env, including the avian gamma-type Env, was recently shown by structural modeling of trimeric Env to be oriented in Env trimers in the same orientation as the PD domain of filoviral GP1 rather than lentiviral SU (22).

The Ig-like RBD of the Type-C retroviruses and HERV-Pb(1) and related endogenous elements (named here RBD-C for simplicity) is only one of the two major RBD types in the retroviruses. AF2 modeling recently showed that the RBD of RDR interference group retroviruses (named RBD-R here) is structurally distinct, composed of a single or duplicated three-strand β-sheet with defined strand topology and intradomain disulfide bond pattern (22, 23). Extant RDR group retroviruses have the duplicated RBD-R domain whereas the endogenous synctin-1 (Syn-1) and sycytin-Mar1 (Syn-Mar1) have a single-domain RBD-R. Another major distinction between RBD-C and RBD-R domains is their location within the Env trimer. Fitting of AF2 Env trimer models to previous electron-density maps of resting (not receptor-engaged) native trimeric MLV Env indicate that the RBD-C of MLV Env is located on the side of the Env trimer loosely associated with the rest of Env through the flexible PRR linker, pointing towards the viral membrane. In contrast, in the Env of an endogenous RDR retrovirus the RBD-R is located at the top of the Env trimer closely associated with SU, joined to the C-domain by a welldefined, mostly α -helical linker (22).

Structural modeling of retroviral Env and SU with AF2 has enabled the identification of the conserved PD and its differential expansions across several different retroviruses, and two major structural RBD types. Thus, AF2 modeling of retroviral Env proteins has extended the understanding of the structural evolution of monomeric and trimeric retroviral Env beyond what sequence analyses are able to reveal. Here a large-scale survey of retroviral SU by AF2 structural modeling was used in conjunction with phylogenetic analyses to understand the evolution of RBD domains in the gamma-type Env of retroviruses. The results indicate intragenic *env* recombination as a major factor driving the evolution retroviral Env, with transfer of RBD regions within and between major retroviral groups and acquisition of RBD-like regions from host genes.

## RESULTS

### Structural definition of RBD classes in the main retroviral Env group

Retroviral gamma-type Env sequences from exogenous extant retroviruses and endogenous elements were retrieved from the non-redundant protein sequence section of GenBank by BLAST searches using previously described gamma-type Env sequences as query. Additional previously described consensus endogenous Env sequences were added for phylogenetic analyses and clustering of Env sequences. All sequences had apparent secretion signal sequences with most including clear signal sequence cleavage sites (Suppl. File 1). A maximum likelihood phylogenetic tree of the TM ectodomains of these Env sequences showed three major clusters with good support values: An alpharetroviral-like gamma-type Env group without a CXXC motif in SU, a major group including the Env sequences of extant retroviruses from different retroviral genera and related endogenous elements including syncytin-1 (Syn-1) encoded by HERV-W, Syn-2, syncytin-A and B, and the HEMO and ENVV2 elements, and a separate major Env cluster related to syncytin-Mar1 and EnvR (Fig. 1). Table 1 describes the origin and nomenclature of Env sequences used here, with additional information and complete sequences included in Supplementary Table 1. The three major Env groups are referred here as the Alpha-like and Main Env Groups and the EnvR Env Supergroup respectively. Of note, endogenous elements are referred to as retroviruses for simplicity. The Main and EnvR groups include gamma-type Env sequences with CXXC motifs in SU. Support values for branching of sequences within each major group were moderate to low except in the alpharetrovirus-like group, although subgroups clustered in a manner similar to previously described phylogenetic analyses of TM ectodomain sequences (16, 24–28).

**Figure 1.**
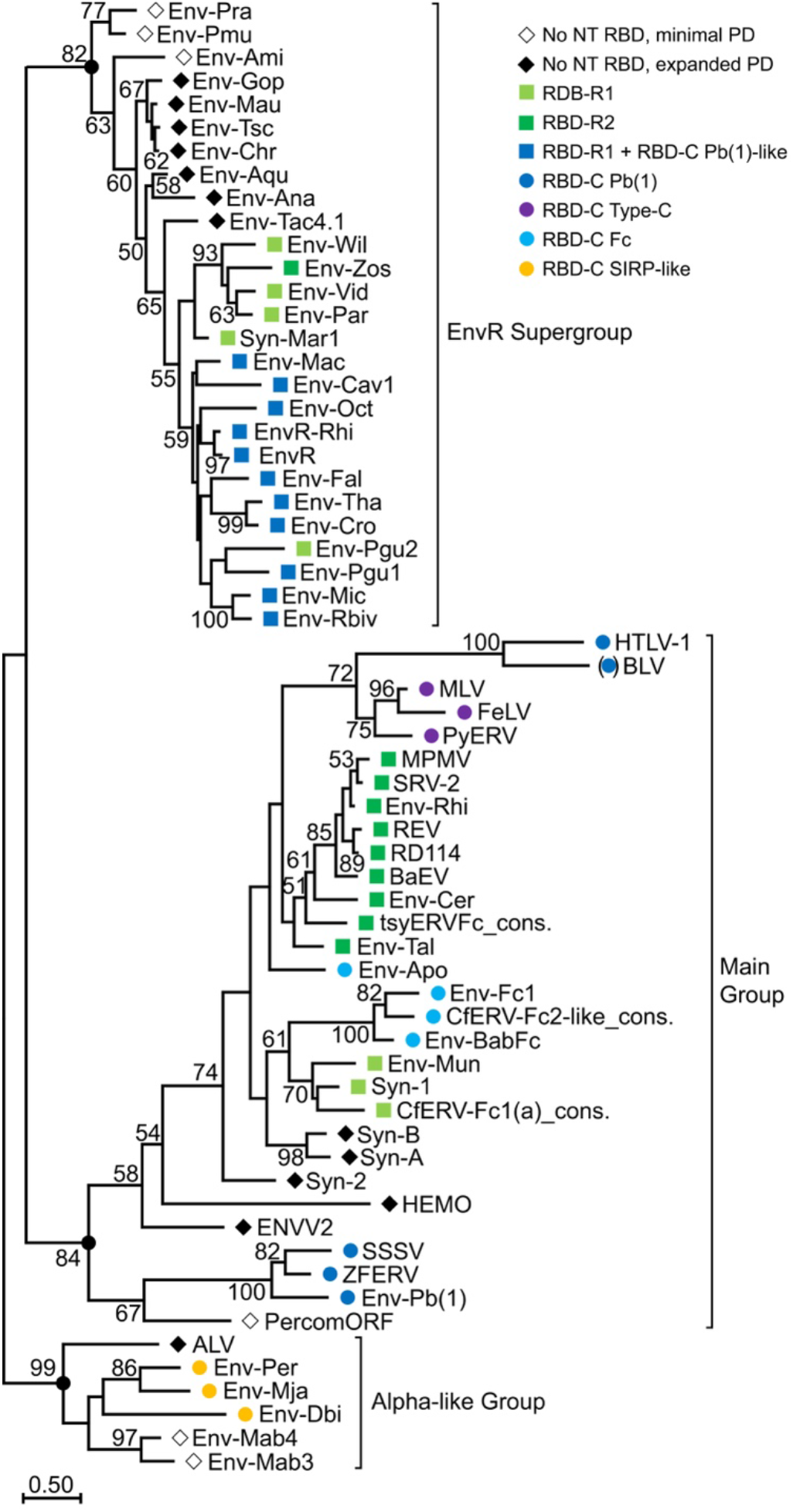
Maximum likelihood phylogenetic tree of gamma-type TM ectodomains. The tree is rooted on the Alpha-like group for clarity. The three main gamma-type Env groups are indicated on the right. Nodes with bootstrap values (100 repeats) greater than 50 are indicated. Black circles indicate the nodes defining each of the three major Env groups. Symbols next to Env names indicate the presence or absence of an amino-terminal RBD and RBD type for each Env. Parentheses in BLV indicate presumed RBD type. The tree is drawn to scale with the distance scale indicating amino-acid substitutions per site.

**Table 1.**
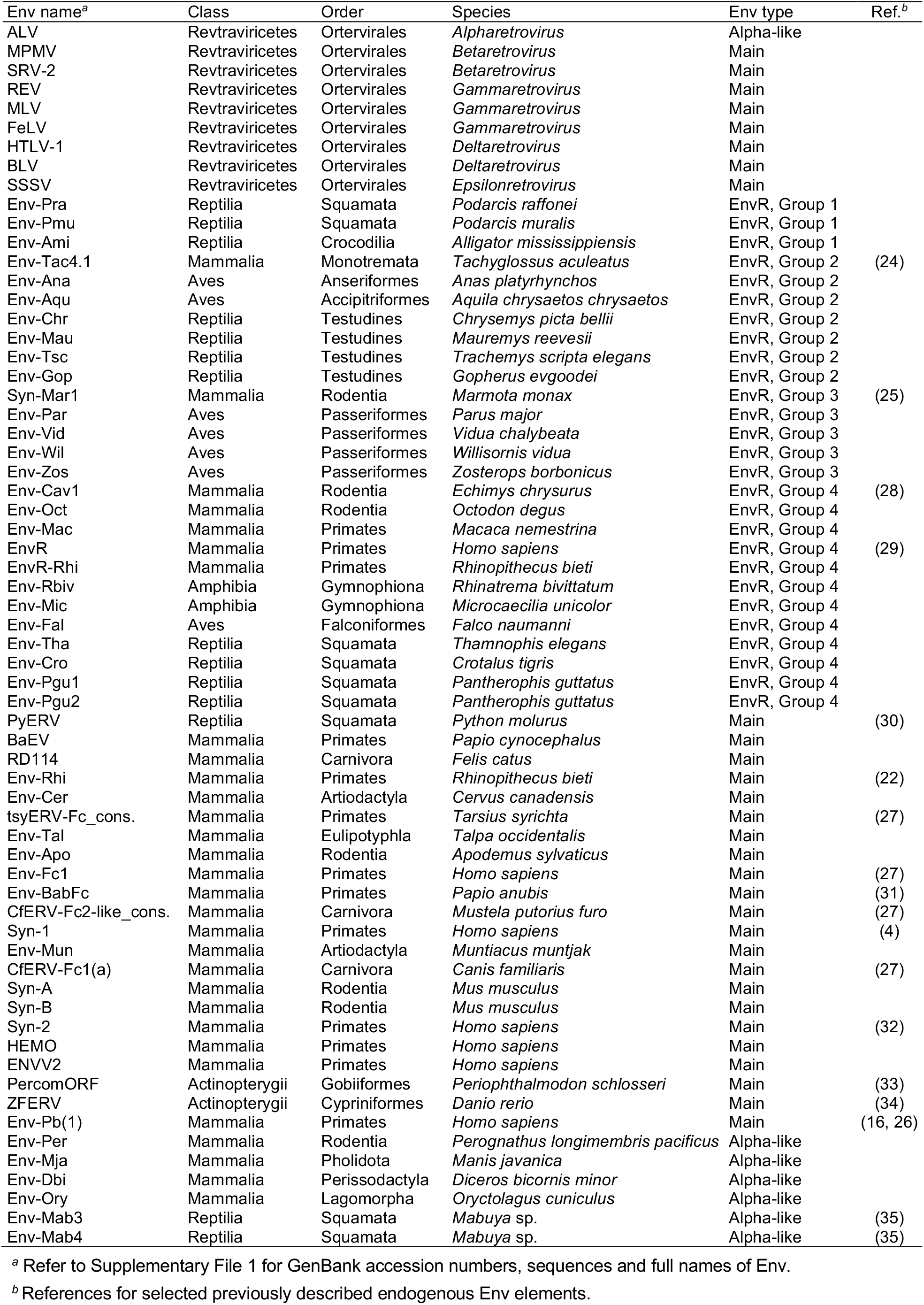
Exogenous and endogenous Env names, taxonomic origin and grouping.

| Env name<sup>a</sup> | Class | Order | Species | Env type | Ref.<sup>b</sup> |
| --- | --- | --- | --- | --- | --- |
| ALV | Revtraviricetes | Ortervirales | *Alpharetrovirus* | Alpha-like |  |
| MPMV | Revtraviricetes | Ortervirales | *Betaretrovirus* | Main |  |
| SRV-2 | Revtraviricetes | Ortervirales | *Betaretrovirus* | Main |  |
| REV | Revtraviricetes | Ortervirales | *Gammaretrovirus* | Main |  |
| MLV | Revtraviricetes | Ortervirales | *Gammaretrovirus* | Main |  |
| FeLV | Revtraviricetes | Ortervirales | *Gammaretrovirus* | Main |  |
| HTLV-1 | Revtraviricetes | Ortervirales | *Deltaretrovirus* | Main |  |
| BLV | Revtraviricetes | Ortervirales | *Deltaretrovirus* | Main |  |
| SSSV | Revtraviricetes | Ortervirales | *Epsilonretrovirus* | Main |  |
| Env-Pra | Reptilia | Squamata | *Podarcis raffonei* | EnvR, Group 1 |  |
| Env-Pmu | Reptilia | Squamata | *Podarcis muralis* | EnvR, Group 1 |  |
| Env-Ami | Reptilia | Crocodylia | *Alligator mississippiensis* | EnvR, Group 1 |  |
| Env-Tac4.1 | Mammalia | Monotremata | *Tachyglossus aculeatus* | EnvR, Group 2 | (24) |
| Env-Ana | Aves | Anseriformes | *Anas platyrhynchos* | EnvR, Group 2 |  |
| Env-Aqu | Aves | Accipitriformes | *Aquila chrysaetos chrysaetos* | EnvR, Group 2 |  |
| Env-Chr | Reptilia | Testudines | *Chrysemys picta bellii* | EnvR, Group 2 |  |
| Env-Mau | Reptilia | Testudines | *Mauremys reevesii* | EnvR, Group 2 |  |
| Env-Tsc | Reptilia | Testudines | *Trachemys scripta elegans* | EnvR, Group 2 |  |
| Env-Gop | Reptilia | Testudines | *Gopherus evgoodei* | EnvR, Group 2 |  |
| Syn-Mar1 | Mammalia | Rodentia | *Marmota monax* | EnvR, Group 3 | (25) |
| Env-Par | Aves | Passeriformes | *Parus major* | EnvR, Group 3 |  |
| Env-Vid | Aves | Passeriformes | *Vidua chalybeata* | EnvR, Group 3 |  |
| Env-Wil | Aves | Passeriformes | *Willisornis vidua* | EnvR, Group 3 |  |
| Env-Zos | Aves | Passeriformes | *Zosterops borbonicus* | EnvR, Group 3 |  |
| Env-Cav1 | Mammalia | Rodentia | *Echimys chrysurus* | EnvR, Group 4 | (28) |
| Env-Oct | Mammalia | Rodentia | *Octodon degus* | EnvR, Group 4 |  |
| Env-Mac | Mammalia | Primates | *Macaca nemestrina* | EnvR, Group 4 |  |
| EnvR | Mammalia | Primates | *Homo sapiens* | EnvR, Group 4 | (29) |
| EnvR-Rhi | Mammalia | Primates | *Rhinopithecus bieti* | EnvR, Group 4 |  |
| Env-Rbiv | Amphibia | Gymnophiona | *Rhinatrema bivittatum* | EnvR, Group 4 |  |
| Env-Mic | Amphibia | Gymnophiona | *Microcaecilia unicolor* | EnvR, Group 4 |  |
| Env-Fal | Aves | Falconiformes | *Falco naumanni* | EnvR, Group 4 |  |
| Env-Tha | Reptilia | Squamata | *Thamnophis elegans* | EnvR, Group 4 |  |
| Env-Cro | Reptilia | Squamata | *Crotalus tigris* | EnvR, Group 4 |  |
| Env-Pgu1 | Reptilia | Squamata | *Pantherophis guttatus* | EnvR, Group 4 |  |
| Env-Pgu2 | Reptilia | Squamata | *Pantherophis guttatus* | EnvR, Group 4 |  |
| PyERV | Reptilia | Squamata | *Python molurus* | Main | (30) |
| BaEV | Mammalia | Primates | *Papio cynocephalus* | Main |  |
| RD114 | Mammalia | Carnivora | *Felis catus* | Main |  |
| Env-Rhi | Mammalia | Primates | *Rhinopithecus bieti* | Main | (22) |
| Env-Cer | Mammalia | Artiodactyla | *Cervus canadensis* | Main |  |
| tsyERV-Fc_cons. | Mammalia | Primates | *Tarsius syrichta* | Main | (27) |
| Env-Tal | Mammalia | Eulipotyphla | *Talpa occidentalis* | Main |  |
| Env-Apo | Mammalia | Rodentia | *Apodemus sylvaticus* | Main |  |
| Env-Fc1 | Mammalia | Primates | *Homo sapiens* | Main | (27) |
| Env-BabFc | Mammalia | Primates | *Papio anubis* | Main | (31) |
| CfERV-Fc2-like_cons. | Mammalia | Carnivora | *Mustela putorius furo* | Main | (27) |
| Syn-1 | Mammalia | Primates | *Homo sapiens* | Main | (4) |
| Env-Mun | Mammalia | Artiodactyla | *Muntiacus muntjak* | Main |  |
| CfERV-Fc1(a) | Mammalia | Carnivora | *Canis familiaris* | Main | (27) |
| Syn-A | Mammalia | Rodentia | *Mus musculus* | Main |  |
| Syn-B | Mammalia | Rodentia | *Mus musculus* | Main |  |
| Syn-2 | Mammalia | Primates | *Homo sapiens* | Main | (32) |
| HEMO | Mammalia | Primates | *Homo sapiens* | Main |  |
| ENVV2 | Mammalia | Primates | *Homo sapiens* | Main |  |
| PercomORF | Actinopterygii | Gobiiformes | *Periophthalmodon schlosseri* | Main | (33) |
| ZFERV | Actinopterygii | Cypriniformes | *Danio rerio* | Main | (34) |
| Env-Pb(1) | Mammalia | Primates | *Homo sapiens* | Main | (16, 26) |
| Env-Per | Mammalia | Rodentia | *Perognathus longimembris pacificus* | Alpha-like |  |
| Env-Mja | Mammalia | Pholidota | *Manis javanica* | Alpha-like |  |
| Env-Dbi | Mammalia | Perissodactyla | *Diceros bicornis minor* | Alpha-like |  |
| Env-Ory | Mammalia | Lagomorpha | *Oryctolagus cuniculus* | Alpha-like |  |
| Env-Mab3 | Reptilia | Squamata | *Mabuya* sp. | Alpha-like | (35) |
| Env-Mab4 | Reptilia | Squamata | *Mabuya* sp. | Alpha-like | (35) |

<sup>a</sup> Refer to Supplementary File 1 for GenBank accession numbers, sequences and full names of Env.<sup>b</sup> References for selected previously described endogenous Env elements.

A survey of RBD-C structures in the Main Env group was done by *de novo* structural modeling of SU with the ColabFold implementation of AF2, without templates to avoid biases in structural modeling. The SU of HEMO and ENVV2 were not successfully modeled. However, these Env variants have CXXC motifs close to the amino-terminus of SU, similar to Syn-2 (32, 36, 37), indicating that these do not have independently-folding amino-terminal RBDs and that the SU of these elements are topologically more similar to Syn-2, Syn-A and Syn-B, given the distance between the CXXC motif and the Env cleavage site. The structure of the SU of other Main group Env members for which there were no previously described structures were successfully modeled, with high confidence scores for most members. All models have C-domains with a typical minimal PD structure, as previously described for the SU of extant Type-C and RDR group retroviruses (23), without any major expansions of the PD region or other notable structural features (not shown).

The RBD-C of MLV, FeLV and Env-Pb(1), while structurally similar, are differentiated by the length of the VRA and VRB regions, longer in the RBD-C of Type-C retroviruses and shorter in the RBD-C of Env-Pb(1) (16). In addition, the RBD-C of Env-Pb(1) lacks an additional amino-terminal β - strand observed in the RBD-C of MLV and FeLV, giving it a more canonical IgV-like fold, lacking only the disulfide between strands 2β and 9β observed in most IgV domains (15, 38) (following the naming system originally used to describe the MLV RBD structure; β -strands B and F in the IgV domain naming convention) (Fig. 2A and B). These two previously described RBD-C structural types are referred to here as RBD-C Type-C and RBD-C Pb(1). As expected, the high-quality RBD structural model for the SU of ZFERV, which clusters with Env-Pb(1) in TM phylogenetic trees, is more similar to the RBD-C Pb(1) type than the RBD-C Type-C structure except for a C-terminal extension introducing an additional β -strand in the domain (Fig. 2C, Suppl. Fig. 1C).

**Figure 2.**
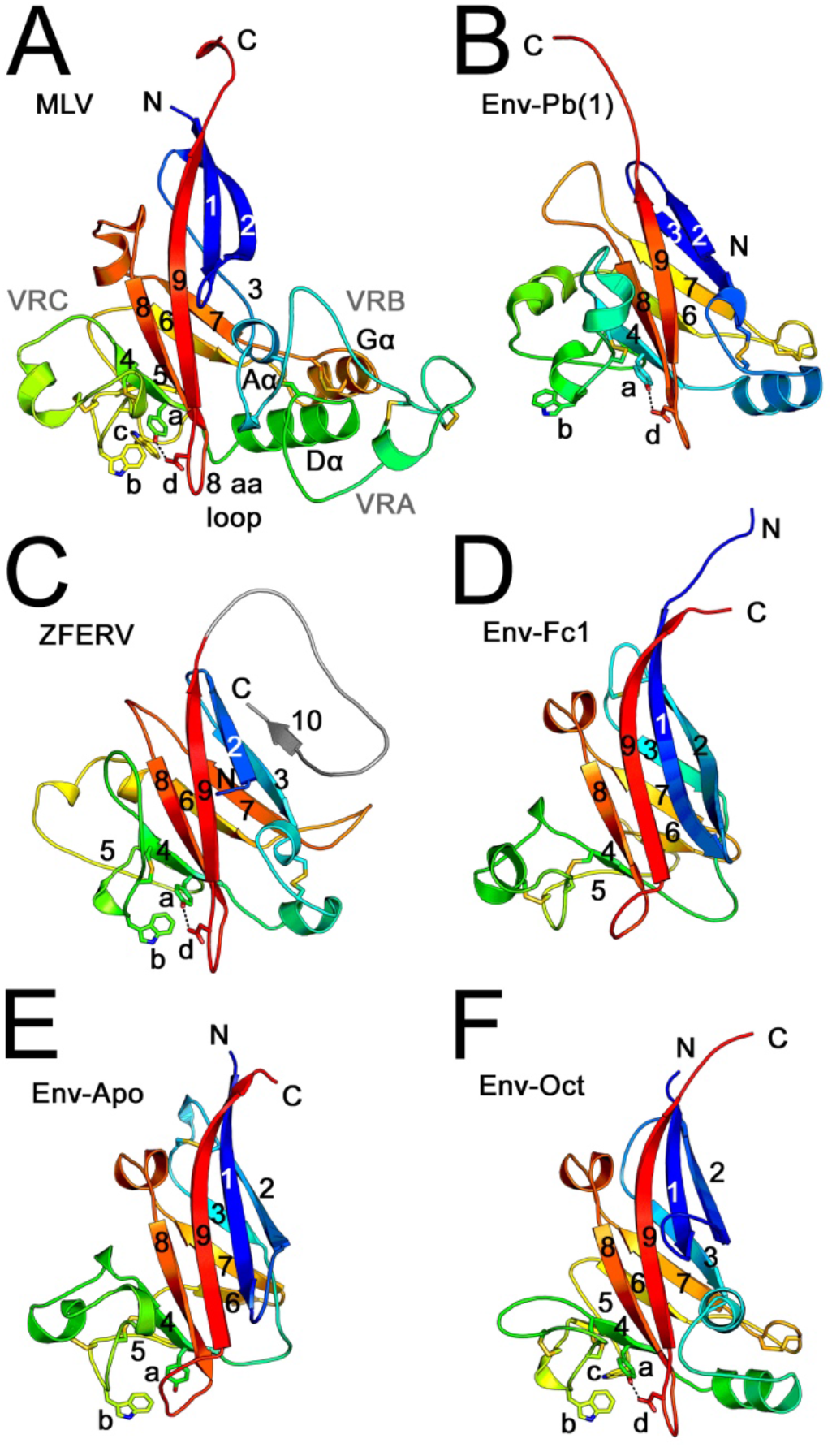
RBD-C structures and models. The crystal structures of the RBD regions of MLV Env (PDB 1AOL) (A) and Env-Pb(1) (PDB 6W5Y, chain A) (B) and AF2 structural models for the RBD-C regions of indicated Env proteins (C-F) are shown in rainbow coloring from blue to red from the amino to the carboxy-terminal ends of the domains. The β-strands are numbered according to the MLV crystal structure, with the VRA, VRB and VRC regions, major helical regions and the structurally conserved 8 aa-loop indicated in panel A with the MLV RBD structure. Disulfides are shown as sticks. Side-chains of the conserved tetrad nearby the VRC region are shown as sticks, labeled a-d, with the hydrogen-bond between residues a and d shown by a dotted line. The carboxy-terminal ZFERV RBD extension with an additional β -strand is colored gray.

The Env in the Env-Fc1/Fc2 cluster within the Main Env group is expected to have an amino-terminal RBD given the location of the CXXC motif in a central location of SU. Two different Fc1/Fc2 types have been described, a major group that clusters separately from Syn-1 and a subset that clusters with Syn-1 (27). Modeling of the putative RBD region of the former group, which includes Env-BabFc (BabFc^*env*^), resulted in high-quality model structures with an IgV-like RBD-C domain resembling that of RBD-C Pb(1) but without discernable VRA and VRB regions (Fig. 2D, Suppl. Fig. 1D). The SU structural model for Env-Apo, which appears to cluster more closely with the RDR Env group by TM sequence, also has an RBD-C similar to RBD-C of Env-Fc1 (Fig. 2E, Suppl. Fig. 1E). This defines a third RBD-C type, named here RBD-C Fc. Minimal sequence similarity is observed between the RBD-C Type-C, Pb(1) and Fc1 classes, limited to subsets of cysteine residues involved in disulfide bonds within the IgV-like domain fold or scattered residues, clear only in structure-aided sequence alignments (Fig. 3).

**Figure 3.**
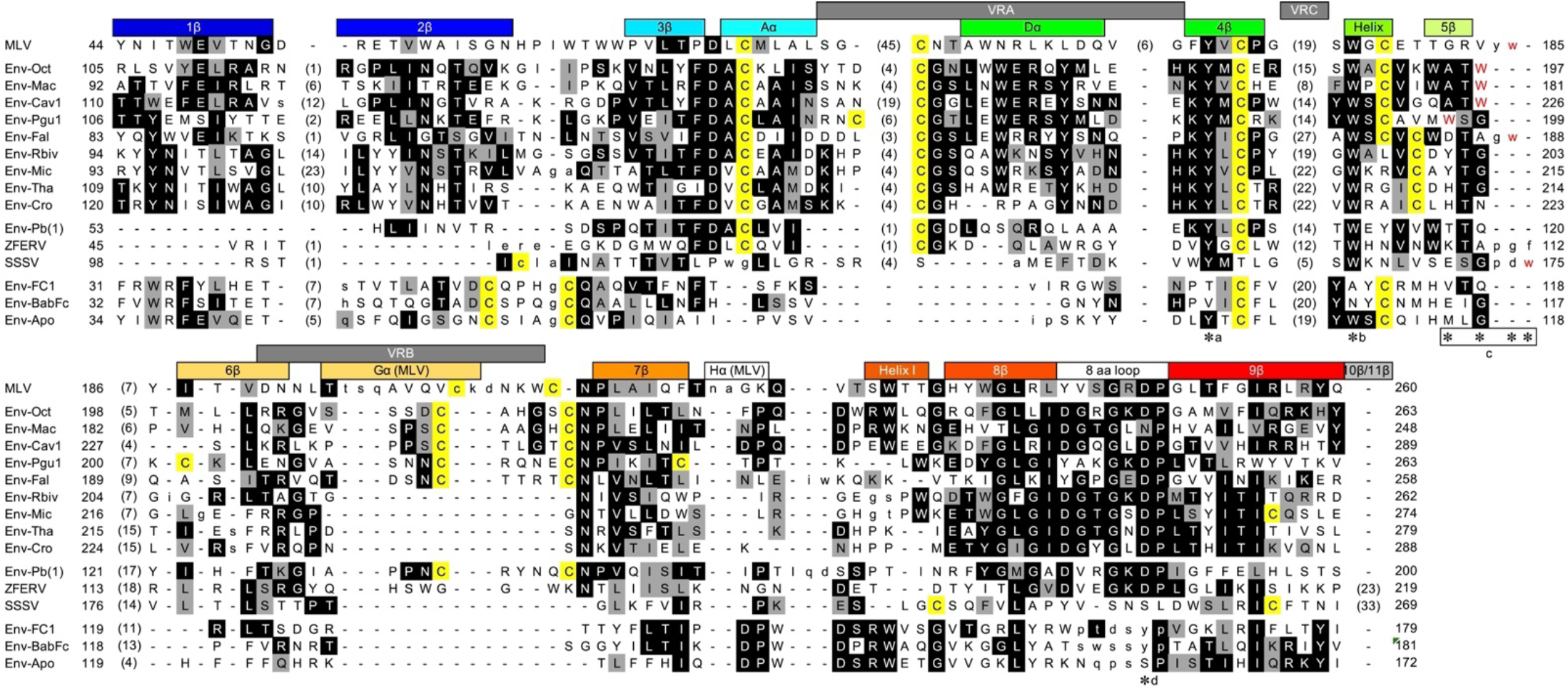
Alignment of RBD-C sequences. RBD-C structures and models were aligned structurally to generate a multiple sequence alignment using the sequence of Env-Oct as reference. Non-conserved loops are omitted from the alignment for clarity, with the number of residues not shown indicated in parentheses. Black shading indicates identity with the consensus residue. Gray shading indicates residues similar to the consensus according to the BLOSUM-62 similarity matrix. Cysteine residues are highlighted in yellow. Residues in lower case have alpha-carbons that are not aligned to the Env-Oct RBD-C model. Dashes indicate gaps. The residues forming the different β-strands and helical regions of the domain are indicated above the alignment, with coloring similar to that shown in Figure 2. The VRA, VRB and VRC regions of the MLV RBD are indicated in gray above the sequence. Residue numbering from the start of the Env sequence. Asterisks indicate the conserved tetrad residues, labeled by a-d letters as in Figure 2. The Trp residues of the tetrad (residue c) in different sequences are highlighted in red, with the box indicating the range of positions for that residue in the alignment. The sequences are clustered according to RBD-C structural type.

The only other Env subgroup within the Main Env group in Figure 1 expected to have an RBD-C is that of the deltaretroviruses HTLV-1, HTLV-2 and BLV. AF2 modeling of the SU of these viruses was not generally successful, one exception being the RBD region of a single HTLV-1 sequence (Suppl. Fig. 2). The HTLV-1 RBD structural model is similar to that of other RBD-C structures and models. The quality scores were high for the entire model except the region structurally homologous to the VRC region, which was mostly unfolded in the AF2 model. However, the regions of high reliability scores in the HTLV-1 RBD model indicate that is has an intermediate structure between those of the Fc and Pb(1) types, including a short helical region for VRA as in Env-Pb(1) but no long loop region corresponding to the VRB region after β-strand 6, as in the RBD-C Fc (Suppl. Fig. 2). The structural modeling results indicate three structurally distinct RBD-C types in retroviral Env, RBD-C Type-C, Pb(1) and Fc, with the RBD-C of deltaretroviruses sharing similarities with the latter two types.

The structural models for the SU of Env sequences clustering with the Env of RDR interference group retroviruses had the typical RBD-R structures previously described for this retroviral group (23). These fall into two different RBD-R subclasses, referred here as RBD-R1 and RBD-R2 (Fig. 1). The RBD-R1 type, observed in Syn-1 and closely-related endogenous elements, has a single β-sheet whereas the RBD-R2 type in extant RDR interference group retroviruses and related endogenous elements has a structurally repeated RBD with two topologically similar β-sheets as previously described (22, 23). No additional RBD-R structural types beyond these two major classes were identified within the set of Main Env group sequences analyzed.

Although bootstrap support values for individual groups are relatively low in TM phylogenetic tree within the Main Env group, the distribution of different RBD-C and RBD-R types, with the possible exception of the epsilonretroviruses and deltaretroviruses, agrees with the clustering of the corresponding TM ectodomain sequences (Fig. 1). Interestingly, more basal Env sequences (by TM sequence alignment) within the Main group have no amino-terminal RBD regions before the CXXC motif, suggesting that independently-folding RBDs evolved relatively late event in this Env group.

### Diversity of PD structural types in the EnvR Supergroup

The Env of the human endogenous retrovirus ERV3-1, EnvR, has an atypical CX_7_CC motif in TM rather than the CX_6_CC motif of the Env of extant gammaretroviruses and alpharetroviruses. Likewise, all Env sequences in the EnvR group cluster have a CX_7-9_CC motif in TM. This group is composed of endogenous Env sequences, including the previously described endogenous Env sequences Syn-Mar1, Env-Cav1 and Env-Tac4 (Fig. 1, Table 1). Env sequences in the EnvR supergroup are derived from species in all tetrapod classes (Table 1). This group is distinct from all other gamma-type Env types, forming a well-supported group in the TM phylogenetic tree (Fig. 1). As for the TM sequences in the Main group, support values for subgroups within this supergroup are variable.

Analysis of SU sequences within the EnvR supergroup revealed 4 major types of domain organization (Fig. 4). Group 1 including sequences of lizards in the *Podarcis* genus had what appears to be an SU comprised of a nearly minimal PD, with the CXXC motif about 40 residues from the imputed amino-terminus of mature Env after the signal sequence cleavage site and about 175 residues from the SU C-terminus. For reference, the smallest gamma-type SU comprised of a minimal PD previously reported has an CXXC motif 25 residues from the Env N-terminus and a distance of 162 residues between this motif and the SU C-terminus (23). Therefore, relatively short internal expansions of the PD and no amino-terminal RBD are expected in the Group 1 SU sequences. This is the only group with a CX_9_CC motif in TM, in contrast with the CX_7_CC motif of other members of the EnvR supergroup. Group 2 includes the previously described endogenous Env-Tac4.1 Env of monotremes and has an SU with an CXXC motif between 21 and 35 residues from the mature Env N-terminus and about 300 residues from the SU C-terminus, indicating significant internal expansion of PD regions but no amino-terminal RBD. Group 3, including Syn-Mar1 and avian endogenous elements, has the CXXC motif about 130 residues from the mature Env N-terminus and a distance of about 220 residues between the CXXC motif and the SU C-terminus, suggesting an amino-terminal RBD-like region and relatively minimal, if any, internal PD expansions. In fact, in Syn-Mar1 this amino-terminal region was previously shown to be comprised of an RBD-R domain and linker region (23). The last group, which includes EnvR, has the CXXC motif about 260 residues from the mature Env N-terminus and 175 residues from the SU C-terminus, indicating a larger amino-terminal RBD-like region than the group 3 sequences and a relatively minimal internal PD expansion.

**Figure 4.**
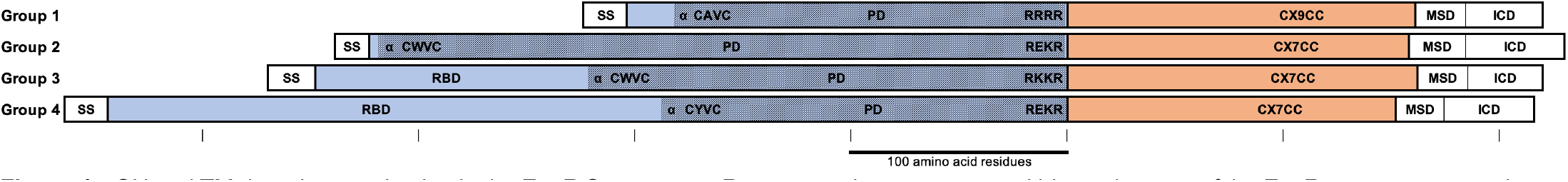
SU and TM domain organization in the EnvR Supergroup. Representative sequences within each group of the EnvR supergroup are shown, with the SU and TM ectodomain sequences in blue and salmon. The shaded region in SU indicates the presumed PD region defined by the location of the CXXC motif and Env cleavage site. RBD indicates the presumed sequences of RBD regions that fold independently in SU in analogy with the SU of gammaretroviruses and as previously shown for Syn-Mar1, in group 3, by structural modeling (23). The number of residues in the TM CX_n_CC motif is shown, with the membrane-spanning (MSD) and intracellular regions (ICD) of TM shown in white boxes. The white boxes in the amino-terminal end of Env labeled as ss indicate predicted signal sequences. Ticks and bar indicate 100 amino acid residues, with the Env cleavage site as origin for shown distances. The approximate location of αPD1 in the models defining the amino-terminal end of the C-domain (22) is indicate with an “α”.

Although most endogenous Env structural models in this group are available in the AlphaFold Structure Database, these include the entire, uncleaved Env which often have the TM region in an unfolded conformation. Thus, to determine if the different SU sequence organization types reflect significantly different structural types, the structure of the SU of representatives of each group from different tetrapod classes were modeled with AF2. Most sequences yielded models of sufficient local pLDDT quality scores for further analyses (Suppl. Fig. 3).

Visual inspection of the PD region of the SU models revealed that the 4 major EnvR groups with different SU domain organizations have different PD expansions, each set including an independent and complete tertiary fold module with disulfide bonds formed mostly within the modules. The expansion regions described here reflect independent structural substructures and the likely sequence of events leading to the different PD types within the EnvR group, described below.

The EnvR group 1 and one member of group 3, Env-Ami, have a PD with a relatively minimal internal PD expansion between strands β6 and β7 in the apical region (23), named here Expansion 1 (Fig. 5A and E, Suppl. Fig. 4). This expansion introduces a β-strand antiparallel to strand β6 in the apical region of the PD.

**Figure 5.**
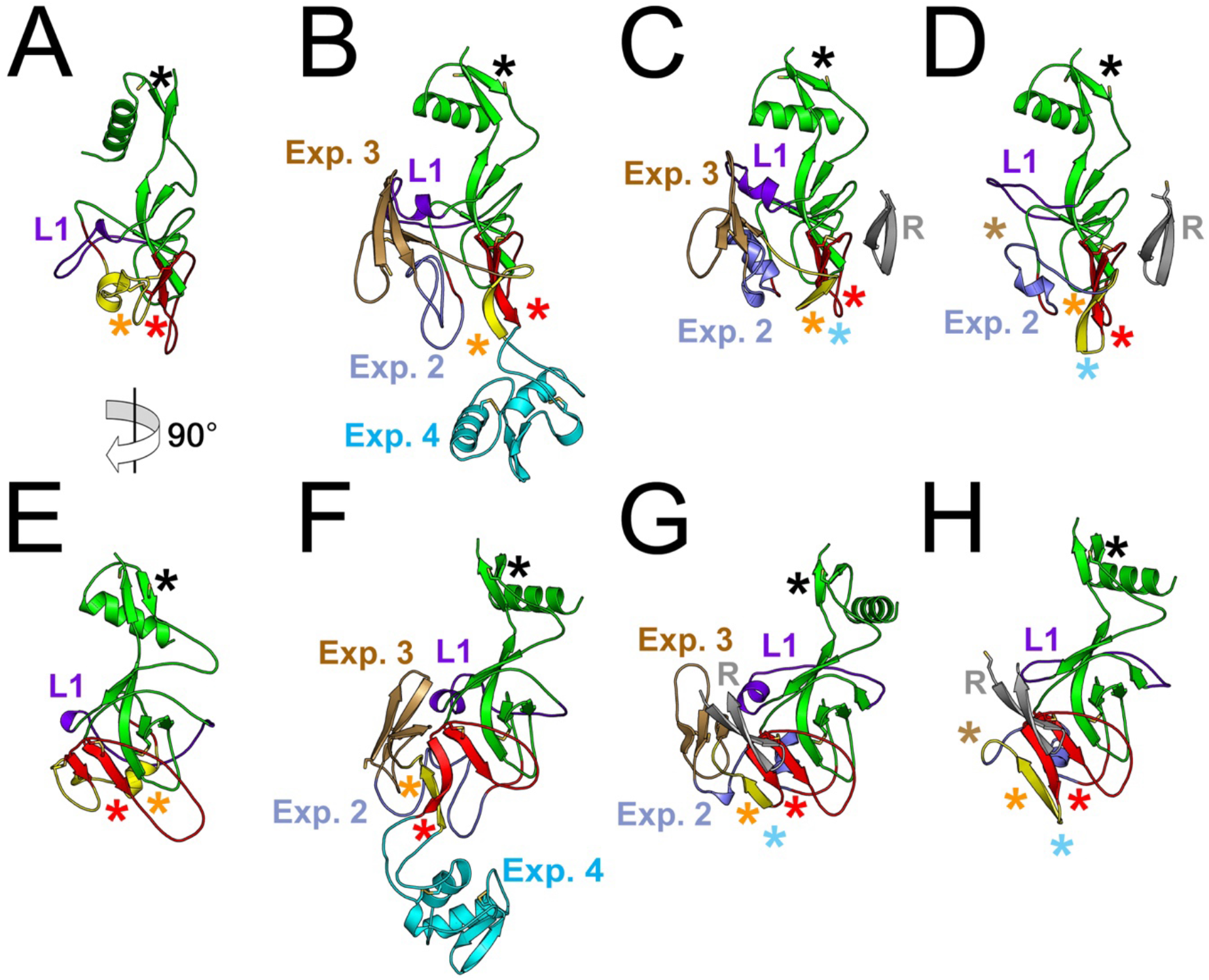
Structural models of the PD regions of SU of different groups in the EnvR supergroup. The structural models of the PD of EnvR groups 1 (A and E), 2 (B and F), 3 (C and G) and 4 (D and H) are shown with Expansion regions 1 to 4 colored in yellow (highlighted with orange asterisks), cyan, brown and light blue, respectively, and indicated by legends. PD β-strands 5 and 6 in the apical region are colored red and highlighted with a red asterisk. The “layer 1” region of the PD (23) is colored in dark blue and the PD core structure is colored green. Panels E-H are rotated 90° relative to panels A-D as indicated in panel A. Black asterisks indicate the location of the CXXC motif in the models. The brown and cyan asterisks indicate the locations from which Expansion regions 3 and 4 emanate in other models. The gray β-strands that are not connected to the PD show the RBD-R1 β-hairpins that contact the PD apical region in EnvR groups 3 and 4 in panels C, D, G and H. Models shown are those for Env-Pra (A and E), Env-Ana (B and F), Syn-Mar1 (C and G) and EnvR (D and H).

The EnvR group 2 has the most complex set of internal PD expansions, with 3 additional discrete subregions of secondary and tertiary structure in addition to Expansion 1 (Fig. 5B and F, Suppl. Fig. 4). Expansion 4 is located between strand β6 of the PD and Expansion 1 (Fig. 5B and F, Suppl. Fig. 4, cyan regions). Expansion 3, lies immediately after Expansion 1 and is composed of a β-sheet with 3 antiparallel β-strands held together by a disulfide bond, with the third β-strand between the first and second β-strands (Fig. 5B and F, Suppl. Fig. 4, brown regions). Expansion 2 follows Expansion 3 and consists of an extended region before the polypeptide chain re-enters the core region of the PD (Fig. 5B and F, Suppl. Fig. 5, light blue regions). One EnvR group 2 Env, Env-Aqu, has Expansion region 5, which consists of a β-hairpin held in place by a disulfide bond and an α-helix (Suppl. Fig. 4, black region). All other EnvR group 2 Env have Expansion regions 1 to 4 only. The PD expansion regions in the EnvR group 2 have the same relative position in all models except Expansion 4, which has a variable position among models that probably reflects structural flexibility (Suppl. Fig. 5A).

EnvR groups 3 and 4 have progressive deletions relative to the PD expansions of group 2. The SU of EnvR group 3 does not include Expansion 4 whereas group 4 SU does not include Expansions 3 and 4 (Fig. 5C, D, G and H, Suppl. Fig. 4). These modeling results identify several discrete PD expansion modules in the EnvR supergroup with secondary and tertiary structures in the region between β-strands 6 and of the PD. Within each group the structures of the expansions are essentially the same (Suppl. Fig. 5).

### Structure of the RBD-R of the EnvR Supergroup

EnvR groups 1 and 2 have no observable RBD in the amino-terminal region of SU structural models, consistent with the location of the CXXC motif in the amino-terminal region of SU in these groups. SU models of Groups 3, which includes syncytin-Mar1, and 4 have a single-repeat RBD-R1 domain in the SU amino-terminus analogous to that of human Syn-1 and similar to the previously described Syn-Mar1 SU model. The RBD-R1 has two additional β-strands that form a β-hairpin outside the main β-sheet (Fig. 6A and B, Suppl. Fig. 6A and B). The five β-strands in this domain are referred to here as strands β1R to β5R, with strands β1R and β2R forming the amino-terminal hairpin loop and stands β3R to β5R forming the three-strand β-sheet (Fig. 6A and B, Suppl. Fig. 6A and B). Local pLDDT reliability scores were high for the RBD-R1 models of groups 3 and 4 (Suppl. Fig. 7A and B).

**Figure 6.**
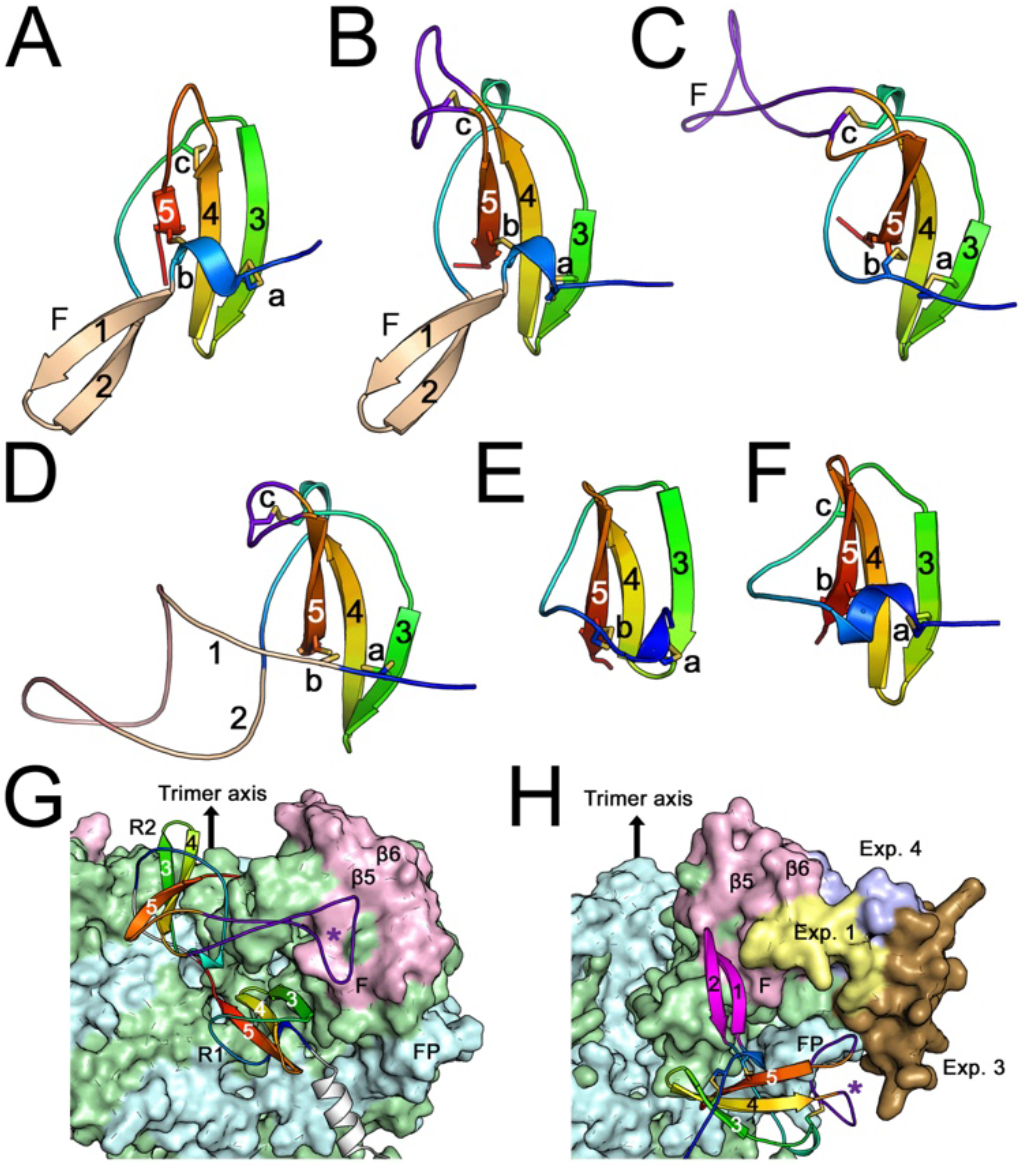
Structural models of the RBD-R regions of the SU of the EnvR supergroup, RDR Env and Syn-1. (A) RBD-R1 of group 4 EnvR shown in rainbow coloring from the amino-terminus of the domain in blue to the carboxy-terminus in red except for the β-hairpin with strands β1R and β2R shown in wheat. Strands are numbered 1 to 5 with disulfide bonds a to c shown as sticks and labeled. F indicates the location of apical loop F of the PD, not shown. (B) The same as panel A for the RBD-R1 of group 3 Syn-Mar1, with the extended loop between strands β4R and β5R shown in purple. (C) Second domain of the RBD-R2 of RDR interference group Env-Rhi labeled as in panel B. (D) RBD-R1 of Syn-1. (E) First domain of the Env-Rhi RBD-R2. (F) First domain of Env-Zos RBD-R2. Models in panels A-F are shown in the same orientation for clarity. (G) Position of the RBD-R2 or Env-Rhi in the previously described trimeric Env model (ModelArchive accession max4xpz). The distal end of the trimer points up as indicated. R1 and R2 indicate the first and second domains of the RBD-R2, each colored in a rainbow pattern as in panels C and E, with the asterisk and β5 and β6 showing the approximate location of the apical PD structures contacted by the β4R/β5R loop of the second RBD-R2 domain. FP, fusion peptide. (H) Expected location of the RBD-R1 of Syn-Mar1 in trimeric Env. The TM regions of the trimer are those of the model of Env-Rhi in panel G, with the SU Syn-Mar1 positioned by alignment of the core regions of the PD with the SU of Env-Rhi. The β1R/β2R β-hairpin is shown in magenta for clarity. Exp. 1, 3 and 4 indicate the locations of PD expansions, colored as in Figure 5C. The purple asterisk indicates the position of the β4R/β5R loop covering the fusion peptide.

**Figure 7.**
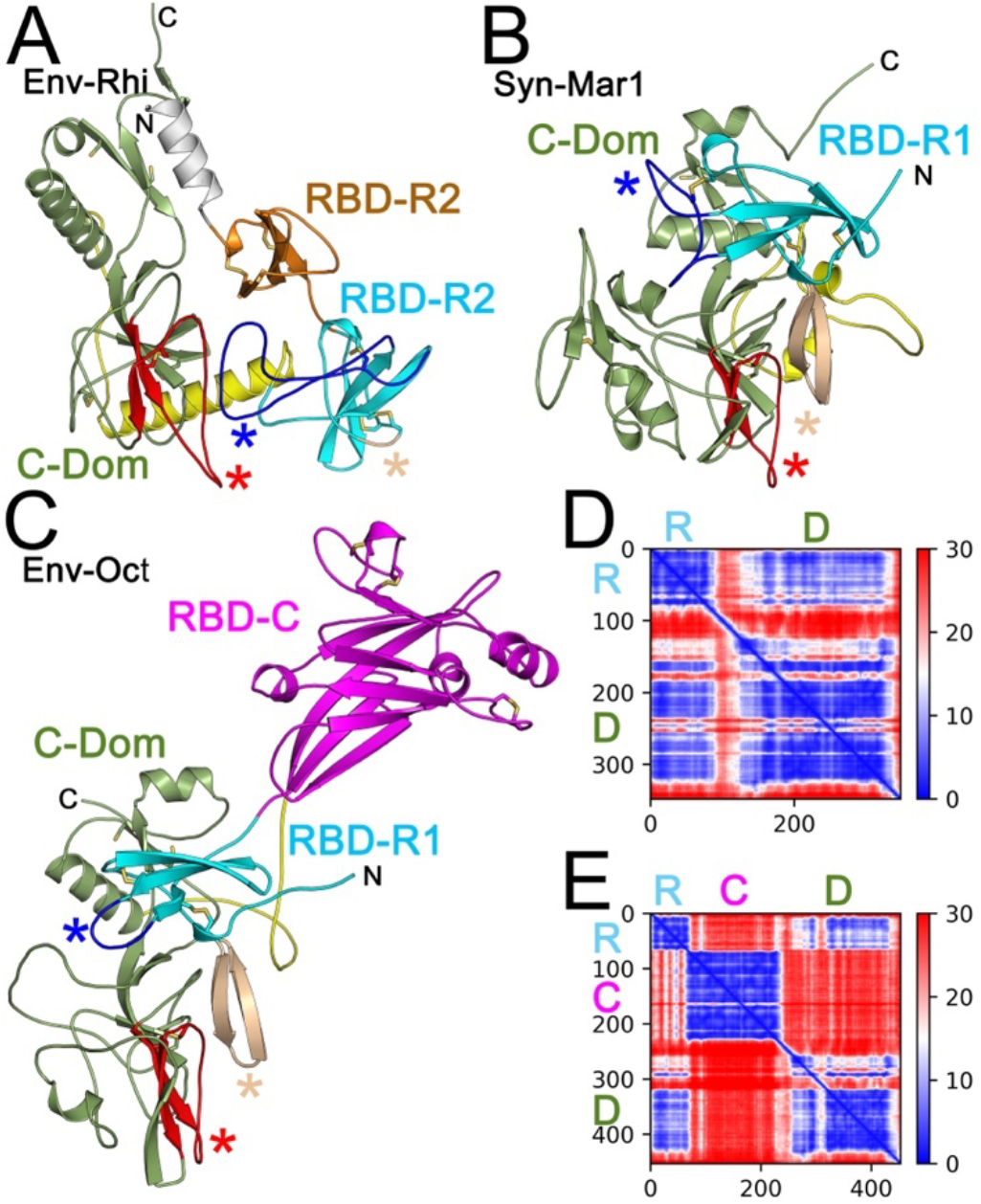
SU models of RDR and EnvR supergroup Env. (A) RDR Env-Rhi SU model, taken from the trimeric Env model (22). The core regions of the PD/C-domain are shown in olive green with the apical π5/π6 strands and connecting loop F shown in red. The first and second domains of the RBD-R2 region are shown in orange and cyan, with the linker between the RBR-R and C-domain (C-Dom) shown in yellow. The red, wheat and blue asterisks indicate the positions of the PD apical region, π1R/π2R-homology turn and π4R/π5R loop, respectively. The white helix is the amino-terminal region of SU not part of the RBD-R. (B) EnvR group 3 Syn-Mar1 SU model, colored as in panel A with the π1R/π2R hairpin shown in wheat. (C) EnvR group 4 Env-Oct SU model colored as in panel B with the RBD-C region between the RBD-R1 and C-domain shown in magenta. The SU models in panels A-C are shown in the same orientation, aligned through the core regions of the PD, in the opposite orientation as in Figures 6G and H, with the distal end of Env at the bottom of the figures. N and C indicate the SU termini. (D) PAE score matrix of the Syn-Mar1 SU model showing structural correlations among distant regions of SU. The error, in Å, is shown in the scale to the right. The approximate positions of the RBD-R and C-domain in the matrix are indicated by colored R and D letters. Numbers indicate amino acid positions within the model. (E) PAE score matrix of the Env-Oct SU model. The approximate positions of the RBD-R, RBD-C and C-domain in the matrix are indicated by colored R, C and D letters. Note that the correlation scores between the RBD-R1 and PD regions are low (good) for both models, whereas the RBD-C in the Env-Oct SU model is poorly correlated with either SU region, indicating low confidence for the position for the RBD-C region in the model.

The single RBD-R1 β-sheet is similar to the second β-sheet of the RBD-R2 domain of extant RDR retroviruses (23). Despite the overall structural similarity, a major difference is observed between the RBD-R1 of Groups 3 and 4 and the second β-sheet of the RBD-R2 of extant and endogenous RDR Env. A long loop emanating between strands β4R and β5R contacts strands β5 and β6 and loop F connecting these β-strands in the apical region of the PD in RBD-R2 in both monomeric SU and trimeric Env models (Fig. 6C and 7A, Suppl. Fig. 6C). In contrast, the RBD-R1 of EnvR groups 3 and 4 has a shorter loop between strands β4R and β5R and has instead the β-hairpin with strands β1R and β2R preceding the β-sheet contacting the same region of the PD (Fig. 6A and B, Fig. 7B and C, Suppl. Fig. 6A and B). RDR Env does not have strands β1R and β2R, with the chain section corresponding topologically to this region forming a short loop connecting directly into strand β3R of the β-sheet (Fig. 6C and 7A, Suppl. Fig. 6C). The topological organization of the RBD-R domain of Groups 3 and 4 is more similar to that of the RBD-R1 of Syn-1 except that in Syn-1 the β-hairpin region is modeled as a long and relatively disordered loop instead, and a close association between the RBD and PD is not observed in the Syn-1 model although this is associated with an overall lower model quality for Syn-1 (Fig. 6D, Suppl. Fig. 7D). Of note, the first β-sheet repeat of the RDR Env RBD-R2 region does not have either one the extended loops, with the exception of the REV RBD-R (Fig. 6E and Suppl. Fig. 6D). All these models have high local pLDDT scores except the first loop of the Syn-1 RBD-R1 (Suppl. Fig. 7A to E).

Differences in disulfide bonding are also observed among RBD-R domains. Two of the disulfide bonds are structurally conserved between the two RBD-R types (Fig 6A to D, disulfides a and b). However, the third disulfide bond varies structurally (Fig. 6A to D, disulfide c). In one, which includes the second repeat of RBD-R2 of RDR Env, Syn-1 and a Syn-Mar1 and a subset of other EnvR group 3 members, the disulfide links the β2R/β3R and β4R/β5R loops. In another subset of group 3 sequences the disulfide occurs between the β2R/β3R loop and β4R.

The major difference between the RBD-R of the EnvR group and RDR Env lies in the apparent position and orientation of the domain in trimeric Env and SU models (Fig. 6G and H and 7A to C). In the previously described trimeric RDR Env model, the second β-sheet repeat of the RBD-R2 domain, which contacts PD strands β5 and β6 and Loop F, forms part of the distal end of the trimer (Fig. 6G). Aligning the model of the SU of Syn-Mar1 through the conserved core region of the PD of SU to the RDR Env trimer model, PD strands β5 and β6 of Syn-Mar1 superpose to the β-strands β5 and β6 of RDR Env. However, the RBD-R1 β-sheet of Syn-Mar1, when SU is oriented in the trimer model by alignment with the trimeric RDR Env model, is located away from the top of the trimer, on the side of the trimer, projecting the β1R/β2R hairpin loop distally to contact PD strands β5 and β6 and Loop F (Fig. 6H). Although the first β-sheet repeat of the RDR Env RBD-R2 is also located in a more proximal region away from the top of the trimer, its location is distinct from that of the Syn-Mar1 RBD-R1 β-sheet, with an approximately 180° rotation relative to the expected orientation of the Syn-Mar1 RBD-R1 β-sheet (Fig. 6G and 7A). Thus, these two β-sheets do not appear to be structurally equivalent in trimeric Env. Based on the structural alignment with the RDR Env trimer model, the main interaction of the RBD-R β-sheet of the EnvR groups 3 and 4 with the rest of Env, excluding the β-hairpin contact with the PD, appears to be between the fusion peptide and an RBD-R1 loop section between strands β4R and β5R (Fig. 6G and Fig. 7B and C). Importantly, similar to the trimeric Env model, the PAE score matrices indicate good reliability (low scores) for the relative position between the RBD-R1 and C-domain regions for most of the EnvR group 3 and 4 SU models (Fig. 7D and E). Exceptions are the models for avian EnvR group 3 SU models, in which the RBD-R regions are in positions different from the one observed in Syn-Mar1 but have high (poor) PAE reliability scores for the positions of these regions relative to the rest of SU (Suppl. Fig. 5B).

Sequence and structure-based alignments show limited sequence similarity between RBD-R regions of members of different EnvR groups and the RBD-R regions of extant and endogenous RDR retroviruses (Fig. 8). The linker region immediately following the RBD-R1 domain of Syn-1 and RBD-R2 in extant and endogenous RDR Env has a SDGGGX_2_DX_2_R motif (10, 39). In the trimeric RDR Env model the non-Gly residues in this motif form several intrachain polar interactions within that region, whereas the consecutive conserved Gly residues allow a sharp turn in the polypeptide chain (22). This motif is followed by a region of strong α-helical propensity that is located at the top of the Env trimer. A similar motif is not observed in the EnvR group 3 and 4 Env, defining another important difference between the RBD-R of the EnvR group and that of Syn-1 and extant and endogenous RDR Env. This is consistent with the different position of the RBD-R in these two Env classes requiring different linker structures.

**Figure 8.**
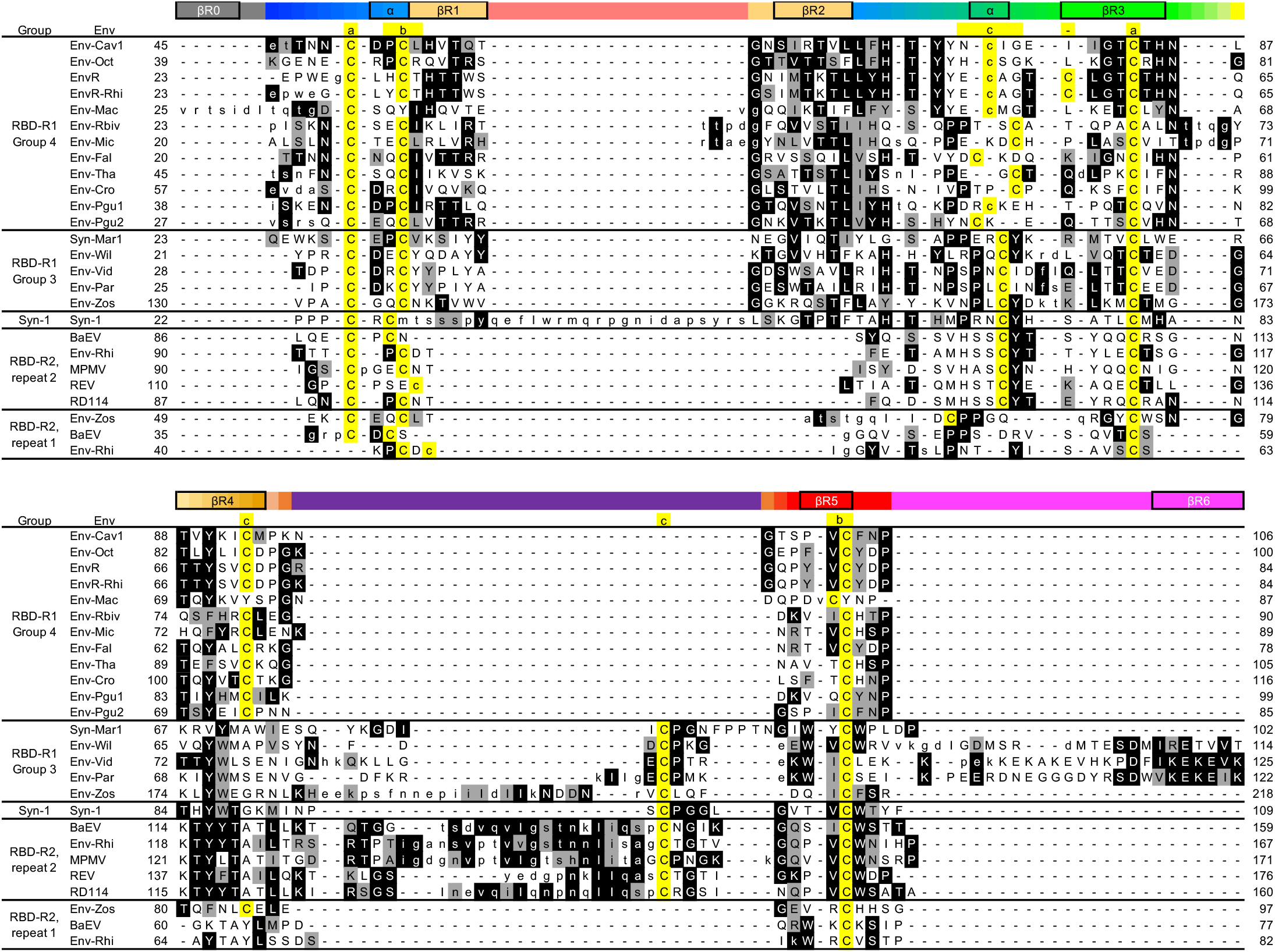
Structure-based alignment of RBD-R sequences. RBD-R1 sequences from EnvR groups 3 and 4 and RBD-R2 sequences from RDR interference group are structurally aligned, using the Env-Oct RBD-R1 sequence as reference. Lower-case characters indicate non-aligned residues. Residues highlighted in black are identical to the consensus sequence whereas gray backgrounds indicate similarity according to the BLOSUM-62 similarity matrix. The consensus is normalized by calculating the consensus of each group to derive the final consensus sequence. The positions of the β-sheets in the domain repeats are indicated above the alignment with coloring similar to that in Figure 6A-F. Cysteine residues are indicated in yellow with the disulfides observed in the models indicated by letters above the alignment as in Figures 6A-F. Residue numbering is from the Env initiation codon. Note that some avian RBD-R1 sequences in the EnvR group 3 have an extension including an additional β-strand at the end of the domain, highlighted in magenta above the alignment whereas Env-Mac has an amino-terminal extension with an added β-strand. The first and second RBD-R2 domain repeats are indicated in the leftmost column as repeat 1 and 2. Dashes indicate gaps.

### Group 4 Env has a dual RBD

Analysis of the SU structural models revealed a surprising RBD configuration in Group 4 Env. That group has the longest RBD region in SU in the EnvR supergroup. As described above, the SU of this group has an RBD-R1 at its amino-terminus. However, this does not account for the longer RBD sequence of this group. The rest of the sequence in the EnvR group 4 RBD region forms a second independently-folding RBD-C-like domain in all structural models (Fig. 2F and 7C). All the EnvR group 4 RBD-C models had high pLDDT scores, indicating good reliability for side-chains through most of the region except those of EnvR and the closely related EnvR-Rhi, which, although retaining a similar fold, had loop regions that were not well-resolved in the models. The model with the highest reliability score for the RBD-C section for this group was that of Env-Oct (Suppl. Fig. 1E), which is used as a prototype to describe the RBD-C of EnvR group 4.

The variable regions A and B (VRA and VRB) of all EnvR group 4 RBD-C structural models are more similar to those of Env-Pb(1) rather than the longer ones from MLV and FeLV RBDs (Fig. 2F). Thus, while similar to the RBD-C of extant Type-C retroviruses, the RBD-C of EnvR group 4 is more closely related structurally to the RBD-C of Env-Pb(1). The major difference between the EnvR group 4 RBD-C structural models and Env-Pb(1) is that whereas all β-strands of the Env-Pb(1) RBD-C conform to the canonical topology of IgV domains, the EnvR group 4 RBD-C structural models have an additional β-strand, strand 1β, at the beginning of the domain that inserts itself between strands 2β and 9β in an antiparallel configuration in the structural model, similar to the configuration of that region observed in the MLV and FeLV RBD-C structures (Fig. 2A, B and E). All EnvR group 4 RBD-C models and Env-Pb(1) had very similar folds, with the largest differences observed around the VRC region and the loop following strands 3β (Suppl. Fig. 1B). Thus, EnvR group 4 has a Pb(1)-like RBD-C region that also shares some structural features with the RBD-C Type-C structures.

Sequence similarities in structural alignments were mostly limited to core residues, with the RBD-C EnvR group sequences sharing more similarity with the Pb(1) than MLV RBD-C sequences (Fig. 3). However, the Type-C and EnvR group 4 RBD-C structures and models share a conserved group of 4 discontinuous residues proximal to the VRC region (Fig. 2A and F and Fig. 3). These correspond to residues Tyr-119, Trp-144, Trp-154 and Asp-217 of MLV Env. These residues are located in strand 4β, an α-helix just before strand 5β, within or just downstream of strand 5β and in a structurally conserved 8 aa-loop, packing together on the surface of the distal end of the MLV RBD (distal relative to the termini of the domain). Residues Tyr-119 and Asp-217 form a hydrogen bond whereas the two Trp residues pack against Tyr-119. Tyr-119 also packs against the disulfide formed by conserved Cys-121 and Cys-141 in strand 4β and just before strand 5β. Residues Trp-144 and Trp-154 are exposed on the surface of the RBD, creating a hydrophobic surface region on the MLV RBD, whereas Tyr-119 remains buried to form a scaffold to anchor Trp-144 and Trp-154. The same residues are conserved in the FeLV RBD structure (not shown) and in the EnvR group 4 RBD-C structural models, with the only difference being the exact position where the Trp-154 homologue is located, with some EnvR group sequences lacking this Trp residue. Although the Cα of Trp residues structurally similar to MLV Trp-154 are in different locations in different structures and models, the side-chains participate in the network of interactions similarly. Only three of these conserved residues are observed in the Env-Pb(1) RBD-C structure and the ZFERV RBD-C model, Tyr-119, Trp-144 and Asp-217, with the Tyr-119 and Asp-217 homologues also forming a hydrogen bond (Fig. 2B and C). However, in the Env-Pb(1), the side-chain of the Trp-144 homologue points in the opposite direction in both copies of the asymmetric unit of the crystal structure (Fig. 2B). None of the residues in the conserved tetrad are observed in the RBD-C Fc structural models and sequences (Fig. 2D and 3), with the partial exception of Env-Apo (Fig. 2E). Of note, the conserved tetrad is not necessary to maintain the core RBD-C β-sheet structure or to constrain the VRC region in place, given the different orientation of the Trp-154-like residue in the Env-Pb(1) structure and the complete absence of the tetrad in most RBD-C Fc structural models. These conserved residues are also not a general feature of IgV-like domain structures. Thus, the conserved tetrad appears to be a unique local surface structural feature of some RBD-C classes. The structural modeling results show a structural similarity between the RBD-C of EnvR group 4 models and the RBD-C of Env-Pb(1), with some sequence features being more universally shared among different RBD-C types. These structural features are not convergent, as the RBD-C of EnvR group4 Env and Env-Pb(1) share significant, if fragmentary, sequence similarities in both β-strand and inter-strand regions, strongly indicating a common ancestor (Fig. 3).

### Sequence of PD and RBD insertion events in the EnvR group

The sequence of internal insertions and deletions in the SU of the EnvR group is not clear from the TM phylogenetic tree due to the low support for the branching of groups within this supergroup. Thus, the TM-based analysis does not provide sufficient information to resolve Env clustering within the group. Therefore, phylogenetic analysis of the EnvR group was expanded by incorporating SU sequences in the alignments for more detailed phylogenetic tree construction.

Although SU sequences are not usually used for retroviral Env phylogenetic analyses due to SU phylogenetic signal being lost with the relatively high sequence diversity in more divergent Env sequences, Env sequences in this group show relatively conserved SU regions (Suppl. Fig. 4), suggesting that these regions might be useful for more detailed phylogenetic analyses on Env of this group. A phylogenetic tree was constructed using the TM ectodomain and core PD sections of SU from the apparent helix αPD1 (22) to strand β6 and between strand β7 to the SU C-terminus, excluding the putative RBD and the PD Expansion regions between strands β6 and β7. The TM ectodomain and core PD regions are relatively conserved in all Env sequences in this supergroup and, because they are mostly structurally independent from the PD expansion regions, should provide more detailed information about branching of different groups to determine insertion and deletion events without RBD and PD expansion region sequences biasing the results.

Maximum likelihood phylogenetic trees showed a clear clustering of sequences according to SU structural types (Fig. 9). Importantly, EnvR groups 1, 2, 3 and 4 formed well-supported individual clusters with bootstrap values of at least 90. Similar tree topologies and bootstrap support values were obtained with neighbor-joining phylogenetic trees (not shown). The only exception is Env-Ami sequence, classified as a group 3 sequence based on the location of the CXXC motif within SU, which clustered between the group 1 and 2 sequences. Thus, Env-Ami appears to be an anomalous EnvR group 1 sequence rather than a group 3 sequence, congruent with its minimal PD structure. The phylogenetic tree indicates the likely steps in PD and RBD evolution in the EnvR group. The basal sequences of the EnvR group have a PD with the short Expansion 1 region and no RBD. Two scenarios are possible for SU evolution within the EnvR group. In the first scenario, the next step in PD evolution was the insertion of Expansions 2 and 3 in the PD in the lineage leading to Groups 2, 3 and 4. Following the split of group 2 from the lineage leading to groups 3 and 4, the SU of group 2 then acquired expansion 4 of the PD, with only Env-Aqu subsequently acquiring Expansion 5 while deleting part of Expansion 4 (Fig. 9 and Suppl. Fig. 4). The branching of Env-Tac4.1 in the consensus tree would appear to be incompatible with this scenario. However, the support value for the point where Env-Tac4.1 splits from groups 3 and 4 is relatively low (Fig. 9), and examination of individual trees in the bootstrap repeats shows that Env-Tac4.1 often clusters with the other group 2 sequences in trees that differ from the consensus tree shown in Figure 9.

**Figure 9.**
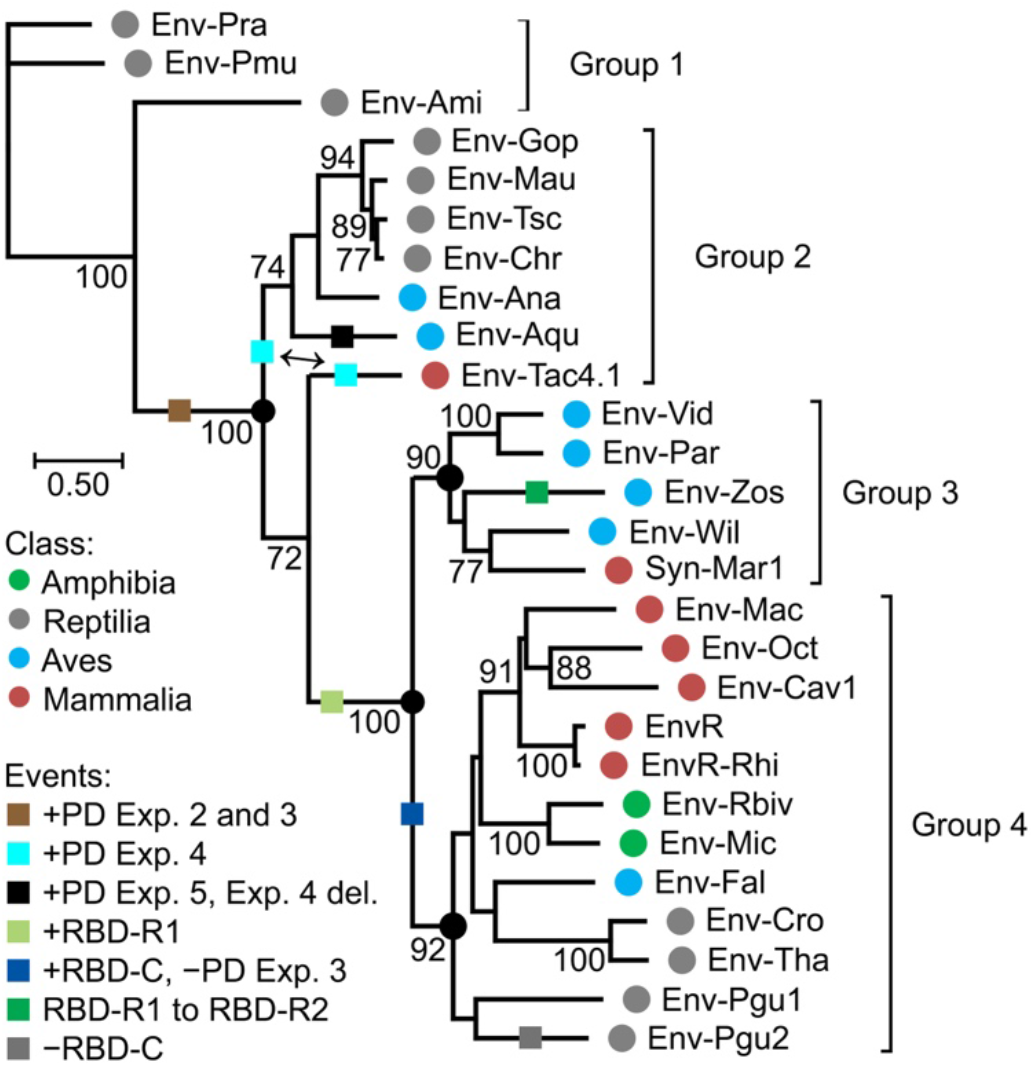
EnvR Group bootstrapped maximum likelihood phylogenetic tree. Env sequences used to construct the tree include the TM ectodomain and SU sequences excluding PD Expansions 1 to 5 and the RBD regions upstream from the putative helix αPD1 just before the CXXC motif. The tree is rooted at the midpoint. The four EnvR Groups are indicated on the right. Bootstrap values (100 repeats) greater than 70 are indicated on nodes. Black circles indicate the nodes defining three of the four major groups. Colored circles next to Env names indicate the tetrapod class for the species harboring the corresponding *env* gene as indicated in the legend to the left. Colored squares on branches indicate PD expansion and RBD insertion and deletion events as indicated in the legend to the left. The double arrow indicates PD Expansion region 4 events discordant with the consensus tree. The tree is drawn to scale with the distance scale indicating amino-acid substitutions per site. Env-Ami is re-classified here as a Group 1 outlier sequence.

In the above scenario, the SU of the common ancestor for EnvR groups 3 and 4 acquired an RBD-R1 region in the amino-terminal region of SU, leading to the origin of the EnvR group 3 Env configuration (Fig. 9). The branch leading to group 4 from this common ancestor underwent a deletion of Expansion 3 after the split from group 3, which retained this Expansion. In this process, group 4 also acquired a Pb(1)-like RBD-C (Fig. 9). The sequence of these two events in the origin of group 4 Env is not clear from the data. Other events in terminal regions of the tree are observed, including the conversion of the RBD-R1 into an RBD-R2 in Env-Zos and the loss of the RBD-C in Env-Pgu2 (Fig. 9). In the second scenario, the non-contiguous PD Expansions 2, 3 and 4 occurred in a common ancestor of groups 2, 3 and 4, with subsequent loss of Expansion 4 in the branch leading to groups 3 and 4, loss of Expansion 3 in group 4, and acquisition of RBD-R and RBD-C regions as in the first scenario. Of these two scenarios, the first one shown in Figure 9 is more parsimonious, requiring fewer PD deletion steps. In either scenario, PD evolution in the EnvR group was associated with both acquisition and subsequent loss of relatively large expansions in the apical region between strands β6 and β7 of the PD. Importantly, each of the PD expansion and RBD types, except for the minimal PD of group 1 and expansion 5 in Env-Aqu, is present in Env sequences from species in 2 to 4 different tetrapod classes, and thus these different PD expansion and RBD types were probably acquired by ancestral retroviral lineages and diffused by viral transmission rather than arising by evolution of endogenous Env sequences after germline integration. Of note, the internal PD expansions in the EnvR supergroup are structurally distinct from those in Syn-2, indicating independent origin for the expanded PD regions in these two Env lineages, consistent with the phylogenetic data of TM regions.

### Acquisition of an RBD-like sequence from a mammalian host by an alpharetrovirus-like Env

The results described above indicate relatively widespread intragene *env* recombination among widely distinct retroviruses leading to evolution of Env proteins with different RBDs. The RBD-C has an IgV-like structure, modified in its amino-terminal and sometimes carboxy-terminal β-strands as well as with significant expansion of loops analogous to the variable loops of immunoglobulin variable regions. However, whether the RBD-C was ultimately derived from a vertebrate IgV domain by recombination between retroviral *env* and host cell genes is not known due to the structural and sequence differences between RBD-C and vertebrate IgV domains. It is possible nonetheless that RBD-C regions were coopted from host genes, similar to the acquisition of oncogenes from vertebrate hosts by oncogenic retroviruses.

This question was addressed by an extensive structural survey of gamma-type SU by AF2 modeling. No RBD-C or RBD-R structural models distinct from those described here were identified for gamma-type Env with a CXXC motif in SU.

However, structural modeling of the SU of Env sequences without a CXXC motif from mammalian species that cluster with the avian gamma-type Env in TM phylogenetic trees identified an amino-terminal IgV domain distinct from other RBD-C structures. These include Env sequences from the Pacific pocket mouse *Perognathus longimembris pacificus* (Env-Per), the south-central black rhinoceros *Diceros bicornis minor* (Env-Dbi), two pangolin species, *Manis javanica* and *M. pentadactyla*, and the European rabbit *Oryctolagus cuniculus*, the latter including a full SU and partial TM sequence (Table 1, Suppl. Fig. 8 and Suppl. File 1). These Env sequences have features more typical of alpharetroviral (avian) gamma-type Env sequences with which they cluster in TM phylogenetic trees, including no CXXC motif in SU and two Cys residues in the fusion peptide in the amino-terminal region of TM (Fig. 1, Suppl. Fig. 8). In the Env-Per SU model a typical minimal PD structure is observed (Suppl. Fig. 9). The five putative RBD structural models associated with these Env sequences are similar. The Env-Dbi RBD is the best model for this set of sequences and is described below as the prototype for this RBD class (Fig. 10A and D).

**Figure 10.**
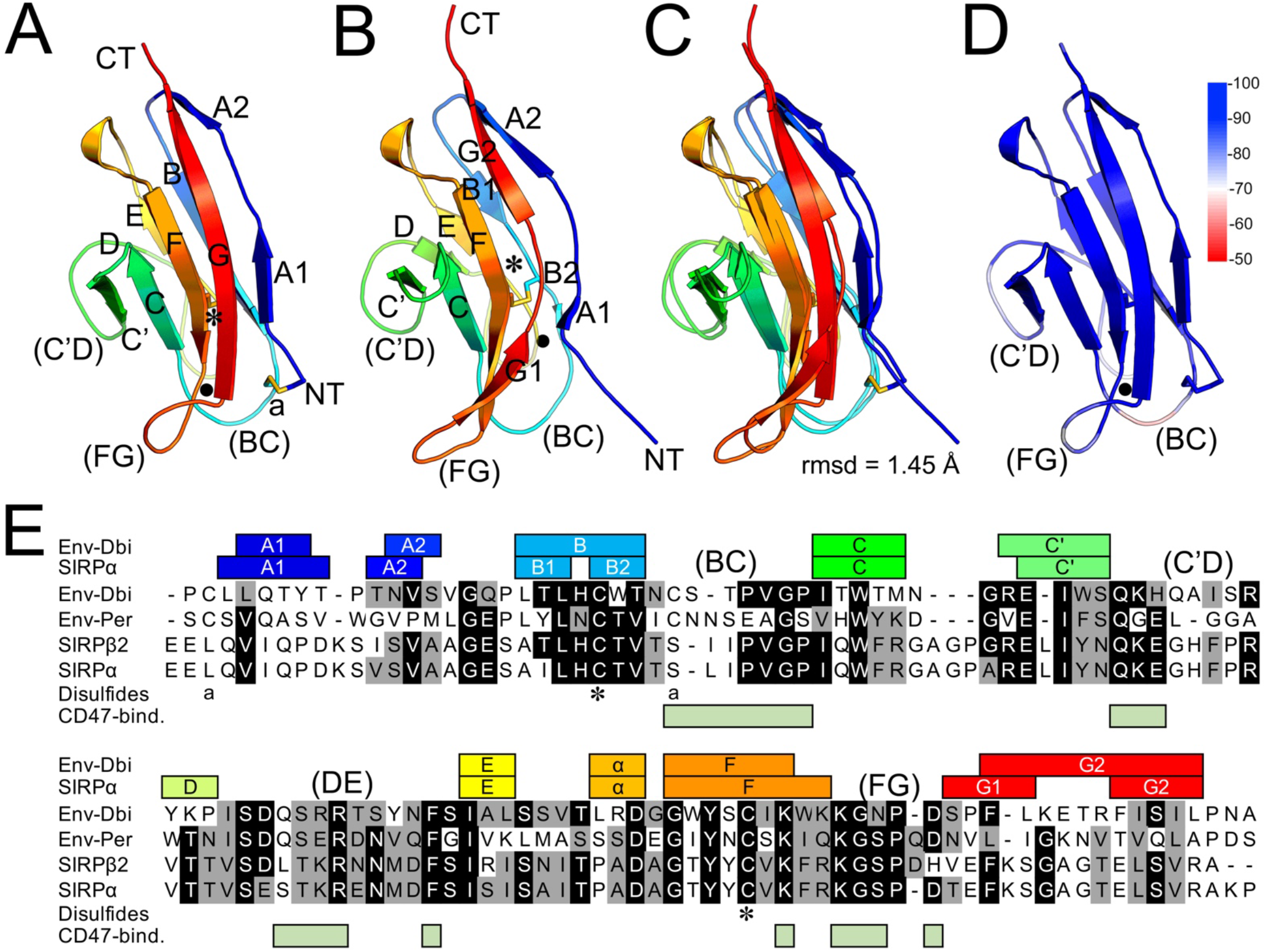
Env-Dbi RBD structural model and sequence. (A) Env-Dbi RBD structural model generated *ab initio*. The chain is colored in a rainbow pattern, starting with blue at the amino-terminus. The A-G labels indicate β-strands according to the IgV domain naming convention. Disulfides are shown as sticks, with the canonical IgV domain strand B-F disulfide indicated with an asterisk. Selected inter-strand loops are indicated in parentheses except for the (DE) loop, indicated with a black dot. (B) Crystal structure of the first SIRPα domain (PBD 2JJS, chain B), in the same orientation and with the same labeling as the Env-Dbi RBD model in panel A. Only the SIRPα section of the complex structure is shown. (C) Alignment of the model and structure in panels A and B. (D) Local pLDDT reliability scores along the chain of the Env-Dbi RBD structural model. The scale to the right indicates pLDDT scores. Only selected loop regions are labeled as in panel A. (E) Alignment of the sequences of the Env-Dbi and Env-Per RBD regions and the first SIRPα and β2 domain sequences. Black and gray backgrounds indicate residue identity and similarity between the Env-Dbi/Env-Per and SIRPα/β sequences according to the BLOSUM-62 matrix to highlight similarities between groups. The β-strand regions of both sequences are shown above the alignment in the same coloring as panels A and B with the main loops labeled. The cysteine residues forming the canonical IgV strand B-F disulfide are indicated by asterisks. The cysteine residues forming an additional disulfide are indicated with the letter “a” below the alignment. The light green boxes below the alignment indicate SIRPα residues in contact with its ligand CD47 in the complex structure.

The structural model of the putative amino-terminal RBD of Env-Dbi adopts a typical IgV-like domain conformation with β-strands and the canonical IgV domain disulfide bond between β-strands B and F (β-strands names according to the IgV domain system) that is absent in the corresponding β-strands of retroviral RBD-C structures and models (Fig. 10A and E). The Env-Dbi RBD also lacks the VRA, VRB and VRC loop expansions of other RBD-C structures and models. Structural alignment of the Env-Dbi RBD model to structures in the Protein Data Bank using the Foldseek server identified the first domain of mammalian signal regulatory proteins (SIRP) α, β and γ structures, which have canonical IgV domain structures (40), as the closest structural homologs (Fig. 10B). Thus, this RBD class is named here RBD-C SIRP-like. The root mean-square deviations of the structural alignments of the Env-Dbi RBD model with human SIRPβ(2) and SIRPα were 1.297 and 1.45 Å, respectively, with the main deviations being observed in loop regions of lower reliability scores in the RBD structural model (Fig. 10C and D). Notably, the RBD sequences in this class can be clearly aligned to that of SIRPα and related proteins in this family (Fig. 10E). In fact, GenBank BLAST searches with the RBD sequences of this group yield, after the related retroviral Env sequences of this group, sequences for SIRP homologues from mammalian species but no matches to proteins from any other taxon or retroviral protein except for a few likely environmental contaminants in ticks. In addition, up to 12 of the 19 residues that are involved in binding of SIRPα to its ligand, CD47 (40), are strictly conserved in Env-Dbi and other proteins of this group (Fig. 10E). The sequence similarity between the RBD-C SIRP-like variants and SIRPα, β and γ starts in the second section of bipartite strand A and ends in strand G at the end of the domain. No sequence similarity is observed upstream from strand A, including the signal sequence preceding the domain, although all of strand A is present in the RBD structural models. The RBD-C SIRP-like models have an additional disulfide not observed in typical IgV-like domains, between cysteine residues preceding and following strands A and B respectively, although only some models show this disulfide formed (Fig. 10A and E). The conserved RBD-C tetrad is not observed in the RBD-C SIRP-like sequences. The structural modeling and sequence analyses indicate a mammalian origin for the RBD-C SIRP-like class.

### Retroviral origin of Env-Dbi

The retroviral origin of Env sequences with SIRP-like RBD-C moieties was addressed by searches of proviral sequences flanking *env* genes. No clear flanking proviral sequences associated with Env-Per, Env-Mja, Env-Mpe or Env-Ory were identified. In contrast, the *env* gene encoding Env-Dbi was found to be embedded in an apparently complete provirus, named here ERV-S.1-Dbi, in the *D. bicornis minor* whole genome shotgun (wgs) contig JAJIAZ010000042.1. This provirus is distinct from recently described gammaretroviral proviruses identified in *D. bicornis minor*, including one in a different position of the JAJIAZ010000042.1 wgs contig (41). The ERV-S.1-Dbi provirus has identical flanking 792-bp long-terminal repeat (LTR) sequences, complete *gag, pol* and *env* genes in the same reading frame separated by stop codons and a primer binding site (PBS) with a perfect match for the 3’ end sequence of the human tRNA^Ser^ with a CGA anticodon (Suppl. Fig. 10). A CA dinucleotide inverted repeat and a 5-nucleotide direct repeat immediately flank the provirus. ERV-S.1-Dbi is integrated into an intron between the fifth and sixth exons of a pseudogene homologous to the acyl-CoA synthetase medium chain family member 1 (*acsm1*) gene in the same orientation as the pseudogene. Additional complete and partial copies of the provirus were identified in *D. bicornis minor* whole genome sequences, including one in wgs contig JAJIAY010000041.1 in the intergenic region between the *znf280d* (XM_058541659) and *tcf12* (XM_058541653) genes (Suppl. Fig. 10). Nearly identical proviruses were also identified in whole genome contigs of the eastern black rhinoceros *D. bicornis michaeli*, including one in wgs assembly JANTPW010000004.1 corresponding to the same integration event as the *D. bicornis minor* JAJIAY010000041.1 provirus, differing from it by 17 nucleotide substitutions and 4 deletions (Suppl. Fig. 10). The identical flanking LTR sequences and the seemingly intact proviral genes and provirus structure indicate that the germline integration of ERV-S.1-Dbi within the *acsm1* pseudogene occurred relatively recently, although the virus originating ERV-S.1-Dbi was circulating in black rhinoceros populations prior to the estimated divergence of *D. bicornis minor* and *D. bicornis michaeli* about 1 Mya (42).

The closest homologues for the Gag and Pol polyproteins of ERV-S.1-Dbi were amphibian and mammalian endogenous retroviral proteins related to muERV-L and the Friend virus susceptibility 1 restriction factor Fv1 (43, 44), confirmed by phylogenetic analyses of Gag and reverse-transcriptase (RT) protein sequences (Fig. 11). Neither murine muERV-L nor the related human HERV-L include an *env* gene whereas the related HERV-S includes three canonical retroviral genes with multiple stop codons and short deletions interrupting open reading frames (45). Thus, ERV-S.1-Dbi is the first known provirus related to muERV-L, HERV-L and HERV-S with a full set of intact retroviral genes and seemingly intact proviral structure. In addition, the results indicate that alpha-like (avian) gamma-type Env sequences have been associated with retroviruses outside extant retroviral groups that were circulating as exogenous viruses relatively recently. Of note, no proviral context information is available for the other two currently known divergent Alpha-like Env, Env-Mab3 and Env-Mab-4.

**Figure 11.**
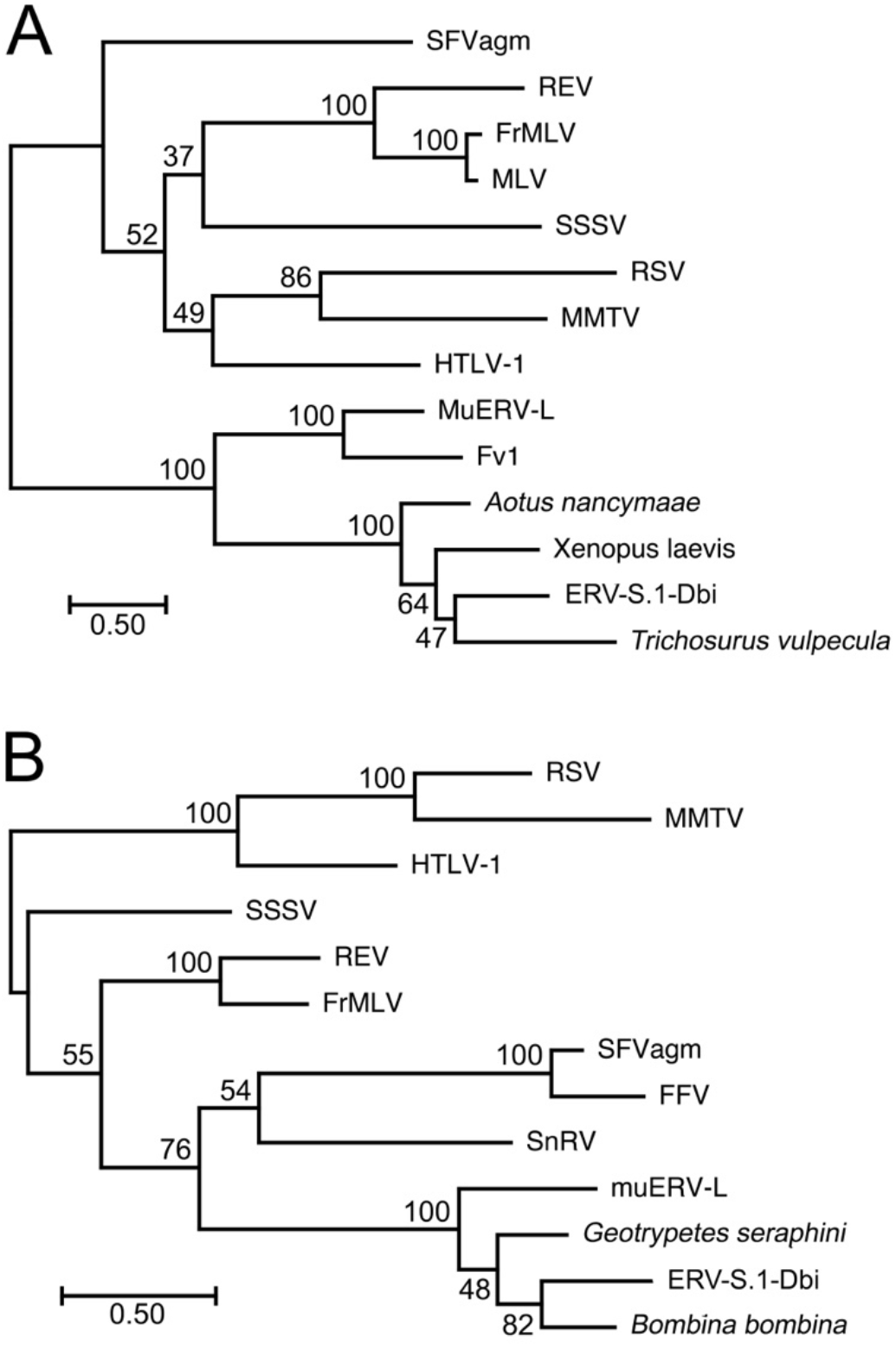
Phylogenetic tree of Gag and Pol protein sequences. (A) Gag polyprotein maximum likelihood phylogenetic tree. (B) Maximum likelihood phylogenetic tree of the RT region of the Pol polyprotein. Both trees include representatives of five extant orthoretroviral genera, spumaretroviruses and selected endogenous elements related to muERV-L and Fv1. Numbers in nodes are bootstrap values from 100 repeats. The trees are mid-point rooted and drawn to scale. The distance scales indicate amino-acid substitutions per site. FrMLV, Friend murine leukemia virus. RSV, Rous sarcoma virus. SnRV, snakehead retrovirus. MMTV, mouse mammary tumor virus. SFVagm, African green monkey simian foamy spumaretrovirus. FFV feline foamy spumaretrovirus.

## DISCUSSION

The SU structural modeling and Env phylogenetic analyses described here show that RBD moieties have been transferred between widely divergent Env groups, leading to the emergence of new combinations of RBD and PD/C-domain moieties in SU during Env evolution, with the emergence of dual RBD Env types in the EnvR supergroup. These RBD transfers are analogous to *env* gene transfers among distinct retroviral lineages but limited to sections of the *env* gene that encode the SU region rather than the entire *env* gene. Thus, intragene *env* recombination appears to have been a major mechanism of diversification and emergence of Env types in the evolution of retroviruses. In addition, the results show, for the first time, the acquisition of an RBD moiety by a retroviral Env from a host gene. This acquisition event is supported not only by high-quality structural modeling of the RBD showing a canonical IgV domain structure nearly identical to the mammalian SIRPα, β and γ immune signaling proteins, but also by the significant sequence similarity between the RBD and SIRP proteins, including residue identities in the ligand-binding loops of SIRPα.

The transfer of RBD moieties between widely divergent Env groups rests partially on the phylogenetic analyses of TM regions of Env. Such phylogenetic trees can be sensitive to the number and diversity of TM sequences that are included in these analyses. However, while the branching of taxons in the terminal regions of the Main Group and the EnvR supergroup vary, the branching of the major groups remains stable when adding or removing taxons. In addition, the position of Env types without independently-folding RBD moieties in the tress remains the same, basal to Env types with RBDs within the Main and EnvR groups. The most parsimonious interpretation of this is that the primordial Env for each of these groups did not have independently-folding RBDs outside the PD/C-domain and that these emerged later within each major group, including by RBD transfer between groups.

A second finding further supports the transfer of RBD moieties between divergent Env groups. There are two major RBD types in retroviral Env, the RBD-C and RBD-R classes. These are structurally and topologically distinct and do not appear to have evolved from each other. The RBD-R is structurally simple, composed of a single or repeated 3-strand β-sheet. Despite its simplicity and the lack of major sequence similarity in widely divergent Env groups, all RBD-R β-sheets have a similar arrangement of β-strands and pattern of 3 disulfide bonds that support their origin from a single ancestor and evolution by internal duplication. The RBD-C types also appear to have a single origin by structural and sequence analysis and the presence of the conserved tetrad. The presence of both RBD classes in individual Env within each of the two major Env groups and the existence of dual RBD Env types in the EnvR supergroup support the transfer of these regions among widely divergent Env. This is further corroborated by the identification of specific RBD subtypes, such as the RBD-C Pb(1), by both structural modeling and analysis of sequence alignments, shared between divergent Env types. The alternative interpretation, that RBDs were present in a primordial Env followed by convergent evolution of an RBD-C from an RBD-R, or vice-versa, with very similar structures and sequence similarities, is highly unlikely. Finally, that these RBD transfer events occurred during active viral replication rather than after germline integration is supported by the presence of each of the RBD combinations in the EnvR supergroup in at least one species from 3 or 4 major tetrapod classes.

An incidental finding in this work was the definition of internal PD expansions in the EnvR supergroup. These are independent from the RBD acquisitions and appear to be structurally distinct from the PD expansions observed in Syn-2. It is interesting that the major expansion of the PD in the EnvR supergroup, Expansion 4 in group 2, is not associated with an independently-folding amino-terminal RBD, whereas the EnvR group 3 and 4 Env with an RBD either do not have the large Expansion 4 or have lost Expansion 3 in the process of RBD acquisition. This suggests a functional equivalence between some of the PD expansions and RBDs, and two different paths for the evolution of receptor-interacting regions in the Env of retroviruses. The structural plasticity of the PD is enabled by its structural modularity, which allows the apical regions to be significantly modified while leaving the core regions of the PD interacting with TM unaffected.

The acquisition of the RBD SIRP-like class by an alpha-like Env appears to have occurred relatively recently, as several residues that in SIRPα contact its ligand, CD47, are conserved in the RBD. Conservation of these receptor-binding residues despite the considerable divergence of RBD-C SIRP-like sequences elsewhere in the domain suggests selection for receptor binding. Of note, the Env’s with the RBD-C SIRP-like domains may not necessarily use CD47 as a receptor, as even small amino acid differences in the interface in SIRPβ and γ are incompatible with binding to CD47 (40). The acquisition of this RBD by an ancestral retrovirus rather than by recombination after germline integration is supported by the fact that this RBD class is observed in endogenous Env’s in the genomes of species from 4 different mammalian orders, with one embedded in an apparently very young provirus. Although the RBD-C and SIRP-like RBDs may not share a common ancestor within the retroviruses, the identification of the SIRP-like RBD in alpha-like Env variants suggests that the RBD-C classes in extant retroviruses and the EnvR supergroup may also have originated by acquisition of an unidentified vertebrate host gene encoding an IgV domain protein by an ancient retrovirus. Whether the RBD-R class has a host origin is not clear. Its structure is relatively simple compared to the RBD-C structures and may have originated *de novo* in the retroviruses, although only once, followed by transfer, duplication of the domain and structural modification of its interaction with the rest of SU.

The results indicate that intragene *env* recombination has been an important factor in the long-term evolution of Env in the retroviruses. The exact direction and specific retroviral types participating in these partial *env* gene transfers is not clear from the data presented here and may involve other Env groups not analyzed here. The examples described here may be only the most obvious examples of intragene *env* recombination due to these occurring between widely distinct retroviral Env classes or yielding clear hybrid structures such as the dual RBD Env types. However, Env evolution by intragene *env* recombination may be more common but more difficult to ascertain in more closely related Env types, one possible example being Env-Apo and the Env-Fc1 group (Fig. 1 and 2E). Identification of recombination events between relatively similar Env types will require detailed analyses of Env sequences and associated proviruses and identification of close relatives of putative parental sequences.

The functional role of the RBD-C in retroviral infection has been well-documented, with less data available for the functional role of the RBD-R beyond cell binding (3, 10, 14, 39, 46–48). A unique feature of Env with RBD-C is the ability of soluble RBD moieties to functionally complement homologous Env mutants with non-functional RBDs within the Env trimer or with deleted RBDs (14, 46–48). That is, covalent linkage between the RBD-C and the rest of the Env protein is not necessary for infection. Any key interactions between the RBD-C and C-domain necessary for infection must be non-covalent, probably mediated by C-domain regions including the apical β5/β6 loop F of the PD (48). The acquisition and transfer of RBD-C moieties in the retroviruses and available functional data suggest a model for the steps in the functional evolution of Env (Fig. 12). A simple primordial Env with an SU lacking an RBD or one with an RBD-R1 would interact with its cognate receptor through a receptor binding site (RBS) in the PD/C-domain and/or RBD-R1 to trigger cell fusion by TM (Fig. 12A). It is unlikely RBD moieties transferred among widely distinct Env types or acquired from a host gene would be able to immediately interact with other SU regions to specifically mediate infection. More likely, these newly acquired RBD moieties would have a more generic function such as enhancement of cell binding through a second receptor, with the interactions with the original receptor mediated by the SU regions of the primordial Env needed for infection (Fig. 11B), analogous to recombinant MLV Env fused to erythropoietin (46). With time the RBD, bound to the second receptor, may have become itself a receptor for the rest of SU, which would switch from its primary receptor to receptor-bound RBD as its “receptor” for infection (Fig. 12C). This model would be compatible with the ability of RBD-C to mediate infection of engineered retroviruses with RBD-defective Env or infection of T-cells by FeLV-T using an endogenous RBD-C moiety, FELIX, as coreceptor in *trans* (46, 47, 49) (Fig. 12D). This model extends a similar model proposed by Barnett *et al*. (46) to include likely steps in the acquisition and functional integration of RBD moieties with the rest of Env to effect receptor-dependent membrane fusion. A tantalizing possibility is that the initial acquisition of an RBD from a cell gene might have initially allowed the coreceptor activity of an immunologically silent RBD to more effectively evade immunity by attaching the virus to the cell even in the presence of neutralizing antibodies to other SU regions, allowing rapid infection of cells through the original receptor once neutralizing antibodies transiently dissociate from SU. The apparently redundant presence of RBD-R and RBD-C moieties in one Env would be explained by the second RBD (RBD-C) having a more generic adhesin function that enhance the established functions of SU through the RBD-R1 moiety as described above. Whether RBD-R acquisition involves the same evolutionary steps is less clear due to the relatively limited functional data available with this type of Env.

**Figure 12.**
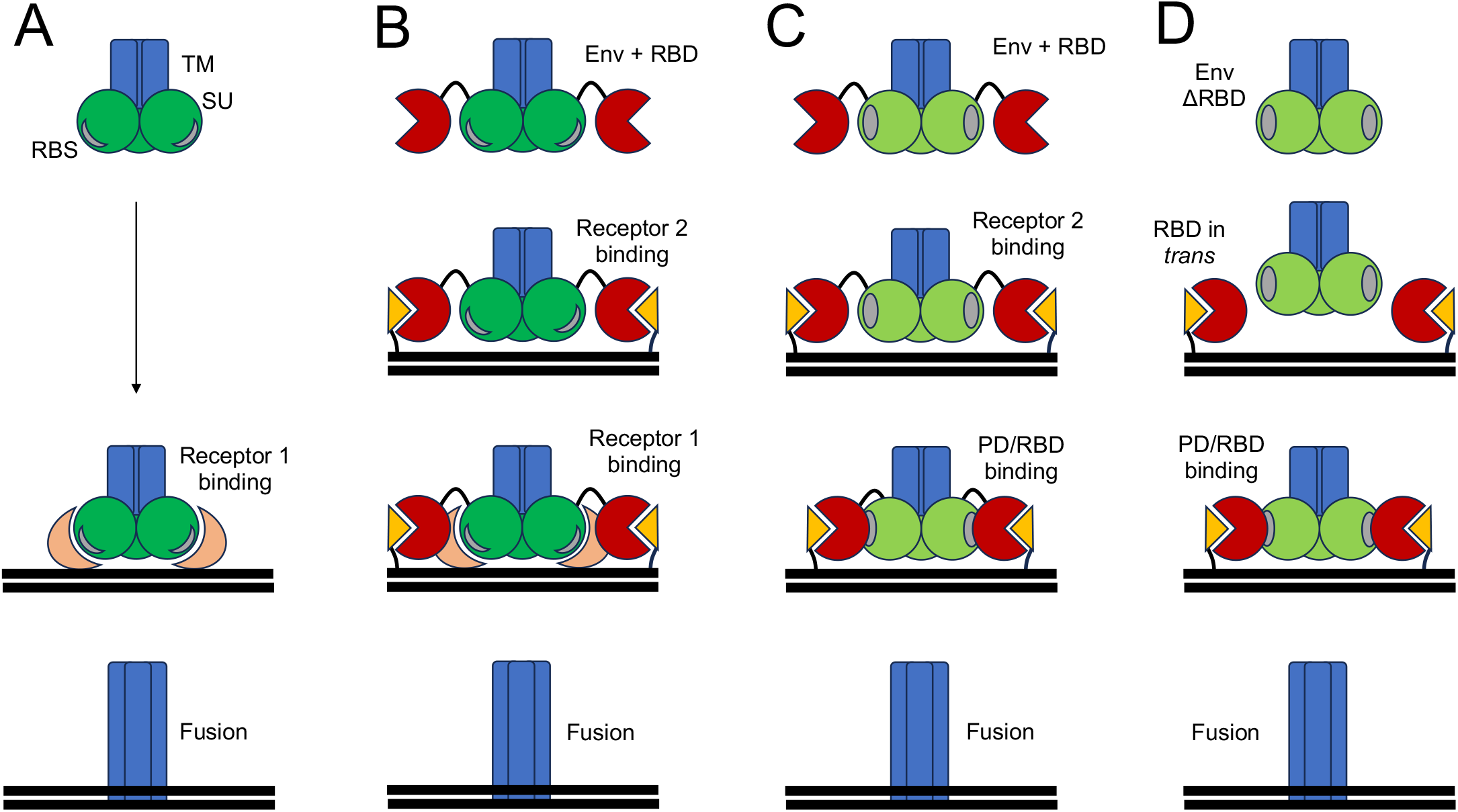
Model for functional integration of acquired RBD-C moieties into Env. (A) Primordial Env without an RBD-C and an SU comprised of a minimal PD/C-domain or an RBD-R1/PD that binds directly to its cognate receptor 1 for triggering of membrane fusion by TM. (B) Retroviral Env with a newly-acquired RBD binding a second receptor to enhance cell adsorption and RBD-independent interaction of the PD/C-domain of SU with its cognate receptor 1. There is no interaction between the RBD and receptor 1. (C) Modified Env binding only receptor 2 and use of the receptor-bound RBD as a receptor-like moiety for the PD/C-domain of SU in a manner analogous to interaction with primordial with receptor 1. (D) Artificial (*in vitro*) use of soluble RBD in *trans* for infection by a retrovirus with a deleted or defective RBD using the same interactions observed in panel C. Receptors 1 and 2 are shown as salmon crescents and orange triangles. The receptor and RBD-binding sites in the PD/C-domain of SU (green) are shown as gray crescents in the primordial Env and ovals in the evolved RBD-dependent Env, respectively. Infection events shown from top to bottom in each panel. The mechanism in panel D is based on existing experimental data. The mechanism in panel C is an extension of the mechanism in panel D. The mechanism in panel A is based on the existence of several retroviral Env without amino-terminal RBDs. The mechanism in panel B is hypothetical, to account for an autonomous function of a newly-acquired RBD not yet fully integrated functionally into the rest of Env. The virus, not shown, is on top of each figure. SU and RBD binding geometries in the figure are shown solely for display.

The results presented here define RBD classes and subclasses, expand the number of gamma-type Env structural types to include variants with multiple receptor-binding domains and adds acquisition and transfer of RBD moieties by intragenic *env* recombination as a mechanism of retroviral and Env diversification. The results also show clear evidence that RBD capture from a host gene has occurred at least once in the retroviruses, uncovering a previously unanticipated mechanism of Env diversification in the long history of retroviral evolution.

## MATERIALS AND METHODS

### Retrieval of full-length Env sequences from GenBank

Retroviral Env sequences were retrieved from the non-redundant section of GenBank by BLAST searches with multiple extant and endogenous retroviral Env sequences. Full-length sequences were further selected by locating amino-terminal signal sequences using the SignalP 5.0 server (https://services.healthtech.dtu.dk/services/SignalP-5.0/).

Sequences without full TM ectodomains were excluded from the dataset. Sequences were examined for potential CXXC motifs and Env cleavage sites preceding a potential fusion peptide manually. Selected Env sequences were grouped by closest homology to the TM of reference Env sequences by BLAST alignments.

### Structural modeling of SU

Structural modeling of SU was done with the ColabFold implementation of AF2 available at https://colab.research.google.com/github/sokrypton/ColabFold/blob/main/AlphaFold2.ipynb as previously described (23, 50). Models were obtained without templates and without amber relaxation. pLDDT scores above 90 indicate high side-chain reliability; scores between 70 and 90 indicate high main-chain reliability; scores between 50 and 70 indicate moderate reliability; scores below 50 indicate poor reliability (18, 19).

### Structural alignments

Multiple structural alignments of RBD models and structures were obtained using the DALI server (http://ekhidna2.biocenter.helsinki.fi/dali/) (51). Searches for structural homologues of RBD models in the Protein Data Bank were done using the FoldSeek server (https://search.foldseek.com/search) (52).

### Phylogenetic analyses

TM phylogenetic trees were constructed using TM ectodomain sequences from the last four SU residues including the Env cleavage site to the residue preceding the membrane-spanning region. Env phylogenetic trees of the EnvR supergroup were constructed using sequences from the TM ectodomain and SU sequences excluding RBD and linker regions and PD expansion regions, determined by structure-aided multiple sequence alignments (Suppl. Fig. 4). Retroviral sequences in GenBank accession ASH96781, CAA77478, AAB59942, NP_056887, CAA73250, NP_034374, YP_001956721, XP_041431051, YP_443922, XP_012328208, XP_036594074, AAD50661, NP_056880 and the ERV-S.1-Dbi Gag sequence in Supplementary Figure 10 were used in Gag polyprotein phylogenetic analyses. Retroviral sequences in GenBank accession ASH96780, CAA77477, NP_056886, CAA73251, YP_001956722, XP_033785733, XP_053567042, YP_443922, AAD50663, NP_056880, NP_043924, QZU26786 and the ERV-S.1-Dbi Pol sequence in Supplementary Figure 10 were used in RT phylogenetic analyses. Multiple sequence alignments were produced with the MAFFT v7 program available at https://toolkit.tuebingen.mpg.de/tools/mafft (53, 54). Maximum Likelihood phylogenetic trees were generated with the IQ-TREE 2 program (55) using automatic model selection, with 100 bootstrap replicates and the aBayes method to estimate branch support (56). Phylogenetic trees were rendered using the MEGA11 software (https://www.megasoftware.net).

## Data availability

Coordinates for the structural models described here are available upon request. Env sequences described here are included and annotated in Supplementary File 1. Multiple sequence alignment files used to construct phylogenetic trees are in Supplementary Files 2 (TM ectodomain), 3 (PD-TM ectodomain of EnvR supergroup), 4 (Gag polyprotein) and 5 (RT).

