## Supplementary Figures for "Transfer and Capture of Envelope Protein Receptor-Binding Domains in the Retroviruses"

Isidro Hötzel

*Millbrae, CA 94030*

### Supplementary Figures

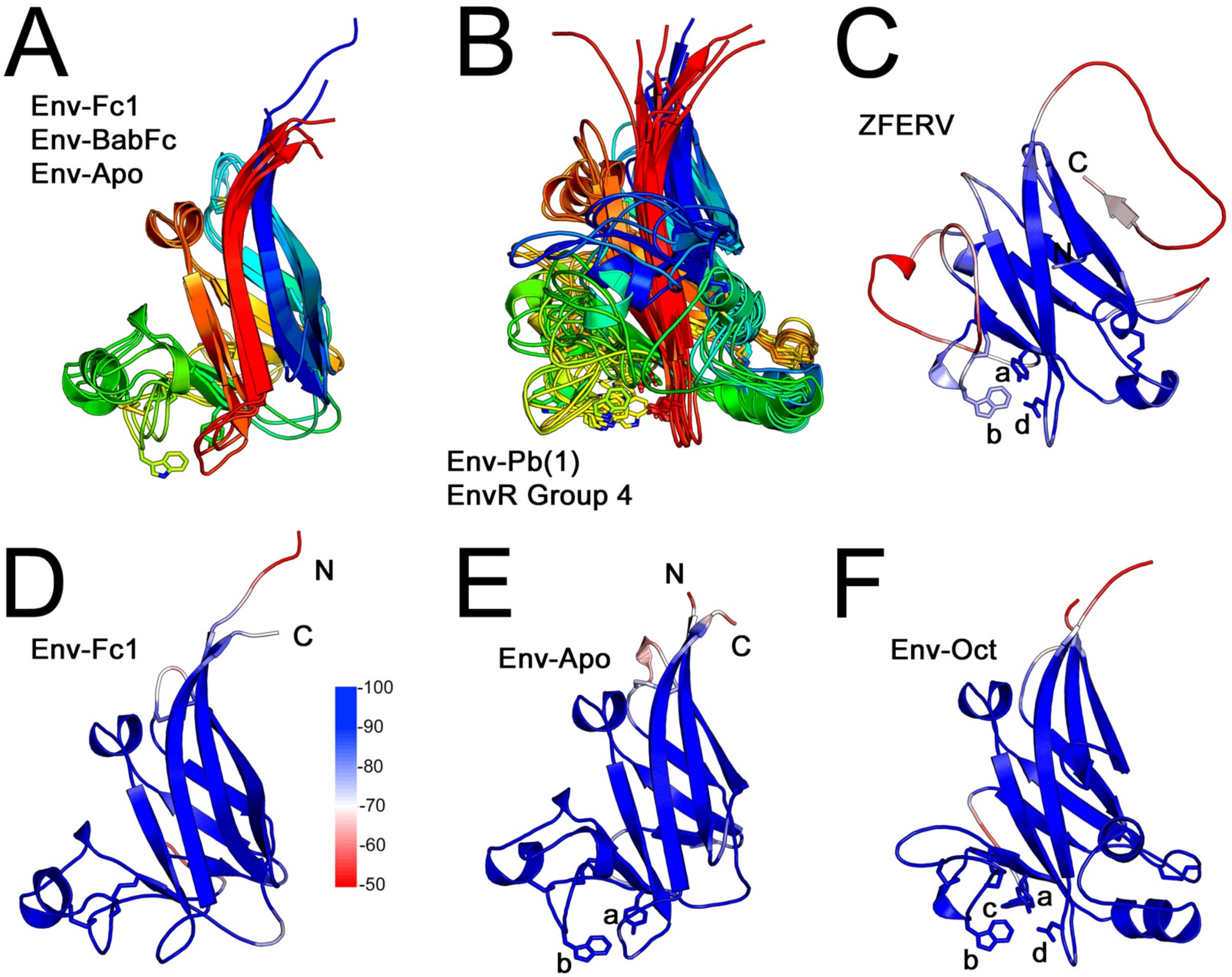


**Supplementary Figure 1** – Alignment of RBD-C models and model quality. Superpositions for (A) RBD-C Fc and (B) RBD-C Pb(1) structural models in the same orientation and coloring as in Figure 2. (C-D) pLDDT scores along the indicated RBD-C models. The scale to the right of the model in panel D indicates pLDDT scores. Residues in the conserved tetrad as shown as sticks and labeled.

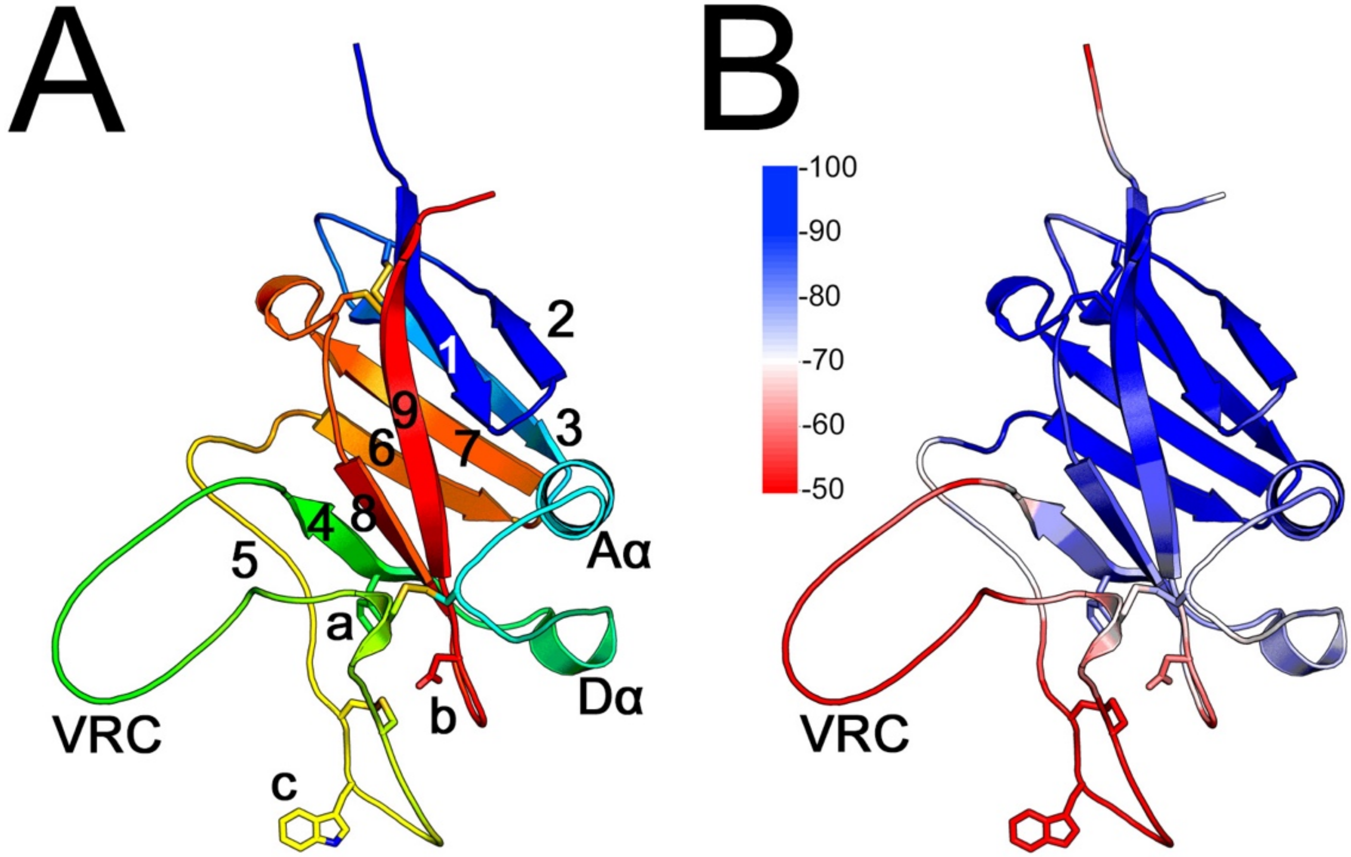


**Supplementary Figure 2** – HTLV-1 RBD model. (A) Model represented as for the RBD-C models in Figure 2. Residues corresponding to residues a-c of the conserved tetrad are shown as sticks. (B) pLDDT scores along the RBD model as shown in panel A. The scale to the left of the model indicates pLDDT scores.

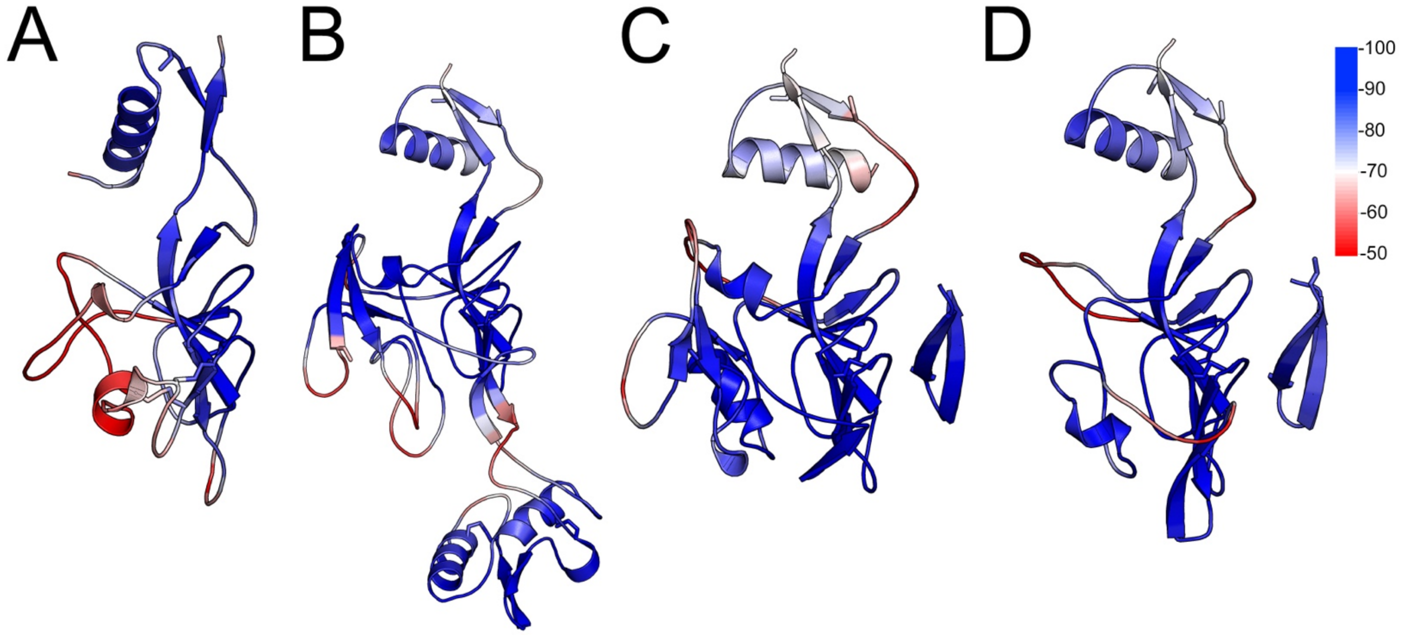


**Supplementary Figure 3** – pLDDT scores along PD section of the SU models shown in Figures 3A-D. The scale to the right of the model in panel D indicates pLDDT scores.

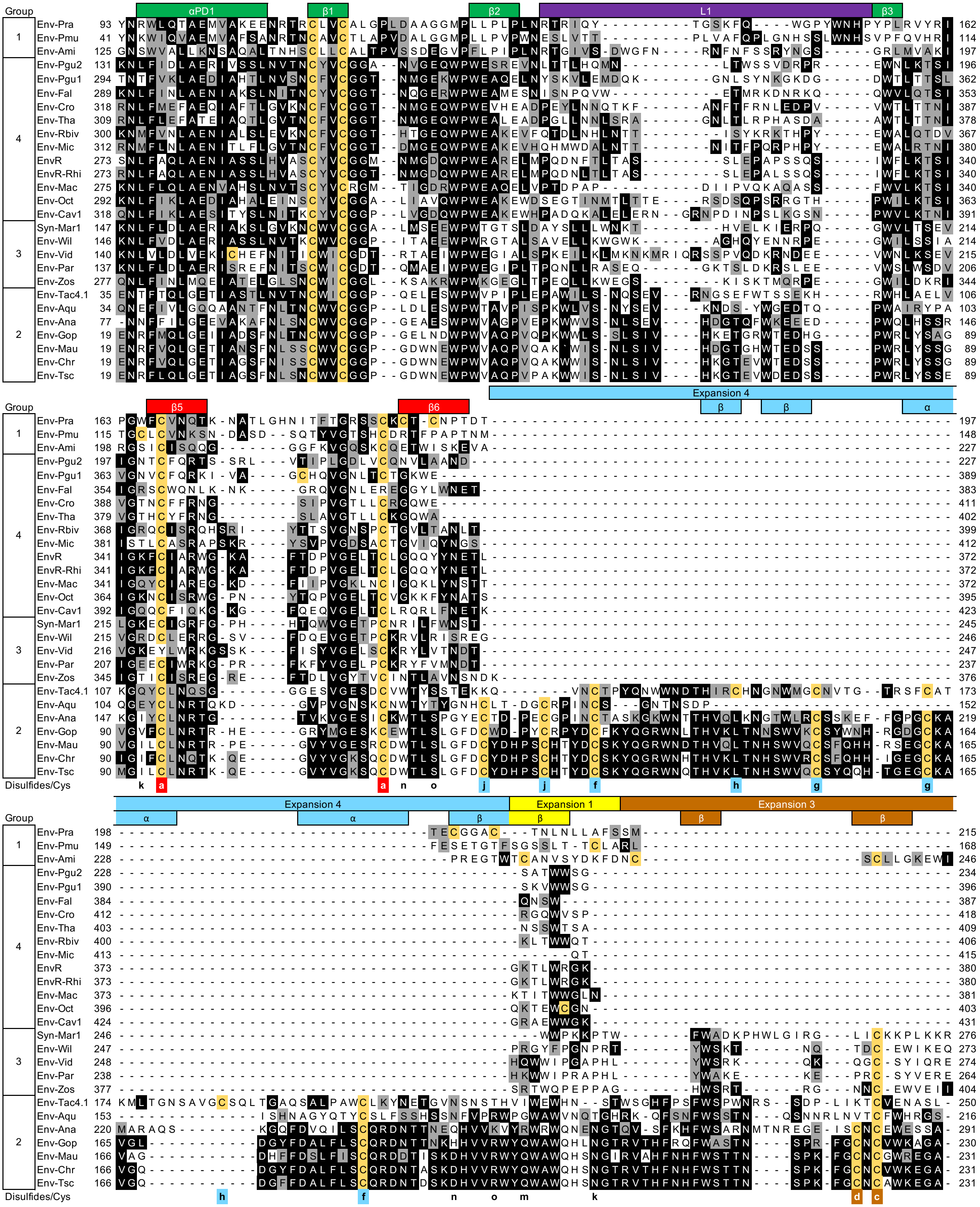

Group
αPD1
β1
β2
L1
β3
β5
β6
Expansion 4
Expansion 1
Expansion 3
β
α
Env-Pra
Env-Pmu
Env-Ami
Env-Pgu2
Env-Pgu1
Env-Fal
Env-Cro
Env-Tha
Env-Rbiv
Env-Mic
EnvR
EnvR-Rhi
Env-Mac
Env-Oct
Env-Cav1
Syn-Mar1
Env-Wil
Env-Vid
Env-Par
Env-Zos
Env-Tac4.1
Env-Aqu
Env-Ana
Env-Gop
Env-Mau
Env-Chr
Env-Tsc
1
2
3
4
Disulfides/Cys

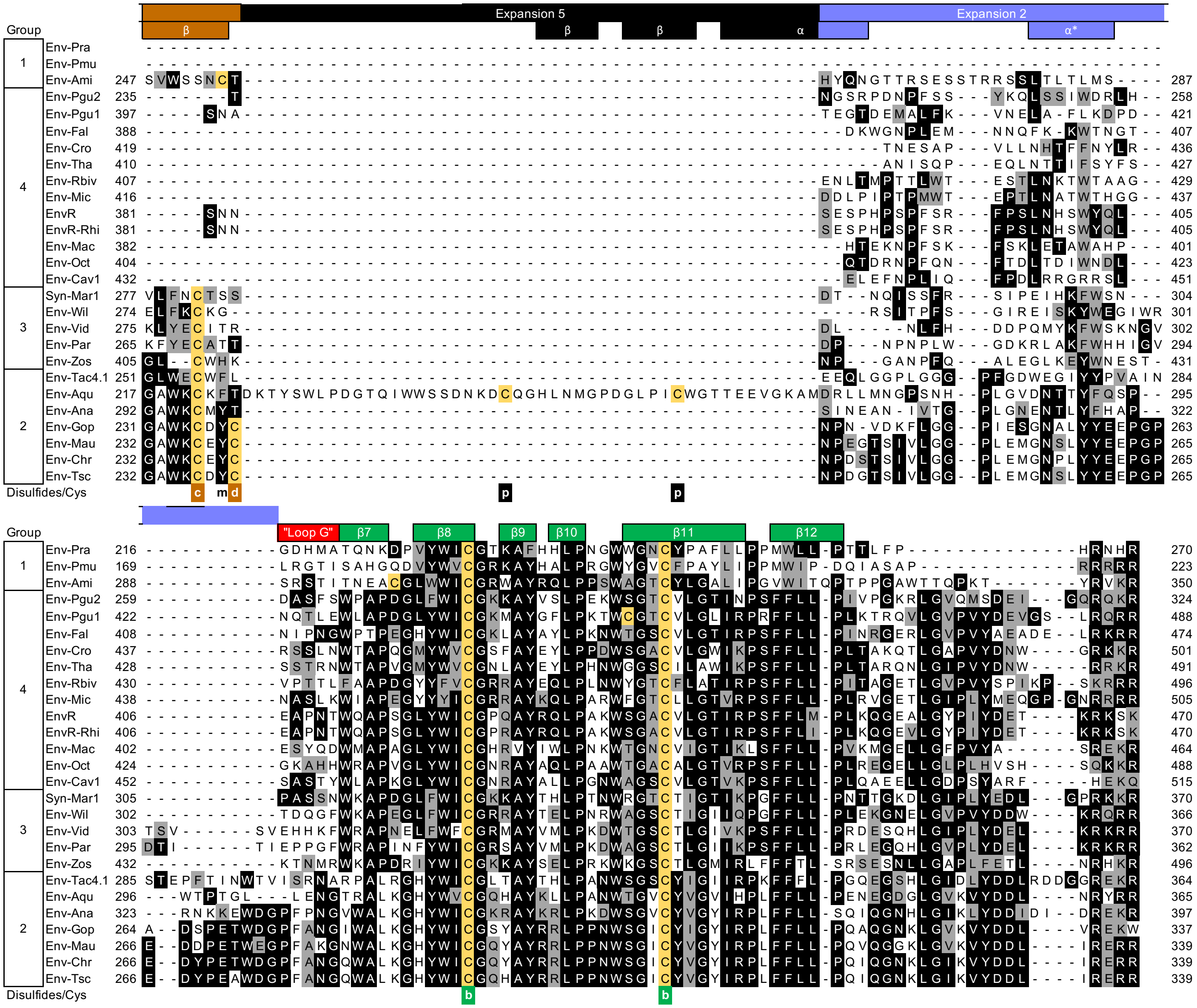

**Supplementary Figure 4** – Alignment of PD region sequences of the EnvR supergroup. Residues in black and gray are identical or similar (BLOSUM-62 matrix) to the normalized consensus sequence. The normalized consensus sequence was computed by obtaining the consensus for each group and then deriving a consensus of the 4 consensus sequences. Cysteine residues are highlighted in yellow and disulfides observed in the models are labeled with letters below the alignment. Secondary structure and expansion regions are indicated above the alignment. Sections of the sequences included in the expansion regions were excluded from multiple sequence alignments used to generate the phylogenetic tree in Figure 9. Residue numbering shown from the initiation codon of Env. The last residue shown corresponds to the last SU residue.

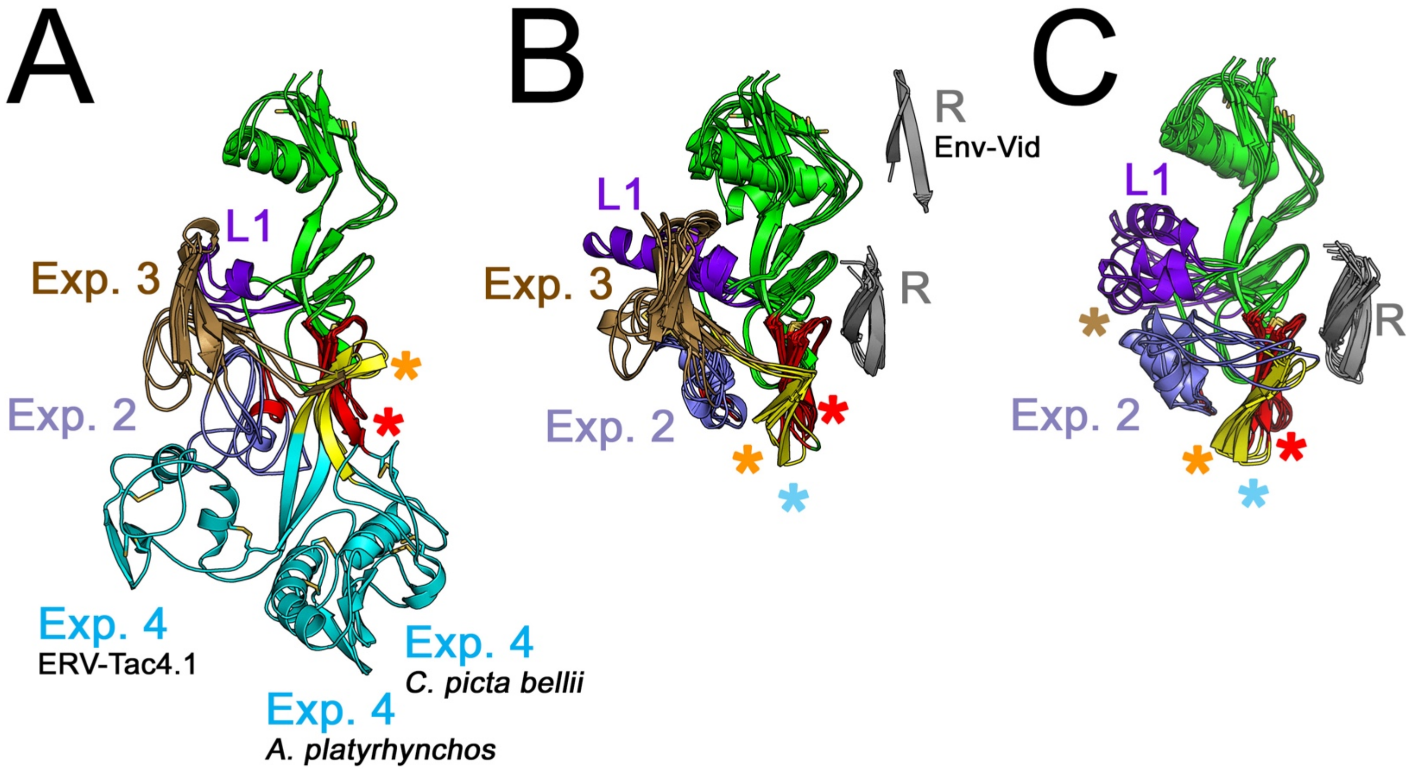


**Supplementary Figure 5** – Superposition of models of the PD region of SU in the EnvR Supergroup. The models in panels A-C correspond to the same groups as in Figure 5B-D. Labeling is the same as for Figure 5.

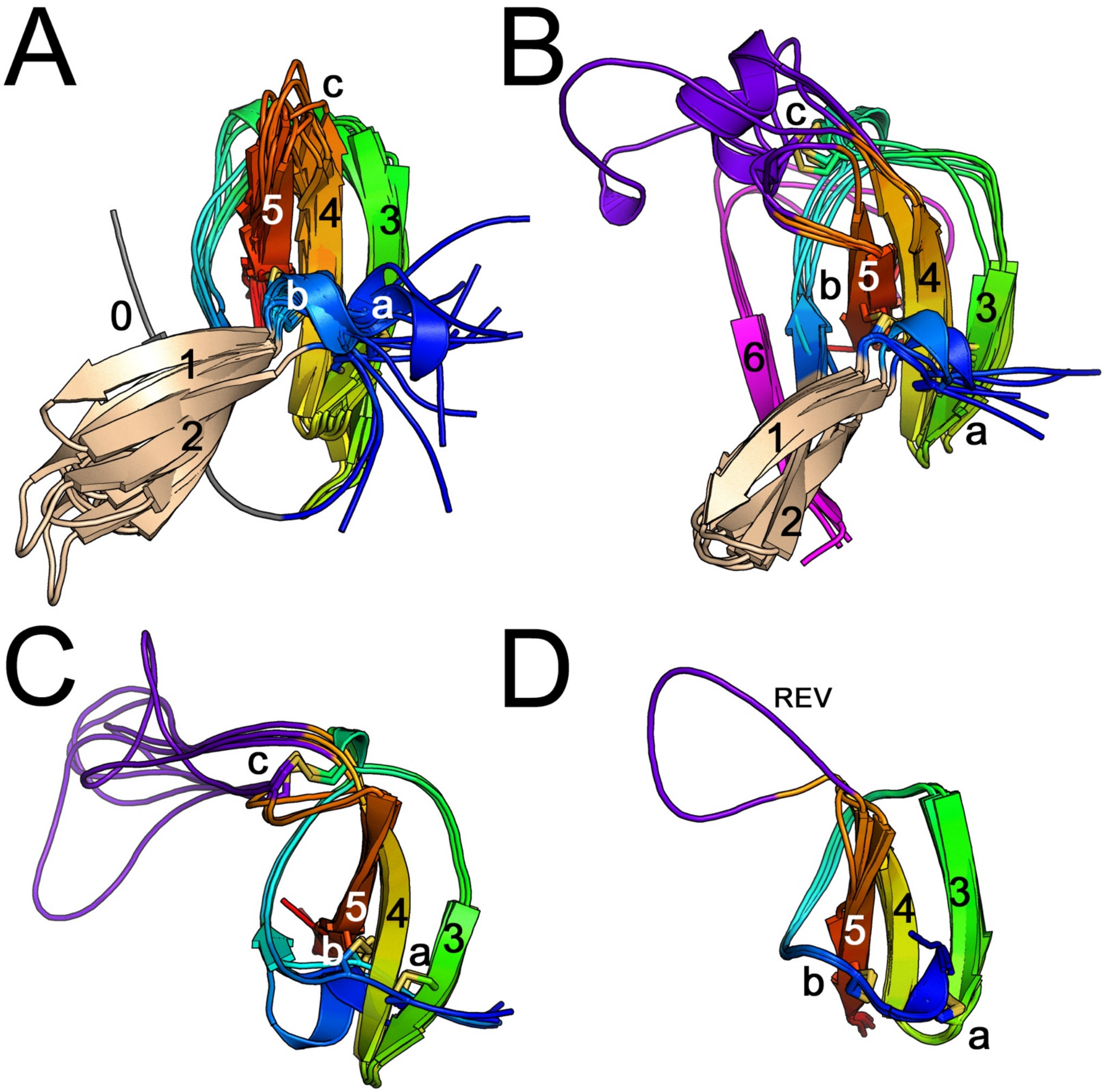


**Supplementary Figure 6** – Superposition of RBD-R models. RBD-R1 from EnvR (A) group 4 and (B) group 3, and (C) second and (D) first repeats of RBD-R2 models. The models are labeled as in Figure 6A, B, C and E. The loop expansion of the first domain of the REV RBD-R2 is indicated.

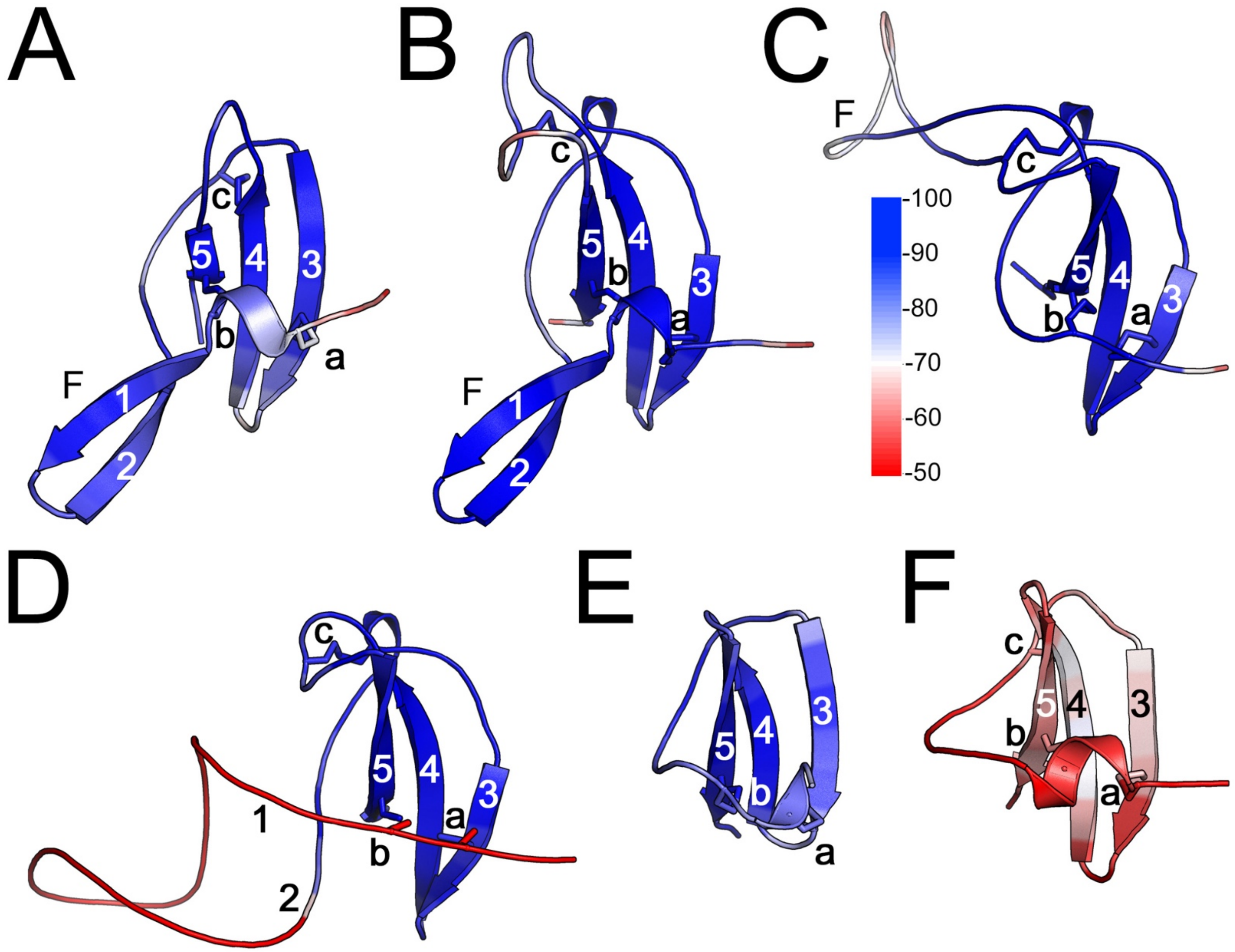


**Supplementary Figure 7** – pLDDT scores along the RBD-R sections of SU models as shown in Figures 6A-F. The scale to the right of the model in panel D indicates pLDDT scores.

```
Env-Mja         ---------------------------MKKNFQYQHPQKELLQSMATNKLHHWNEASWIV
Env-Mpe         -------------------------------------------MSLNRFRHWSED---P
Env-Per         MKRKAPDTTVLLEESFKKMRITGGQERTVRTRSATLPTRGQMKTLSQEAEGCLKDNGQET
Env-Dbi         ------------------------------------------------------------
Env-Ory         ----------------------------------------------------------MP

                           [SU/RBD starts
Env-Mja         KFLLLILVITFIHAHTAQACTITASVSSPPYLGRGVVLQCVSTCS-TPVSPIVWKKDSKE
Env-Mpe         YMIMKFFILISIHIHITQACTITASISSPPYFGRGVVLHCISTCS-TPVGPMIWKKDGNE
Env-Per         NVMSLFVMMMALLHSTSASCSVQASVWGVPMLGEPLYLNCTVICNNSEAGSVHWYKDGVE
Env-Dbi         ------IFLLCLLCCAALPCLLQTYTPTNVSVGQPLTLHCWTNCS-TPVGPITWTMNGRE
Env-Ory         LGTLLTIGIFLKTNESCHLSTRQASVTVSVGTPVILNCSCVYPNGSGMAGFVQWVRKNQT
                      . :          .   :           :   *    .   .. : *  ..

Env-Mja         IYRQKYGGKLGWNNATDQSKRDNTRYDIYKTVLTRNDIGNYSYEKYQKRSPDILYKSSYV
Env-Mpe         IYRQKDGGKLGWDNATDQSKRDNTRY-IYKTVLTRNDIGNYSCEKFQKGSPDQIYKSSYV
Env-Per         IFSQGELG-GAWTNISDQSERDNVQFGIVKLMASSSDEGIYNCSKIQKGSPQDNVLIGKN
Env-Dbi         IWSQKHQAISRYKPISDQSRRTSYNFSIALSSVTLRDGGWYSCIKWKKGNPDSPFLKETR
Env-Ory         IFRQKNITHWQWRNVYDQSKREIFNFSIAKWAMLS-DAGVYSCQFWKKNSPD--VLNGTV
                *: *       :    ***.*   .: *        * * *.    :* .*:

               RBD ends]Linker [PD/D-domain (CXXC-like)
Env-Mja         YLDISLSPEREFSSPNMWLQLASRRLNKSSFCVARDSSISGLLKENLIGIGLPFSVFTNI
Env-Mpe         YLNISLTPENEYSSPNMWLQLANQGLNKTSFCVAGDSSISRILRKNLIGTGLTFSGFTNI
Env-Per         VTVQLAPDSELTSSPNLWLQLAVDTMKKNSFCIKEGTSVQNILTQNLVGLAVSLDGYGNI
Env-Dbi         FISILPNATTPLKNTNLWLHLAHTVLNETAFCLRNSNNPSGLLQANLLGLGVTLP-----
Env-Ory         IVTLSVNASFSTEFGNFWLHLAATTN-TTTFCVNTADSPSHLLANNLTGIRSSREEFTSI
                            .  *:**:**      .:**:    . . :*  ** *   .

Env-Mja         SGINITHSPKVLTCELQLNLPVSTTIVSECICINCTHIPS-YCNSTIRVKWPFGYRLPNG
Env-Mpe         SGINITLAPKILTQEIQLNLPVSTTIVSSCICINCTNIPS-YCNSTIHVKWPYGYRLPNG
Env-Per         TQDEISQQ-EDEILENELTLDLEFIFHHECLCFDCEHHDLSNCPSVTRLSLPRGKRLGTG
Env-Dbi         ----NVLHAKVTIPPSVVTMPISGVTSTTCYCINCTYIPF-YCNKTIPVSIPTPSRLPAG
Env-Ory         TG-----------------LSITSLEIDNVTCLLFTVPWQLNRTSYREFHPLNGTRAPKG
                                   : :         *:           .   .      *   *

                                             SU ends][TM starts
Env-Mja         WVFLCNNKTYQAFSSHYQGVCGVRRILPALIDHPGYKHR--QISRALSPDCDDNLKLLDP
Env-Mpe         WVFLCDNKTYQAFSSHYQSICGIGCILPALIDHPGFKHR--RVTRALPPDCDDNLKLLDS
Env-Per         WSFFCDGKAYRALKKNYSGLCYVGRMVPLLLSKLTHQNKGHRKRRDLPPDCDDNLVLLNP
Env-Dbi         YFWLCDNAAYSHIPTDTSVACGIGPIVPLIAAFPTKNSR---VKRALTADCDPHVNLLPI
Env-Ory         WVFLCSQYTFTSLAPDDFRNCSLGRVAPTLASVFPKTRR----RRSPPENCDDTVLLSGE
                : ::*.  ::  :  .    * :  : * :        :     *  . :**  : *

Env-Mja         IVVAVAAALILPVPGLVVGMQNEISKMACAYSKTTNLTASVLSELNQELGEVRVAVLQDR
Env-Mpe         VAVAVTAAFALPVPGLVAGMQKEVSKLACAYSKTADLTASILSELNQEPGEVRVAVLQNR
Env-Per         ADMAVAAAFVLPVPGLVVGIQKEVSKLACVFNKVTNLTAVILEEINQEMKELRTGVLQNR
Env-Dbi         STVATLAALLLPDPGLIVDNHRQIEHLSCLLTHVMNDTVRAFLAISQELQELRRETLIQR
Env-Ory         DTVFSMAVSLVGVPGLAYGITRGMRELACLVTETVNLTAAHCPM----------------
                  :   *.  :  ***  .  . : .::*  ... : *.

Env-Mja         ATIDSYLLLKEHMGCEQFPGMCCFNLSDFSQTIHNHIDNIHHIIDKFSQMPELPKWFSWF
Env-Mpe         AAID-YLLLKEHLGCEQFPGMCCFNISDFSQTIQNQLDDIHHIIDKLSQLPGLPDWFSWF
Env-Per         AAID-YLLLKNHLGCEQFEGMCCFDITDNSQKIQSQLEQLKREMAKMKTSDDWG-FPNWF
Env-Dbi         AALD-YLFLLHQTPCTKIPGMCCFNVSDQTNVIFNAVHDLQDQVNNISYSKGLFDWLPWG
Env-Ory         ------------------------------------------------------------


Env-Mja         HWIWPVIVGLFLLCICLPVLIAWMRHIVTSLLKPMHAYATFLEVMSKK------------
Env-Mpe         HWLWPILVGLLLLCICLPVLVTWVHHLIAGLIKPIHAYATFQEVMAKKIIIACYFLESPC
Env-Per         SWLWPFMSPILAIMGVILLAPLLLQCMSSSIQRLLAKS----------------------
Env-Dbi         LGPLLRDSFIIIVVIGFWIIGCALCTCVRSSLSVRMVSYG--------------------
Env-Ory         ------------------------------------------------------------


Env-Mja         ------------
Env-Mpe         DRSLYIFEFRSN
Env-Per         ------------
Env-Dbi         ------------
Env-Ory         ------------
```

**Supplementary Figure 8** – CLUSTALW alignment of Alpha-like Env sequences with an RBD-C SIRP-like. Main sequence features including the CXXC-like motif in SU, cysteine residues in the fusion peptide of TM and CXCC motif are shown.

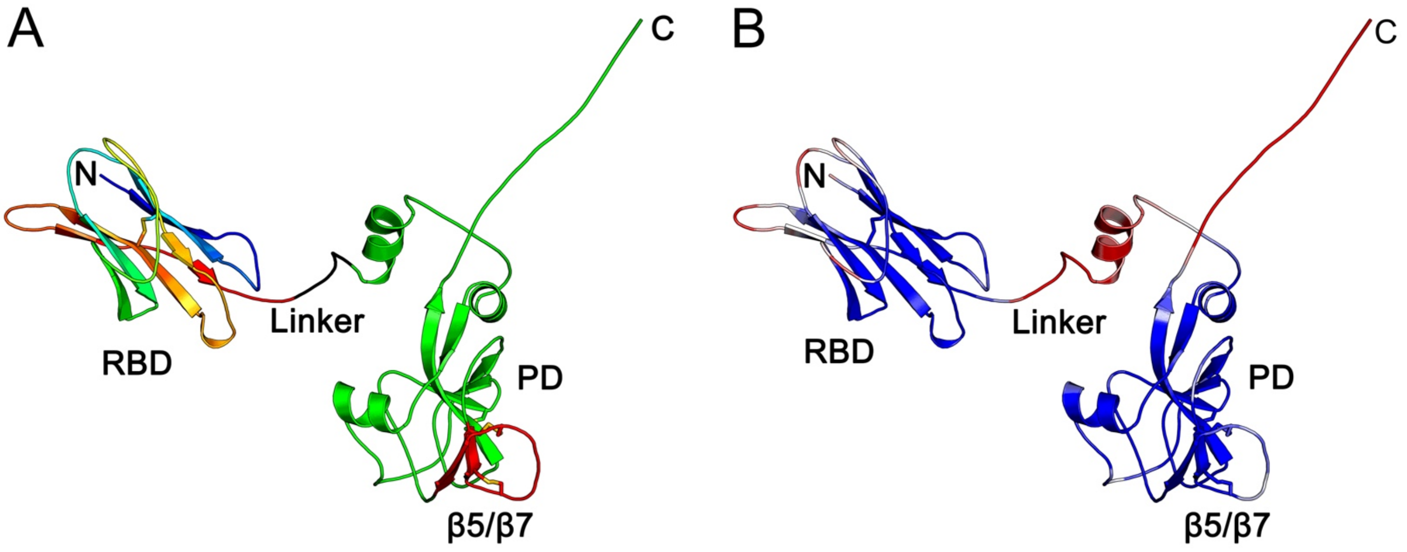


**Supplementary Figure 9** – Env-Per SU model showing the RBD and PD sections of the model. Panel B shows pLDDT scores along the model chain. The pLDDT score scale is the same as in Supplementary Figure 7C.

```
Provirus flanking regions, JAJIAZ010000042.1 (1st seq.): CAGCTTCTAGTTGGTCTGCCCTGG [provirus] CCTGGTCATCTGGAAAGGGAGAA
Provirus flanking regions, JAJIAY010000041.1 (2nd seq.): TGTGACCCAAGAAATTTATATCCC [provirus] ATCCCCCAAATTGTTATTCAAAT
Provirus flanking regions, JANTPW010000004.1 (3rd seq.): TGTGACCCAAGAAATTTATATCCC (provirus) ATCCCCCAAATTGTTATTCAAAT

   1 TGTCTTGACA TGAGTGCAGC CATGTTTGGC CGCTCCATTG ACCTACAGCC ACCAGCACAA AAAGAAATAT AAAAGCTGCT TTCTGCGTGG TGGGAACTGG
     TGTCTTGACA TGAGTGCAGC CATGTTTGGC CGCTCCGTTG ACCTACAGCC ACCAGCACAA AAAGAAATAT AAAAGCTGCT TTCTGCGTGG TGGGAACTGG
     TGTCTTGACA TGAGTGCAGC CATGTTTGGC CGCTCCGTTG ACCTACAGCC ACCAGCACAA AAAGAAATAT AGAAGCTGCT TTCTGCGTGG TGGGAACTGG

 101 CCCTTCTCTC ACGGAGGGAG TTGCAGGTTG TAACATTTCA AGGCTCTTGG AGCGTCGTAG CACGGGATGG ACTGGCCATC TATGCCGGGC TGGGACGATA
     CCCTTCTCTC ACGGAGGGAG TTGCAGGTTG TAACATTTCA AGGCTCTTGG AGCGTCGTAG CACGGGATGG ACTGGCCATC TATGCCGGGC TGGGACGATA
     CCCTTCTCTC ACGGAGGGAG TTGCAGGTTG TAACATTTCA AGGCTCTTGG AGCGTCGTAG CACGGGATGG ACTGGCCATC TATGCCGGGC TGGGACGATA

 201 ACCAAAGAGA ACAATGCACG TACACAACTA CTGGGCTGGG ACGTTAGCCA GGGGGCTCAG TGCACGTACG CCACTGATAA CCATTTTGTG CATGGGAACA
     ACCAAAGAGA ACAATGCACG TACACAACTA CTGGGCTGGG ACGTTAGCCA GGGGGCTCAG TGCACGTACG CCACTGATAA CCATTTTGTG CATGGGAACA
     ACCAAAGAGA ACAATGCACG TACACAACTA CTGGGCTGGG ACGTTAGCCA GGCGGCTCAG TGCACGTACG CCACTGATAA CCATTTTGTG CATGGGAACA

 301 CTAAATGCTT GCTCTGCAAA CTCTGTAATT GCTTACTCTG AAAATTACTG ATGGTATAAA AACTGCTGGA AACTGAGACC GAGAGAGAGT TCCACCTGTG
     CTAAATGCTT GCTCTGCAAA CTCTGTAATT GCTTACTCTG AAAATTACTG ATGGTATAAA AACTGCTGGA AACTGAGACC CAGAGAGAGT TCCACCTGTG
     CTGAATGCTT GCTCTGCAAA CTCTGTAATT GCTTACTCTG AAAATTACTG ATGGTATAAA AACTGCTGGA AACTGAGACC CAGAGAGAGT TCCACCTGTG
                                                              ^TATA Box

 401 GAAGGGACAC CTCGCCCAGG ACGTGTGATC CTTGCCCAGC CGCGTCGATT GACAGGGCTC TCCTGGCAGT GGGACGTGGA TGTTGTGAGT AACTGATTTG
     GAAGGGACAC CTCGCCCAGG ACGTGTGATC CTTGCCCAGC CGCGTCAATT GACAGGGCTC TCCTGGCAGT GGGACGTGGA TGTTGTGAGT AACTGATTTA
     GAAGGGACAC CTCGCCCAGG ACGTGTGATC CTTGCCCAGC CGCGTCAATT GACAGGGCTC TCCTGGCAGT GGGACGTGGA TGTTGTGAGT AACTGATTTA

 501 ATGTGAAGCA ATTGATTTGA TATGAAGTAA TTGATTTGAT GTGTTCTCTT TTTGGTCAAC CTGGTGTTTC TCTTTCTGGT TGATTTGTGT CATTTCTCTT
     ATGTGAAGTA ATTGATTTGA TGTGTAGTAA GTGATTTGAT GTGTTCTTTT TTTGGTCAAC CTGGTGTTTC TCTTTCTGGT TGATTTGTGT CATTTCTCTT
     ATGTGAAGTA ATTGATTTGA TGTGTAGTAA GTGATTTGAT GTGTTCTTTT TTTGGTCAAC CTGGTGTTTC TCTTTCTGGT TGATTTGTGT CATTTCTCTT

 601 CAGCCGATGT GGGTGCTTTC CCTTTCCAGT CGGGCTGATG TTTTTTCCCT CTTAATATTT GACCTATTTG GATCTGTTTG CAGACTGTCG CCTTTACTAT
     CAGCCGATGT GGGTGCTTTC CCTTTCCAGT CGGGCTGATG TTTTTTCCCT CTTAATATTT GACCTATTTG GATCTGTTTG CAGACTGTCG CCTTTACTAT
     CAGCCGATGT GGGTGCTTTC CCTTTCCAGT CGGGCTGATG TTTTTTCCCT CTTAATATTT GACCTATTTG GATCTGTTTG CAGACTGTCG CCTTTACTAT

 701 CCCCTATATA ATAAAATATA CCTTCAGTCC ATTTGTTTGG AGTGGAAAGT GTCTTTTACG TCTCCGATCG AATCCCCGAA CCTCTCGTAA CATAATTGGC
     CCTCTATATA ATAAAATATA CCTTCAGTCC ATCTGTTTGG AGTGGAAAGT GTCTTTTACG TCTCCGATCG AATCCCCGAA CCTCTCATAA CATAATTGGC
     CCTCTATATA ATAAAATATA CCTTCAGTCC ATCTGTTTGG AGTGGAAAGT GTCTTTTACG TCTCCGATCG AATCCCCGAA CCTCTCATAA CATAATTGGC
               ^Poly(A)                                                                          *       ^End of LTR
                                                                                    PBS tRNA-Ser-CGA 1-1^

 801 GCTGTGAGCA GGATTCGATT AAGGAGATGC CTTTCCTATT CAAACGAAAG GTAACTGAAG CCAAGGTGGA AGAAATTCCA GGATGGGAGG ACCCTCCCTT
     GCTGTGAGCA GGATTCGATT AAGGAGATGC CTTTCCTATT CAAACGAAAG GTAACTGAAG TCAAGGTAGA AGAAATTCCA GGATGGGAGG ACCCTCCCTT
     GCTGTGAGCA GGATTCGATT AAGGAGATGC CTTTCCTATT CAAACGAAAG GTAACTGAAG TCAAGGTAGA AGAAATTCCA GGATGGGAGG ACCCTCCCTT
   1                               M  P   F  L  F   K  R  K   V  T  E  A   K  V  E   E  I  P   G  W  E  D   P  P  F
                                 ^gag open reading frame               ^Splice donor consensus

 901 TTGCACTCTG GCTGGGGAGT GGGCTCGTAG GTCAGGGCTA TGTGAAGCTT GGGATGCTCG TCTGCGACGA TTTGAACCGT TGGATGTGAC CAAAGCTTTG
     TTGCACTCTG GCTGGGGAGT GGGCTCGTAG ATCAGGGTTA TGTGAAGGTT GGGATGCTCG TCTGCGACGA TTTGAACCGT TGGATGTGAC CAAAGCTTTG
     TTGCACTCTG GCTGGGGAGT GGGCTCGTAG GTCAGGGTTA TGTGAAGGTT GGGATGCTCG TCTGCGACGA TTTGAACCGT TGGATGTGAC CAAAGCTTTG
  26  C  T  L   A  G  E  W   A  R  R   S  G  L   C  E  A  W   D  A  R   L  R  R   F  E  P  L   D  V  T   K  A  L

1001 CAGACGGTGG AAAGAAAAGC GAAAGATGAT GTTTCTTTTT TGGCACGGCG AGGTTGGTGT TTGTTGACTG CTTATCGCCG GGCTATAGAA GATAGGGATA
     CAGACGGTGG AAAGAAAAGC GAAAGATGAT GTTTCTTTTT TGGCACGGCG AGGTTGGTGT TTGTTGACTG CTTATCGCCG GGCTATAGAA GATAGGGATA
     CAGACGGTGG AAAGAAAAGC GAAAGATGAT GTTTCTTTTT TGGCACGGCG AGGTTGGTGT TTGTTGACTG CTTATCGCCG GGCTATAGAA GATAGGGATA
  59 Q  T  V  E   R  K  A   K  D  D   V  S  F  L   A  R  R   G  W  C   L  L  T  A   Y  R  R   A  I  E   D  R  D  K

1101 AGATACAGAC AGAGAAGGCA CTCTCGGAAA AAGAAAATTT AGATTTGCAG TCTCGCTTGG CTATGATTCA ATGTCAATAT CATGCACTGG GAGATCAGGC
     AGATACAGAC AGAGAAGGCA CTCTCGGAAA AAGAAAATTT AGATTTGCAG TCTCGCTTGG CTATGATTCA ATGTCAATAT CTTGCACTGG GAGATCAGGC
     AGATACAGAC AGAGAAGGCA CTCTCGGAAA AAGAAAATTT AGATTTGCAG TCTCGCTTGG CTATGATTCA ATGTCAATAT CTTGCACTGG GAGATCAGGC
  93  I  Q  T   E  K  A   L  S  E  K   E  N  L   D  L  Q   S  R  L  A   M  I  Q   C  Q  Y   H  A  L  G   D  Q  A

1201 TCGTAACTAC CAAGCTATAG CAGAAAAGGC TGCTGTAAGA GTTGCCAAAT ATAAATATAG AAAGAGGAGA GGAAAGGTAA ACGAACGAAA AGTTCATATG
     TCGTAACTAC CAAACTATAG CAGAAAAGGC TGCTGTAAGA GTTGCCAAAT ATAAATACAG AAAGAGGAGA GGAAAGGTAA ACGAACGAAA AGTTCATATG
     TCGTAACTAC CAAACTATAG CAGAAAAGGC TGCTGTAAGA GTTGCCAAAT ATAAATACAG AAAGAGGAGA GGAAAGGTAA ACGAACGAAA AGTTCATATG
 126  R  N  Y   Q  A  I  A   E  K  A   A  V  R   V  A  K  Y   K  Y  R   K  R  R   G  K  V  N   E  R  K   V  H  M

1301 GCCATAGCAT CGGCCGGTCC TAAGTGGGAC CCGGACACAT GGGATGGTGA TATCTGGGAT GATGATTCTG AAACTGATTA TGAGGAAGAG GAAAAACTGC
     GCCATAGCAT CGGCCGGTCC TAAGTGGGAC CCGGACACAT GGGATGGTGA TATCTGGGAT GATGATTCTG AAACTGATTA TGAGGAAGAG GAAAAACCGC
     GCCATAGCAT CGGCCGGTCC TAAGTGGGAC CCGGACACAT GGGATGGTGA TATCTGGGAT GATGATTCTG AAACTGATTA TGAGGAAGAG GAAAAACCGC
 159 A  I  A  S   A  G  P   K  W  D   P  D  T  W   D  G  D   I  W  D   D  D  S  E   T  D  Y   E  E  E   E  K  L  P

1401 CCAAACACAC AGAGGCACGT CCTCTTCGCC GTCGGCGGAT GGAAATGGAT GGTAATCATC CTGTAAGGAA AGATGTGATT GAGGATTTTA ACCAACAAGA
     CCAAACACAC AGAGGCACGT CCTCTTCGCC GTCGGCGGAT GGAAATGGAT GGTAATCATC CTGTAAGGAA AGATGTGATT GAGGATTTTA ACCAACAAGA
     CCAAACACAC AGAGGCACGT CCTCTTCGCC GTCGGCGGAT GGAAATGGAT GGTAATCATC CTGTAAGGAA AGATGTGATT GAGGATTTTA ACCAACAAGA
 193  K  H  T   E  A  R   P  L  R  R   R  R  M   E  M  D   G  N  H  P   V  R  K   D  V  I   E  D  F  N   Q  Q  E
```

```
1501 AATTTCTGAT ATTTTAGCCC GGGCTACACA AAAGCCTGGA GAAGCCATTA TAACCTGGCT TGTTCGCCTA TATGATATGG GCGCTACAGG AATTAATCTG
     AATTTCTGAT ATTTTAGCCC GGGCTACACA AAAGCCTGGA GAAGCCATTA TAACCTGGCT TCTTCGCCTA TATGATATGG GCGCTACAGG AATTAATCTG
     AATTTCTGAT ATTTTAGCCC GGGCTACACA AAAGCCTGGA GAAGCCATTG TAACCTGGCT TGTTCGCCTA TATGATATGG GCGCTACAGG AATTAATCTG
 226  I  S  D   I  L  A  R   A  T  Q   K  P  G   E  A  I  I   T  W  L   V  R  L   Y  D  M  G   A  T  G   I  N  L

1601 GATGCAGGAG ATGCTTTTAA ATTTGTAGTT TTGTCCAATG ATCCAGTAAT ACGGGGAGCT TTCAGAGACT GGAGCATGAA TGCTCACCCC CCAAATGATC
     GATGCAGGAG ATGCTTTTAA ATTTGTAGTT TTGTCCAATG ATCCAGTAAT ACGGGGAGCT TTCAGAGACT GGAGCATGAA TGCTCACCCC CCAAATGATC
     GATGCAGGAG ATGCTTTTAA ATTTGTAGTT TTGTCCAATG ATCCAGTAAT ACG-GGAGCT TTCAGAGACT GGAGCATGAA TGCTCACCC- CCAAATGATC
 259 D  A  G  D   A  F  K   F  V  V   L  S  N  D   P  V  I   R  G  A   F  R  D  W   S  M  N   A  H  P   P  N  D  R

1701 GTGGACAAGA CACTGATGGT ACAACTTTAT TGGCTCTTGC AGTAAAAGGA GCTCAAGATA AATACCCCAC GGATGATATG TGGCCAGACA CTAATAGACC
     GTGGACAAGA CACTGATGGT ACAACTTTAT TGGCTCTTGC AGTAAAAGGA GCTCAAGATA AATACCCCAC GGATGATATG TGGCCAGACA CTAATAGACC
     GTGGACAAGA CACTGATGGT ACAACTTTAT TGGCTCTTGC AGTAAAAGGA GCTCAAGATA AATACCCCAC GGATGATATG TGGCCAGACA CTAATAGACC
 293  G  Q  D   T  D  G   T  T  L  L   A  L  A   V  K  G   A  Q  D  K   Y  P  T   D  D  M   W  P  D  T   N  R  P

1801 ATGGTATACT TTAAAAGATT GTGTCCAAAA GATGAGAGAA GAAACCATGA AACGGGCTGT GTTTCTGGGG TGTGGTGAAG CTTTAAATGA AATGACTTTA
     ATGGTATACT TTAAAAGATT GTGTCCAAAA GATGAGAGAA GAAACCATGA AACGGGCTGT GTTTCTGGGG TGTGGTGAAG CTTTAAATGA AATGACTTTA
     ATGGTATACT TTAAAAGATT GTGTCCAAAA GATGAGAGAA GAAACCATGA AACGGGCTGT GTTTCTGGGG TGTGGTGAAG CTTTAAATGA AATGACTTTA
 326  W  Y  T   L  K  D  C   V  Q  K   M  R  E   E  T  M  K   R  A  V   F  L  G   C  G  E  A   L  N  E   M  T  L

1901 AGTGTCCAGG TGCGAAACAA GTTGATAAAA ACTGCACCTC CAGCATACAA ACAAATAATT ACAACCCTTC TGATGAATGA AACTGAAAGC CCTATATCAA
     AGTGTCCAGG TGCGAAACAA GTTGATAAAA ACTGCACCTC CAGCATACAA ACAAATAATT ACAACCCTTC TGATGAATGA AACTGAAAGC CCTATATCAA
     AGTGTCCAGG TGCGAAACAA GTTGATAAAA ACTGCACCTC CAGCATACAA ACAAATAATT ACAACCCTTC TGATGAATGA AACTGAAAGC CCTATATCAA
 359 S  V  Q  V   R  N  K   L  I  K   T  A  P  P   A  Y  K   Q  I  I   T  T  L  L   M  N  E   T  E  S   P  I  S  T

2001 CACTTGCAGA TAAGATTATG CAGCTAGCAG ACTTGGGAGA CTGGCAGACA ACTGGAAACT CTCCAAGAAG GGAAAA-AAGA GAAACATTCT TTAAAAGAGA
     CACTTGCAGA TAAGATTATG CAGCTAGCAG ACTTGGGAGA CTGGCAGACA ACTGAAAATT CTCCAAGAAG AGAAAAAAGA GAAACATTCT TTAAAAGAGA
     CACTTGCAGA TAAGATTATG CAGCTAGCAG ACTTGGGAGA CTGGCAGACA ACTGAAAATT CTCCAAGAAG AGAAAA-AAGA GAAACATTCT TTAAAAGAGA
 393   L  A  D   K  I  M   Q  L  A  D   L  G  D   W  Q  T   T  G  N  S   P  R  R   E  K   R   E  T  F  F   K  R  D

2101 TAAAATGACT CGAGGTGATA TGTTTTTGGC TCTACTCCAG GCAGGAGTAC CTAAAGACCA GATTGATGGA GTTGACACTA AAGAATTATG GAAACTATAT
     TAAAATGACT CGAGGTGATA TGTTTTTGGC TCTACTCCAG GCAGGAGTAC CTAAAGACCA GATTGATGGA GTTGACACTA GAGAATTATG GAAATTATAT
     TAAAATGACT CGAGGTGATA TGTTTTTGGC TCTACTCCAG GCAGGAGTAC CTAAAGACCA GATTGATGGA GTTGACACTA GAGAATTATG GAAATTATAT
 426  K  M  T   R  G  D  M   F  L  A   L  L  Q   A  G  V  P   K  D  Q   I  D  G   V  D  T  K   E  L  W   K  L  Y

2201 AAAGAAAAAG GGCTGAATGT AAAACAAGTG AACAGAAGGG ACAGTATAGA GAATGCTTCC AACCCAGTGC CTGCTGCACC TAGTGCACCA CCTCCTCCCT
     AAAGAAAAAG GGCTGAATGT AAAACAAGTG AACAGAAGGG ACAGTATAGA GAATGCTTCC AACCCAGTGC CTGCTGCACC TAGTGCACCA CCTCCTCCCT
     AAAGAAAAAG GGCTGAATGT AAAACAAGTG AACAGAAGGG ACAGTATAGA GAATGCTTCC AACCCAGTGC CTGCTGCACC TAGTGCACCA CCTCCTCCCT
 459 K  E  K  G   L  N  V   K  Q  V   N  R  R  D   S  I  E   N  A  S   N  P  V  P   A  A  P   S  A  P   P  P  P  L

2301 TGGATCCTCT TCTTGCTGAC TTATGTGATT GAGGGGAAGG CCAAACCTCT GTATCGATTA GTGCCACTAT AAGAAAAGAC ATTAGACCAC ATGTACCAAT
     TGGATCCTCT TCTTGCTGAC TTATGTGATT GAGGGGAAGG CCAAACCTCT GTATCGATTA GTGCCACTAT AAGAAAAGAC ATTAGACCAC ATGTACCAAT
     TGGATCCTCT TCTTGCTGAC TTATGTGATT GAGGGGAAGG CCAAACCTCT GTATCGATTA GTGCCACTAT AAGAAAAGAC ATTAGACCAC ATGTACCAAT
 493   D  P  L   L  A  D   L  C  D  *   G  E  G   Q  T  S   V  S  I  S   A  T  I   R  K  D   I  R  P  H   V  P  I
                                        ^pol

2401 TAAAATATTT TGGAAGAATG GATCTACAAC TAGTGTGAAG GCACTTGTAG ATACAGGGGC GGAGGCCACC CTTATCTATG GGAATCCAAC TAAGTTTCAA
     TAAAATATTT TGGAAGAATG GATCTACAAC TAGTGTGAAG GCACTTGTAG ATACAGGGGC GGAGGCCACC CTTATCTATG GGAATCCAAC TAAGTTTCAA
     TAAAATATTT TGGAAGAATG GATCTACAAC TAGTGTGAAG GCACTTGTAG ATACAGGGGC GGAGGCCACC CTTATCTATG GGAATCCAAC TAAGTTTCAA
  24 K  I  F   W  K  N  G   S  T  T   S  V  K   A  L  V  D   T  G  A   E  A  T   L  I  Y  G   N  P  T   K  F  Q

2501 GGTACATCAA TTTTGATAAC TGGATTGGGA GGAAAAGAAA TCCAGGCAGT TACCACTAAT CTTTCTATGC AAATTGGGCA ACTACCCCGG CGCACATATC
     GGTACATCAA TTCTGATAAC TGGATTGGGA GGAAAAGAAA TCCAGGCAGT TACCACTAAT CTTTCTATGC AAATTGGACA ACTACCCCGG CGCACATATC
     GGTACATCAA TTCTGATAAC TGGATTGGGA GGAAAAGAAA TCCAGGCAGT TACCACTAAT CTTTCTATGC AAATTGGACA ACTACTCCGG CGCACATATC
  57 G  T  S  I   L  I  T   G  L  G   G  K  E  I   Q  A  V   T  T  N   L  S  M  Q   I  G  Q   L  P  R   R  T  Y  L

2601 TAGTTATGAT TGTACCAATA CCAGAATATA TAATTGGGAT TGATATTCTA AAAGGATTGA CATTAAATTT GGCTGATGGA CAATACCAAT TTGCTATTAG
     TAGTTATGAT TGTACCAATA CCAGAATATA TAATTGGGAT TGATATTCTA AAAGGATTGA CATTAAATTT GGCTGATGGA CAATACCAGT TTGCTATTAG
     TAGTTATGAT TGTACCAATA CCAGAATATA TAATTGGGAT TGATATTCTA AAAGGATTGA CATTAAATTT GGCTGATGGA CAATACCAGT TTGCTATTAG
  91   V  M  I   V  P  I   P  E  Y  I   I  G  I   D  I  L   K  G  L  T   L  N  L   A  D  G   Q  Y  Q  F   A  I  R

2701 AGCTTTGTCT TTTCATATTA ATGCTGTAAT TGTGGGATGT TTGCATCATG AAGTAGAAAT TCCACCTGCC ACGCAAGTGA TCAATCAAAG ACAGTATCGC
     AGCTTTGTCT TTCCATATTA ATGCTGTAAT TGTGGGATGT TTGCATCATG AAGTAGAAAT TCCACCTGCC ACGCAAGTGA TCAATCAAAG ACAGTATCGC
     AGCTTTGTCT TTCCATATTA ATGCTGTAAT TGTGGGATGT TTGCATCATG AAGTAGAAAT TCCACCTGCC ACGCAAGTGA TCAATCAAAG ACAGTATCGC
 124  A  L  S   F  H  I  N   A  V  I   V  G  C   L  H  H  E   V  E  I   P  P  A   T  Q  V  I   N  Q  R   Q  Y  R

2801 ATCCCTGGAG GACAACAAGA AATATCTGAG ACCATAGAAG ATTATTTGGC TGTAAAAGTC CTTCGCAGAG TAACTACAGC CTGGAATAAT CCTATCTGGC
     ATCCCTGGAG GACAACAAGA AATATCTGAG ACCATAGAAG ATTATTTGGC TGTAAAAGTC CTTCGCAGAG TAACTACAGC CTGGAATAAT CCTATCTGGC
     ATCCCTGGAG GACAACAAGA AATATCTGAG ACCATAGAAG ATTATTTGGC TGTAAAAGTC CTTCGCAGAG TAACTACAGC CTGGAATAAT CCTATCTGGC
 157 I  P  G  G   Q  Q  E   I  S  E   T  I  E  D   Y  L  A   V  K  V   L  R  R  V   T  T  A   W  N  N   P  I  W  P

2901 CTGTTAAAAA AGGTGATGGG ACATGGAGAA TGACTGTAGA TTACAGAGAG CTAAACAAAG TGACTCCGGC TATTCAAGCG GCCGTCCCTG ATTTGATTAC
     CTGTTAAAAA AGGTGATGGG ACATGGAGAA TGACTGTAGA TTACAGAGAG CTAAACAAAG TGACTCCGGC TATTCAAGCG GCCGTCCCTG ATTTGATTAC
     CTGTTAAAAA -GGTGATGGG ACATGGAGAA TGACTGTAGA TTACAGAGAG CTAAACAAAG TGACTCCGGC TATTCAAGCG GCCGTCCCTG ATTTGATTAC
 191   V  K  K   G  D  G   T  W  R  M   T  V  D   Y  R  E   L  N  K  V   T  P  A   I  Q  A   A  V  P  D   L  I  T
```

3001 TCTTATAGAA AAAGTACAAT GTTATCCAGG CACTTGGTAT GCAGTCATAG ACTTGGCTAA TGCCTTTTTC ACCATACCTA TACCTAAAAG TGTCCAAGAA
TCTTATAGAA AAAGTGCAAT GTTATCCAGG CACTTGGTAT GCAGTCATAG ACTTGGCTAA TGCCTTTTTC ACCATACCTA TACCTAAAAG TGTCCAAGAA
TCTTATAGAA AAAGTGCAAT GTTATCCAGG CACTTGGTAT GCAGTCATAG ACTTGGCTAA TGCCTTTTTC ACCATACCTA TACCTAAAAG TGTCCAAGAA
224 L I E K V Q C Y P G T W Y A V I D L A N A F F T I P I P K S V Q E

3101 CAGTTTGCTT TCACGTGGCT GGGACAACAA TATACCTTTA CTAGACTTCC TCAGGGATAT GTGCATAGTC CCACTATCTG CCATCGACAG GTTGCTGAAA
CAGTTTGCTT TCACGTGGCT GGGACAACAA TATACCTTTA CTAGACTTCC TCAGGGATAT GTGCATAGTC CCACTATCTG CCATCGACAG GTTGCTGAAA
CAGTTTGCTT TCACGTGGCT GGGACAACAA TATACCTTTA CTAGACTTCC TCAGGGATAT GTGCATAGTC CCACTATCTG CCATCGACAG GTTGCTGAAA
257 Q F A F T W L G Q Q Y T F T R L P Q G Y V H S P T I C H R Q V A E T

3201 CCTTGGCAAA GGTACCTGCT TCTGATGATG TCCAAATAGT ACATTATATA GATGATATTA TGATCCAAGG AAATGAAGAA GAAAAAGTAC AACAACAGTT
CCTTGGCAAA GGTACCTGCT TCTGATGATG TCCAAATAGT ACATTATATA GATGATATTA TGATCCAAGG AAATGAAGAA GAAAAAGTAC AACAACAGTT
CCTTGGCAAA GGTACCTGCT TCTGATGATG TCCAAATAGT ACATTATATA GATGATATTA TGATCCAAGG AAATGAAGAA GAAAAAGTAC AACAACAGTT
291 L A K V P A S D D V Q I V H Y I D D I M I Q G N E E E K V Q Q Q L

3301 AGAAAAAGTA ATGGACACAC TTGAGGAGGA TGGATGGAAA ACTAATCCAG CTAAAATTCA GGGCCCGAGT TCAAATGTTA AATTTTTGGG AGTACTTTGG
AGAAAAAGTA ATGGACATAC TTGAGGAGGA TGGATGGAAA ACTAATCCAG CTAAAATTCA GGGCCCGAGT TCAAATGTTA AATTTTTGGG AGTACTTTGG
AGAAAAAGTA ATGGACATAC TTGAGGAGGA TGGATGGAAA ACTAATCCAG CTAAAATTCA GGGCCCGAGT TCAAATGTTA AATTTTTGGG AGTACTTTGG
324 E K V M D T L E E D G W K T N P A K I Q G P S S N V K F L G V L W

3401 AATAATGGAA AACAAGAAAT TTTGCCTAAA GCTAGACAAA AGATATTAGA TTTTGCAGCA CCCCGTAACA AAAAGGAAGC ACAAAAATTT ATTGGATTAT
AATAATGGAA AACAAGAAAT TTTGCCTAAA GCTAGACAAA AGATATTAGA TTTTGCAGCA CCCCGTAACA AAAAGGAAGC ACAAAAATTT ATTGGATTAT
AATAATGGAA AACAAGAAAT TTTGCCTAAA GCTAGACAAA AGATATTAGA TTTTGCAGCA CCCCGTAACA AAAAGGAAGC ACAAAAATTT ATTGGATTAT
357 N N G K Q E I L P K A R Q K I L D F A A P R N K K E A Q K F I G L F

3501 TCGGATTTTG GAGGGCACAT ATTCCACATT TAAGTCGGCT CCTTGCACCT TTATATAAAG TTACCAGGAA AAAATATGAA TTTGAATGGG GAATACAACA
TCGGATTTTG GAGGGCACAT ATTCCACATT TAAGTCGGCT CCTTGCACCT TTATATAAAG TTACCAGGAA AAAATATGAA TTTGAATGGG GAATACAACA
TCGGATTTTG GAGGGCACAT ATTCCACATT TAAGTCGGCT CCTTGCACCT TTATATAAAG TTACCAGGAA AAAATATGAA TTTGAATGGG GAATACAACA
391 G F W R A H I P H L S R L L A P L Y K V T R K K Y E F E W G I Q Q

3601 AAAAGAAGCT TTTGAAGCAG CAAAGCATGC TATTCAGACT GCTTTAGATC TATGGCCAAT TAAACCGGGA CCCATTGAGT TACAGGTAGA TGTCATTGAT
AAAAGAAGCT TTTGAAGCAG CAAAGCATGC TATTCAGACT GCTTTAGATC TATGGCCAAT TAAACCGGGA CCCATTGAGT TACAGGTAGA TGTCATTGAT
AAAAGAAGCT TTTGAAGCAG CAAAGCATGC TATTCAGACT GCTTTAGATC TATGGCCAAT TAAACCGGGA CCCATTGAGT TACAGGTAGA TGTCATTGAT
424 K E A F E A A K H A I Q T A L D L W P I K P G P I E L Q V D V I D

3701 CAGTATGCCA GCTGGAGTTT ATGGCAAAAA CAAGGTGCCC GACGTTATCC TCTGGGATTT TGGAGTCGCA AGCTTCCCTG TTCCAGTGAA AGATACAGTC
CAGTATGCCA GCTGGAGTTT ATGGCAAAAA CAAGGTGCCC GACGTTATCC TCTGGGATTT TGGAGTCGCA AGCTTCCCTG TTCCAGTGAA AGATATAGTC
CAGTATGCCA GCTGGAGTTT ATGGCAAAAA CAAGGTGCCC GACGTTATCC TCTGGGATTT TGGAGTCGCA AGCTTCCCTG TTCCAGTGAA AGATATAGTC
457 Q Y A S W S L W Q K Q G A R R Y P L G F W S R K L P C S S E R Y S P

3801 CTTTTGAAAA ACAGTTGTTA GCCTGCTATT GGGCTCTAAT AGACTCAGAA AGCCTTACCC TGGGACATGA GGTAATTATG CGACCTCAGA TACCCATCAT
CTTTTGAAAA ACAGTTGTTA GCCTGCTATT GGGCTCTAAT AGACTCAGAA AGCCTTACCC TGGGACATGA GGTAATTATG CGACCTCAGA TACCCATCAT
CTTTTGAAAA ACAGTTGTTA GCCTGCTATT GGGCTCTAAT AGACTCAGAA AGCCTTACCC TGGGACATGA GGTAATTATG CGACCTCAGA TACCCATCAT
491 F E K Q L L A C Y W A L I D S E S L T L G H E V I M R P Q I P I M

3901 GCAATGGATT CATAGCACGC CTGTCACTCA TAAAATTGGA CATGCACAAG AATTTAATAT CATCAAATGG AAATGGTATA TTCAAGACCG GGCCAAACCT
GCAATGGATT CATAGCACGC CTGTCACTCA TAAAATTGGA CATGCACAAG AATTTAATAT CATCAAATGG AAATGGTATA TTCAAGACCG GGCCAAACCT
GCAATGGATT CATAGCACGC CTGTCACTCA TAAAATTGGA CATGCACAAG AATTTAATAT CATCAAATGG AAATGGTATA TTCAAGACCG GGCCAAACCT
525 Q W I H S T P V T H K I G H A Q E F N I I K W K W Y I Q D R A K P

4001 GGCCCAAAGG GGTTGTCCCT ATTGCATGAA CAAGTGTCTC AATATAGCAG TGAGCCAACT GTGTCACCCA AGACTGATAT AACAGAATCT CCTGTTAAAT
GGCCCAAAGG GGTTGTCCCT ATTGCATGAA CAAGTGTCTC AATATAGCAG TGAGCCAACT GTGTCACCCA AGACTGATAT AACAGAATCT CCTGTTAAAT
GGCCCAAAGG GGTTGTCCCT ATTGCATGAA CAAGTGTCTC AATATAGCAG TGAGCCAACT GTGTCACCCA AGACTGATAT AACAGAATCT CCTGTTAAAT
557 G P K G L S L L H E Q V S Q Y S S E P T V S P K T D I T E S P V K W

4101 GGGGTGTCTC ATATGATCAA TTGACTGAAG AACAACAAAA ACATACCTGG TTTACGGATG GTTCTGCCAA AATGATATCG AGCTCACGGA AATGGAAAGC
GGGGTGTCTC ATATGATCAA TTGACTGAAG AACAACAAAA ACATACCTGG TTTACGGATG GTTCTGCCAA AACGATATCG AGCTCACGGA AATGGAAAGC
GGGGTGTCTC ATATGATCAA TTGACTGAAG AACAACAAAA ACATACCTGG TTTACGGATG GTTCTGCCAA AACGATATCG AGCTCACGGA AATGGAAAGC
591 G V S Y D Q L T E E Q Q K H T W F T D G S A K M I S S S R K W K A

4201 AGTTGCCTAC AACCCATCCA CCCAACAAAC AATTGTCACC ACTGGAGATA ATATGAGTAG CCAATATGCT GAACTTTATG CGGTGTATCA GGCACTTCAA
AGTTGCCTAC AACCCATCCA CCCAACAAAC AATTGTCACC ACTGGAGATA ATATGAGTAG CCAATATGCT GAACTTTATG CGGTGTATCA GGCACTTCAA
AGTTGCCTAC AACCCATCCA CCCAACAAAC AATTGTCACC ACTGGAGATA ATATGAGTAG CCAATATGCT GAACTTTATG CGGTGTATCA GGCACTTCAA
624 V A Y N P S T Q Q T I V T T G D N M S S Q Y A E L Y A V Y Q A L Q

4301 CAAGAACAGG GGCAACAATG CCATATATAT ACTGATTCAT GGGCAGTGGC ACAAGGTTTG GCAACGTGGA TGCCACAATG GAAAAAACAT GATTGGAAAA
CAAGAACAGG GGCAACAATG TCATATATAT ACTGATTCAT GGGCAGTGGC ACAAGGTTTG GCAACGTGGA TGCCACAATG GAAAAAACAT GATTGGAAAA
CAAGAACAGG GGCAACAATG TCATATATAT ACTGATTCAT GGGCAGTGGC ACAAGGTTTG GCAACGTGGA TGCCACAATG GAAAAAACAT GATTGGAAAA
657 Q E Q G Q Q C H I Y T D S W A V A Q G L A T W M P Q W K K H D W K I

4401 TTAATGATAA AGAAATATGG GGAAAGAAAT TATGGGAAGA CATATGGTTA TGGTGTCAAA ACACAATTGT TACTGTATTT CATGTTGATG CTCACAGTTC
TTAATGATAA AGAAATATGG GGAAAGAAAT TATGGGAAGA CATATGGTTA TGGTGTCAAA ACACAATTGT TACTGTATTT CATGTTGATG CTCACAGTTC
TTAATGATAA AGAAATATGG GGAAAGAAAT TATGGGAAGA CATATGGTTA TGGTGTCAAA ACACAATTGT TACTGTATTT CATGTTGATG CTCACAGTTC
691 N D K E I W G K K L W E D I W L W C Q N T I V T V F H V D A H S S

```
4501 TTTAATTTCT GCAGAACGAA AGCATAATTC TCATGCTAAT CAGCTGGTGC AGATTCGTGC AACGCACAAC CCAGGCTTAG CACAATGGGT CCACGAAAAG
     TTTAATTTCT GCAGAACGAA AGCATAATTC TCATGCTAAT CAGCTGGTGC AGATTCGTGC AACGCACAAC CCAGGCTTAG CACAATGGGT CCACGAAAAG
     TTTAATTTCT GCAGAACGAA AGCATAATTC TCATGCTAAT CAGCTGGTGC AGATTCGTGC AACGCACAAC CCAGGCTTAG CACAATGGGT CCACGAAAAG
 724  L  I  S   A  E  R  K   H  N  S   H  A  N   Q  L  V  Q   I  R  A   T  H  N   P  G  L  A   Q  W  V   H  E  K

4601 AGTGGACACT TGGGAGAACA AGCTTCATAC AGGTGGGCAC AACAAAGGGG AATCCTTGTC ATCCATGATG ATATTGCCAC TGCAGTACAA CAATGTCAGT
     AGTGGACACT TGGGAGAACA AGCTTCATAC AGGTGGGCAC AACAAAGGGG AATCCTTGTC ACCCATGATG ATATTGCCAC TGCAGTACAA CAATGTCAGT
     AGTGGACACT TGGGAGAACA AGCTTCATAC AGGTGGGCAC AACAAAGGGG AATCCTTGTC ACCCATGATG ATATTGCCAC TGCAGTACAA CAATGTCAGT
 757 S  G  H  L   G  E  Q   A  S  Y   R  W  A  Q   Q  R  G   I  L  V   I  H  D  D   I  A  T   A  V  Q   Q  C  Q  L

4701 TATGCCAACA GTTAAATAAA CGGGGAGTTC CACACCCCTA CCAAGGTCAT ATCCAAAAGG GATTATTTCC TGCCCATACC TGGCAAATAG ATTTTATTGG
     TATGCCAACA GTTAAATAAA CGGGGAGTTC CACACCCCTA CCAAGGTCAT ATCCAAAAGG GATTATTTCC CGCCCATACA TGGCAAATAG ATTTTATTGG
     TATGCCAACA GTTAAATAAA CGGGGAGTTC CACACCCCTA CCAAGGTCAT ATCCAAAAGG GATTATTTCC CGCTCATACA TGGCAAATAG ATTTTATTGG
 791  C  Q  Q   L  N  K   R  G  V  P   H  P  Y   Q  G  H   I  Q  K  G   L  F  P   A  H  T   W  Q  I  D   F  I  G

4801 ACCATTGCCA AATTCTTGTG GATATACTTA TGCCTGCACT GCTGTTGATA CATATTCTGG TTATTTATTG GCTATACCAG CTAAAGCGGC CACCCAACAG
     ACCATTGCCA AATTCTTGTG GATATACTTA TGCCTGCACT GCTGTTGATA CATATTCTGG TTATTTATTG GCTATACCAG CTAAAGCGGC CACCCAACAG
     ACCATTGCCA AATTCTTGTG GATATACTTA TGCCTGCACT GCTGTTGATA CATATTCTGG TTATTTATTG GCTATACCAG CTAAAGCGGC CACCCAACAG
 824  P  L  P   N  S  C  G   Y  T  Y   A  C  T   A  V  D  T   Y  S  G   Y  L  L   A  I  P  A   K  A  A   T  Q  Q

4901 AGTGCCATAA AATTATTAGA TACAATCAAG TTATATTATG GAACACCAAG GCAAATTCAG AGTGACAACG GTTCCCACTT TACAGGTAAG TTGATTCAAA
     AGTGCCATAA AATTATTAGA TACAATCAAG TTATATTATG GAACACCAAG GCAAATTCAG AGTGACAACG GTTCCCACTT TACAGGTAAG TTGATTCAAA
     AGTGCCATAA AATTATTAGA TACAATCAAG TTATATTATG GAACACCAAG GCAAATTCAG AGTGACAACG GTTCCCACTT TACAGGTAAG TTGATTCAAA
 857 S  A  I  K   L  L  D   T  I  K   L  Y  Y  G   T  P  R   Q  I  Q   S  D  N  G   S  H  F   T  G  K   L  I  Q  T

5001 CCTATACCAA GGAAAATTAT ATAGAATGGA TTTATCATAT ACCTTATTAC CCCCAAGCTG CAGGCCTCAT AGAACGAATG AACGGATTGC TTAAACAACA
     CCTATACCAA GGAAAATTAT ATAGAATGGA TTTATCATAT ACCTTATTAC CCCCAAGCTG CAGGCCTCAT AGAACGAATG AACGGATTGC TTAAACAACA
     CCTATACCAA GGAAAATTAT ATAGAATGGA TTTATCATAT ACCTTATTAC CCCCAAGCTG CAGGCCTCAT AGAACGAATG AACGGATTGC TTAAACAACA
 891  Y  T  K   E  N  Y   I  E  W  I   Y  H  I   P  Y  Y   P  Q  A  A   G  L  I   E  R  M   N  G  L  L   K  Q  Q

5101 ATTGAGGAAA CTTGGTCATG GCTCATTAAA GGGGTGGCAC AACCATCTAA GTACTGCTCT AAATGTGTTA AATAACAGGC CACTAGGCCC TAATGAGACT
     ATTGAGGAAA CTTGGTCATG GCTCATTAAA GGGGTGGCAC AACCATCTAA GTACTGCTCT AAATGTGTTA AATAACAGGC CACTAGGCCC TAATGAGACT
     ATTGAGGAAA CTTGGTCATG GCTCATTAAA GGGGTGGCAC AACCATCTAA GTACTGCTCT AAATGTGTTA AATAACAGGC CACTAGGCCC TAATGAGACT
 924  L  R  K   L  G  H  G   S  L  K   G  W  H   N  H  L  S   T  A  L   N  V  L   N  N  R  P   L  G  P   N  E  T

5201 CCTCTGTCAA GGTTGTTACC AGCAAAGATT ACAGAAGTGG CAACCGCCAC CCATGAGTTT ATGACCTTAA CCTATTGGCC CCTGACTCAG TCTGCTAAAC
     CCTCTGTCAA GGTTGTTACC AGCAAAGATT ACAGAAGTGG CAACCGCCAC CCATGAGTTT ATGACCTTAA CCTATTGGCC CCTGACTCAG TCTGCTAAAC
     CCTCTGTCAA GGTTGTTACC AGCAAAGATT ACAGAAGTGG CAACCGCCAC CCATGAGTTT ATGACCTTAA CCTATTGGCC CCTGACTCAG TCTGCTAAAC
 957 P  L  S  R   L  L  P   A  K  I   T  E  V  A   T  A  T   H  E  F   M  T  L  T   Y  W  P   L  T  Q   S  A  K  P

5301 CGCCTTTTCG TGCCACGGCC GAAGCGGCTG GATTGGATTT ATCTACTGTT CATTCTATTC AATTGCCCCC TAAGAAGCAG TCTATAGTCC CTACTGGGAT
     CGCCTTTTCG TGCCACGGCC GAAGCGGCTG GATTGGATTT ATCTACTGTT CATTCTATTC AATTGCTCCC TAGGAAGCAG TCTATAGTCC CTACTGGGAT
     CGCCTTTTCG TGCCACGGCC GAAGCGGCTG GATTGGATTT ATCTACTGTT CATTCTATTC AATTGCTCCC TAGGAAGCAG TCTATAGTCC CTACTGGGAT
 991  P  F  R   A  T  A   E  A  A  G   L  D  L   S  T  V   H  S  I  Q   L  P  P   K  K  Q   S  I  V  P   T  G  I

5401 AGGGGTACAA TTTCCTAAAA ATACCTTTGG TTTGTTAAAA GGACGCTCTG GACTGGCAGC AAAAGGTATT GATGTACTGG GAGGAGTTAT TGATCCAGAT
     AGGGGTACAA TTTCCCAAAA ATACCTTTGG TTTGTTAAAA GGACTCTCTG GACTGGCAGC AAAAGGTATT GATGTACTGG GAGGAGTTAT TGATCCAGAT
     AGGGGTACAA TTTCCCAAAA ATACCTTTGG TTTGTTAAAA GGACGCTCTG GACTGGCAGC AAAAGGTATT GATGTACTGG GAGGAGTTAT TGATCCAGAT
1024  G  V  Q   F  P  K  N   T  F  G   L  L  K   G  R  S  G   L  A  A   K  G  I   D  V  L  G   G  V  I   D  P  D

5501 TACCAGGGAG AAATAAAATT TATATTATAT AATGGTAATG ACACAGTACT AACTGTAGAG TCTGGGGAAC GAGTGGGACA ATTATTGGTG CTTCCCCTCT
     TACCAGGGAG AAATAAAATT TATATTATAT AATGGTAGTG ACACAGTACT AACTGTAGAG TCTGGGGAAC GAGTGGGACA ATTATTGGTA CTTCCCCTCT
     TACCAGGGAG AAATAAAATT TATATTATAT AATGGTAGTG ACACAGTACT AACTGTAGAG TCTGGGGAAC GAGTGGGACA ATTATTGGTA CTTCCCCTCT
1057 Y  Q  G  E   I  K  F   I  L  Y   N  G  N  D   T  V  L   T  V  E   S  G  E  R   V  G  Q   L  L  V   L  P  L  L

5601 TGACTCCAAA TGTGTCACAA GGACAACCTC CAAGTGAATT AACTAAAAGA GGAATACAAG GATTTGGATC TACAGATGGA TGGAAACCTG GAGCAAAAGT
     TGACTCCAAA TGTGTCACAA GGACAACCTC CAAGTGAATT AACTAAAAGA GGAATACAAG GATTTGGATC TACAGATGGA TGGAAACCTG GAGCAAAAGT
     TGACTCCAAA TGTGTCACAA GGACAACCTC CAAGTGAATT AACTAAAAGA GGAATACAAG GATTTGGATC TACAGATGAA TGGAAACCTG GAGCAAAAGT
1091  T  P  N   V  S  Q   G  Q  P  P   S  E  L   T  K  R   G  I  Q  G   F  G  S   T  D  G   W  K  P  G   A  K  V

5701 GTGGGTGACA CATCCACCCA ATGCGGCTCC GCGTGCCGCA GAAGTAGTGG CCCAAGGGCC CACTGCAACA CTCATAGTAA TGTATCCAGG AAATGAACAA
     GTGGGTGACA CATCCACCCA ATGCGGCTCC CCGTGCCGCA GAAGTAGTGG CCCAAGGGCC CACTGCAACA CTCATAGTAA TGTATCCAGG AAATGAACAA
     GTGGGTGACA CATCCACCCA ATGCGGCTCC CCGTGCCGCA GAAGTAGTGG CCCAAGGGCC CACTGCAACA CTCATAGTAA TGTATCCAGG AAATGAACAA
1124  W  V  T   H  P  P  N   A  A  P   R  A  A   E  V  V  A   Q  G  P   T  A  T   L  I  V  M   Y  P  G   N  E  Q

5801 TTATATCATG TTCCTAAAAA TTCTGTTTCT TTGCGTGAAT AGATATTCCT GCTGTGTTTA CTGTGCTGTG CTGCACTTCC ATGTCTGCTG CAGACCTACA
     TTATATCATG TTCCTAAAAA TTCTGTTTCT TTGCGTGAAT AGATATTCCT GCTGTGTTTA CTATGCTGTG CTGCACTTCC ATGTCTGCTG CAGACCTACA
     TTATATCATG TTCCTAAAAA TTCTGTTTCT TTGCGTGAAT AGATATTCCT GCTGTGTTTA CTATGCTGTG CTGCACTTCC ATGTCTGCTG CAGACCTACA
1157 L  Y  H  V   P  K  N   S  V  S   L  R  E  *   I  F  L   L  C  L   L  C  C  A   A  L  P   C  L  L   Q  T  Y  T
                              Splice acceptor (presumed)^ ^env open reading frame                   ^SU

5901 CGCCAACTAA TGTCTCCGTA GGACAACCTT TGACCCTGCA CTGCTGGACC AACTGCTCTA CACCAGTGGG ACCTATAACC TGGACTATGA ATGGCCGAGA
     CGCCAACTAA TGTCTCCGTA GGACAACCTT TGACCCTGCA CTGCTGGACC AACTGTTCTA CACCAGTGGG ACCTATAACC TGGACTATGA ATGGCCGGGA
     CGCCAACTAA TGTCTCCGTA GGACAACCTT TGACCCTGCA CTGCTGGACC AACTGTTCTA CACCAGTGGG ACCTATAACC TGGACTATGA ATGGCCGGGA
  21  P  T  N   V  S  V   G  Q  P  L   T  L  H   C  W  T   N  C  S  T   P  V  G   P  I  T   W  T  M  N   G  R  E
```

```
6001 AATTTGGTCT CAGAAACATC AAGCAATCTC TCGGTACAAG CCTATATCTG ACCAGTCACG CAGAACTAGT TATAATTTCT CCATTGCTCT GTCTTCTGTT
     AATTTGGTCT CAGAAACATC AAGCAATCTC TCGGTACAAG CCTATATCTG ACCAGTCACA CAGAACTAGT TATAATTTCT CCATTGCTCT GTCTTCTGTT
     AATTTGGTCT CAGAAACATC AAGCAATCTC TCGGTACAAG CCTATATCTG ACCAGTCACA CAGAACTAGT TATAATTTCT CCATTGCTCT GTCTTCTGTT
  54 I W S  Q K H  Q A I  S R Y K  P I S  D Q S R  R T S  Y N F S  I A L  S S V

6101 ACCCTCCGTG ATGGGGGCTG GTATTCTTGT ATTAAATGGA AGAAGGGAAA TCCAGATTCC CCGTTTCTTA AAGAAACTCG ATTTATTAGC ATCCTTCCTA
     ACCCTCCGTG ATGGGGGCTG GTATTCTTGT ATTAAATGGA AGAAGGGAAA TCCAGATTCC CCGTTTCTTA AAGAAACTCG ATTTATTAGC ATCCTTCCTA
     ACCCTCCGTG ATGGGGGCTG GTATTCTTGT ATTAAATGGA AGAAGGGAAA TCCAGATTCC CCGTTTCTTA AAGAAACTCG ATTTATTAGC ATCCTTCCTA
  87 T L R D  G G W  Y S C  I K W K  K G N  P D S  P F L K  E T R  F I S  I L P N

6201 ATGCCACTAC TCCTCTTAAA AACACTAATC TGTGGTTACA TTTAGCTCAT ACAGTTCTTA ATGAAACTGC TTTTTGTCTC AGAAACAGTA ATAATCCTTC
     ATGCCACTAC TCCTCTTAAA AACACTAATC TGTGGTTACA TTTAGCTCAT ACAGTTCTTA ATGAAACTGC TTTTTGTCTC AGAAACAGTA ATAATCCTTC
     ATGCCACTAC TCCTCTTAAA AACACTAATC TGTGGTTACA TTTAGCTCAT ACAGTTCTTA ATGAAACTGC TTTTTGTCTC AGAAACAGTA ATAATCCTTC
 121  A T T  P L K  N T N L  W L H  L A H  T V L N  E T A  F C L  R N S N  N P S

6301 TGGACTTCTG CAAGCTAATC TGTTGGGTCT AGGTGTTACT TTACCTAATG TGTTACATGC TAAAGTTACT ATTCCTCCTT CTGTAGTAAC GATGCCCATA
     TGGACTTCTG CAAGCTAATC TGTTGGGTCT AGGTGTTACT TTACCTAATG TGTTACATGC TAAAGTTACT ATTCCTCCTT CTTTAGTAAC GATGCCCATA
     TGGACTTCTG CAAGCTAATC TGTTGGGTCT AGGTGTTACT TTACCTAATG TGTTACATGC TAAAGTTACT ATTCCTCCTT CTTTAGTAAC GATGCCCATA
 154 G L L  Q A N L  L G L  G V T  L P N V  L H A  K V T  I P P S  V V T  M P I

6401 TCAGGAGTGA CCTCCACTAC ATGTTACTGT ATTAATTGTA CATATATTCC ATTCTATTGT AATAAGACTA TCCCTGTTAG TATTCCTACC CCCTCACGAT
     TCAGGAGTGA CTTCCACTAC ATGTTACTGT ATTAATTGTA CATATATTCC ATTCTATTGT AATAAGACTA TCCCTGTTAG TATTCCTACC CCCTCACGAT
     TCAGGAGTGA CTTCCACTAC ATGTTACTGT ATTAATTGTA CATATATTCC ATTCTATTGT AATAAGACTA TCCCTGTTAG TATTCCTACC CCCTCACGAT
 187 S G V T  S T T  C Y C  I N C T  Y I P  F Y C  N K T I  P V S  I P T  P S R L

6501 TACCTGCTGG ATATTTCTGG TTATGTGATA ATGCTGCGTA TTCACACATT CCAACTGACA CATCTGTTGC ATGTGGTATT GGTCCAATAG TTCCCCTTAT
     TACCTGCTGG ATATTTCTGG TTATGTGATA ATGCTGCGTA TTCACACATT CCAACTGACA CATCTGTTGC ATGTGGTATT GGTCCAATAG TTCCCCTTAT
     TACCTGCTGG ATATTTCTGG TTATGTGATA ATGCTGCGTA TTCACACATT CCAACTGACA CATCTGTTGC ATGTGGTATT GGTCCAATAG TTCCCCTTAT
 221  P A G  Y F W  L C D N  A A Y  S H I  P T D T  S V A  C G I  G P I V  P L I

6601 AGCCGCTTTC CCTACCAAGA ATTCTCGAGT GAAACGAGCA CTCACTGCAG ATTGTGACCC TCATGTTAAC CTTCTTCCCA TCTCTACTGT GGCAACGCTG
     AGCCGCTTTC CCTACCAAGA ATTCTCGAGT GAAACGAGCA CTCACTGCAG ATTGTGACCC TCATGTTAAC CTTCTTCCCA TCTCTACTGT GGCAACGCTG
     AGCCGCTTTC CCTACCAAGA ATTCTCGAGT GAAACGAGCA CTCACTGCAG ATTGTGACCC TCATGTTAAC CTTCTTCCCA TCTCTACTGT GGCAACGCTG
 254  A A F  P T K N  S R V  K R A  L T A D  C D P  H V N  L L P I  S T V  A T L
                                ^TM

6701 GCCGCCCTTC TACTACCTGA TCCTGGTCTA ATTGTAGACA ACCATCGTCA AATTGAACAC CTTTCCTGCT TGTTGACACA TGTAATGAAT GATACTGTTA
     GCCGCCCTTC TACTACCTGA TCCTGGTCTA ATTGTAGACA ACCATCGTCA AATTGAACAC CTTTCCTGCT TGTTGACACA TGTAATGAAT GATACTGTTA
     GCCGCCCTTC TACTACCTGA TCCTGGTCTA ATTGTAGACA ACCATCGTCA AATTGAACAC CTTTCCTGCT TGTTGACACA TGTAATGAAT GATACTGTTA
 287 A A L L  P D P  G L I  V D N  H R Q  I E H  L S C L  L T H  V M N  D T V R

6801 GAGCCTTCTT GGCTATATCT CAAGAATTAC AAGAACTGCG TAGAGAGACT CTTATACAAC GAGCTGCTTT AGACTATTTG TTTCTTCTCC ATCAAACACC
     GAGCCTTCTT GGCTATATCT CAAGAATTAC AAGAACTGCG TAGAGAGACT CTTATACAAC GAGCTGCTTT AGACTATTTG TTTCTTCTCC ATCAAACACC
     GAGCCTTCTT GGCTATATCT CAAGAATTAC AAGAACTGCG TAGAGAGACT CTTATACAAC GAGCTGCTTT AGACTATTTG TTTCTTCTCC ATCAAACACC
 321  A F L  A I S  Q E L Q  E L R  R E T  L I Q R  A A L  D Y L  F L L H  Q T P

6901 TTGTACTAAA ATACCGGGTA TGTGTTGTTT CAATGTATCA GATCAAACTA ATGTTATCTT TAATGCTGTA CATGATTTGC AAGATCAAGT TAATAACATT
     TTGTACTAAA ATACCGGGTA TGTGTTGTTT CAATGTATCA GATCAAACTA ATGTTATCTT TAATGCTGTA CATGATTTGC AAGATCAAGT TAATAACATT
     TTGTACTAAA ATACCGGGTA TGTGTTGTTT CAATGTATCA GATCAAACTA ATGTTATCTT TAATGCTGTA CATGATTTGC AAGATCAAGT TAATAACATT
 354  C T K  I P G M  C C F  N V S  D Q T N  V I F  N A V  H D L Q  D Q V  N N I

7001 TCATATTCTA AGGGGTTATT TGATTGGTTA CCTTGGGGTT TGGGTCCTTT ATTGCGAGAT AGTTTTATAA TTATTGTAGT CATTGGATTT TGGATTATTG
     TCATATTCTA AGGGGTTATT TGATTGGTTA CCTTGGGGTT TGGGTCCTTT ATTGCGAGAT AGTTTTATAA TTATTGTAGT CATTGGATTT TGGATTATTG
     TCATATTCTA AGGGGTTATT TGATTGGTTA CCTTGGGGTT TGGGTCCTTT ATTGCGAGAT AGTTTTATAA TTATTGTAGT CATTGGATTT TGGATTATTG
 387 S Y S K  G L F  D W L  P W G L  G P L  L R D  S F I I  I V V  I G F  W I I G

7101 GATGTGCCTT ATGTACTTGT GTACGAAGTT CATTGTCTGT GCGTATGGTG TCTTATGGTT AATGGAGACG GCCACGGCCG AATATGTCTT GACATGAGTG
     GATGTGCCTT ATGTACTTGT GTACGAAGTT CATTGTCTGT GCGTATGGTG TCTTATGGTT AATGGAGACG GCCACGGCCG AATATGTCTT GACATGAGTG
     GATGTGCCTT ATGTACTTGT GTACGAAGTT CATTGTCTGT GCGTATGGTG TCTTATGGTT AATGGAGACG GCCACGGCCG AATATGTCTT GACATGAGTG
 421  C A L  C T C  V R S S  L S V  R M V  S Y G *      Polypurine tract    ^LTR

7201 CAGCCATGTT TGGCCGCTCC ATTGACCTAC AGCCACCAGC ACAAAAAGAA ATATAAAAGC TGCTTTCTGC GTGGTGGGAA CTGGCCCTTC TCTCACGGAG
     CAGCCATGTT TGGCCGCTCC GTTGACCTAC AGCCACCAGC ACAAAAAGAA ATATAAAAGC TGCTTTCTGC GTGGTGGGAA CTGGCCCTTC TCTCACGGAG
     CAGCCATGTT TGGCCGCTCC GTTGACCTAC AGCCACCAGC ACAAAAAGAA ATATAGAAGC TGCTTTCTGC GTGGTGGGAA CTGGCCCTTC TCTCACGGAG

7301 GGAGTTGCAG GTTGTAACAT TTCAAGGCTC TTGGAGCGTC GTAGCACGGG ATGGACTGGC CATCTATGCC GGGCTGGGAC GATAACCAAA GAGAACAATG
     GGAGTTGCAG GTTGTAACAT TTCAAGGCTC TTGGAGCGTC GTAGCACGGG ATGGACTGGC CATCTATGCC GGGCTGGGAC GATAACCAAA GAGAACAATG
     GGAGTTGCAG GTTGTAACAT TTCAAGGCTC TTGGAGCGTC GTAGCACGGG ATGGACTGGC CATCTATGCC GGGCTGGGAC GATAACCAAA GAGAACAATG

7401 CACGTACACA ACTACTGGGC TGGGACGTTA GCCAGGGGGC TCAGTGCACG TACGCCACTG ATAACCATTT TGTGCATGGG AACACTAAAT GCTTGCTCTG
     CACGTACACA ACTACTGGGC TGGGACGTTA GCCAGGGGGC TCAGTGCACG TACGCCACTG ATAACCATTT TGTGCATGGG AACACTAAAT GCTTGCTCTG
     CACGTACACA ACTACTGGGC TGGGACGTTA GCCAGGCGGC TCAGTGCACG TACGCCACTG ATAACCATTT TGTGCATGGG AACACTGAAT GCTTGCTCTG

7501 CAAACTCTGT AATTGCTTAC TCTGAAAATT ACTGATGGTA TAAAAACTGC TGGAAACTGA GACCGAGAGA GAGTTCCACC TGTGGAAGGG ACACCTCGCC
     CAAACTCTGT AATTGCTTAC TCTGAAAATT ACTGATGGTA TAAAAACTGC TGGAAACTGA GACCCAGAGA GAGTTCCACC TGTGGAAGGG ACACCTCGCC
     CAAACTCTGT AATTGCTTAC TCTGAAAATT ACTGATGGTA TAAAAACTGC TGGAAACTGA GACCCAGAGA GAGTTCCACC TGTGGAAGGG ACACCTCGCC
```

```
7601 CAGGACGTGT GATCCTTGCC CAGCCGCGTC GATTGACAGG GCTCTCCTGG CAGTGGGACG TGGATGTTGT GAGTAACTGA TTTGATGTGA AGCAATTGAT
     CAGGACGTGT GATCCTTGCC CAGCCGCGTC AATTGACAGG GCTCTCCTGG CAGTGGGACG TGGATGTTGT GAGTAACTGA TTTAATGTGA AGTAATTGAT
     CAGGACGTGT GATCCTTGCC CAGCCGCGTC AATTGACAGG GCTCTCCTGG CAGTGGGACG TGGATGTTGT GAGTAACTGA TTTAATGTGA AGTAATTGAT

7701 TTGATATGAA GTAATTGATT TGATGTGTTC TCTTTTTGGT CAACCTGGTG TTTCTCTTTC TGGTTGATTT GTGTCATTTC TCTTCAGCCG ATGTGGGTGC
     TTGATGTGTA GTAAGTGATT TGATGTGTTC TTTTTTTGGT CAACCTGGTG TTTCTCTTTC TGGTTGATTT GTGTCATTTC TCTTCAGCCG ATGTGGGTGC
     TTGATGTGTA GTAAGTGATT TGATGTGTTC TTTTTTTGGT CAACCTGGTG TTTCTCTTTC TGGTTGATTT GTGTCATTTC TCTTCAGCCG ATGTGGGTGC

7801 TTTCCCTTTC CAGTCGGGCT GATGTTTTTT CCCTCTTAAT ATTTGACCTA TTTGGATCTG TTTGCAGACT GTCGCCTTTA CTATCCCCTA TATAATAAAA
     TTTCCCTTTC CAGTCGGGCT GATGTTTTTT CCCTCTTAAT ATTTGACCTA TTTGGATCTG TTTGCAGACT GTCGCCTTTA CTATCCTCTA TATAATAAAA
     TTTCCCTTTC CAGTCGGGCT GATGTTTTTT CCCTCTTAAT ATTTGACCTA TTTGGATCTG TTTGCAGATT GTCGCCTTTA CTATCCTCTA TATAATAAAA
                                                                                                      Poly(A)
7901 TATACCTTCA GTCCATTTGT TTGGAGTGGA AAGTGTCTTT TACGTCTCCG ATCGAATCCC CGAACCTCTC GTAACA
     TATACCTTCA GTCCATCTGT TTGGAGTGGA AAGTGTCTTT TACGTCTCCG ATCGAATCCC CGAACCTCTG ATAACA
     TATACCTTCA GTCCATCTGT TTGGAGTGGA AAGTGTCTTT TACGTCTCCG ATCGAATCCC CGAACCTCTG ATAACA
                                                                              *      ^End LTR
```

**Supplementary Figure 10** – ERV-S.1-Dbi proviral sequence aligned to related proviruses in the *D. bicornis minor* and *D. bicornis michaelis* genomes. The first nucleotide sequence in each row is from ERV-S.1-Dbi in GenBank accession JAJIAZ010000042.1:6,306,454-6,314,429 (reverse-complement strand). The second nucleotide sequence in each row is from a *D. bicornis minor* provirus in JAJIAY010000041.1:52,088,928-52,096,904 (reverse complement strand). The third nucleotide sequence is from a *D. bicornis michaeli* provirus in JANTPW010000004.1:23,894,873-23,902,845 (reverse-complement strand) in the same integration site and orientation as the *D. bicornis minor* JAJIAY010000041.1 provirus. The sequences flanking both proviruses are shown before the provirus alignment in the same order as the nucleotide alignment, with the 5-bp direct repeats underlined and the identical flanking sequences in the second and third proviruses in different subspecies shown in blue. Nucleotide substitutions in the JAJIAY010000041.1 and JANTPW010000004.1 proviruses relative to ERV-S.1-Dbi are highlighted in black background and insertions and deletions in red backgrounds. Major sequence features are underlined or indicated with an arrowhead and labeled. The only position differing between the LTR sequences in the JAJIAY010000041.1 sequence is indicated with an asterisk. Amino acid sequences are translations from the ERV-S.1-Dbi provirus (top) nucleotide sequence. Only the numbering for the ERV-S.1-Dbi provirus (top nucleotide sequence in each row and amino acid sequence) is shown.
